# Dynamic mapping of proteome trafficking within and between living cells by TransitID

**DOI:** 10.1101/2023.02.07.527548

**Authors:** Wei Qin Xu, Joleen S. Cheah, Charles Xu, James Messing, Brian D. Freibaum, Steven Boeynaems, J. Paul Taylor, Namrata D. Udeshi, Steven A. Carr, Alice Y. Ting

**Author notes:** The authors contributed equally to this work (Wei Qin Xu, Joleen S. Cheah).

## Abstract

The ability to map trafficking for thousands of endogenous proteins at once in living cells would reveal biology currently invisible to both microscopy and mass spectrometry. Here we report TransitID, a method for unbiased mapping of endogenous proteome trafficking with nanometer spatial resolution in living cells. Two proximity labeling (PL) enzymes, TurboID and APEX, are targeted to source and destination compartments, and PL with each enzyme is performed in tandem via sequential addition of their small-molecule substrates. Mass spectrometry identifies the proteins tagged by both enzymes. Using TransitID, we mapped proteome trafficking between cytosol and mitochondria, cytosol and nucleus, and nucleolus and stress granules, uncovering a role for stress granules in protecting the transcription factor JUN from oxidative stress. TransitID also identifies proteins that signal intercellularly between macrophages and cancer cells. TransitID introduces a powerful approach for distinguishing protein populations based on compartment or cell type of origin.

## Introduction

Proteins are highly dynamic and often transit through multiple organelles or cells over the course of their lifetimes^1, 2^. Whereas the movement of individual proteins can be studied by fluorescent protein tagging and microscopy, no method exists for the unbiased mapping or discovery of endogenous proteins that transit between specific organelles or cells. Such a capability could shed enormous light on the molecular mechanisms used by organelles and cells to communicate with and regulate one another, how distinct signaling functions are compartmentalized by cells, and the roles of individual signaling molecules.

Here we report TransitID (TRafficking ANalysis by Sequentially Incorporated Tags for IDentification), a general and versatile method for mapping the trafficking and dynamics of endogenous proteomes in living cells. TransitID builds upon proximity labeling (PL)^3^, a method catalyzed by promiscuous enzymes such as APEX2^4^, BioID^5, 6^, and TurboID^7^. PL has been widely applied and shown to be highly effective for generating single-timepoint snapshots of organelle proteomes^8–10^ and protein interactomes^11, 12^ in living samples. However, these datasets reveal little about how proteins *move* in order to signal, provide regulation, and/or respond to extracellular and intracellular cues. To provide this capability, we envisioned multiplexing orthogonal PL enzymes to perform two PL reactions in tandem, within the same biological sample (**Figure 1A**). The first labeling, catalyzed by a PL enzyme targeted to a “source” organelle or cell, would be followed by a chase period ranging from minutes to days. The second labeling would be catalyzed by a second PL enzyme, expressed in a “destination” organelle or cell. Subsequently, the samples would be lysed, and proteins dual-labeled by *both* PL enzymes would be enriched and identified by mass spectrometry (**Figure 1B**). By virtue of their dual labels, these proteins must have originated from the “source” organelle and trafficked to the “destination” organelle over the user-selected chase period.

**Figure 1.**
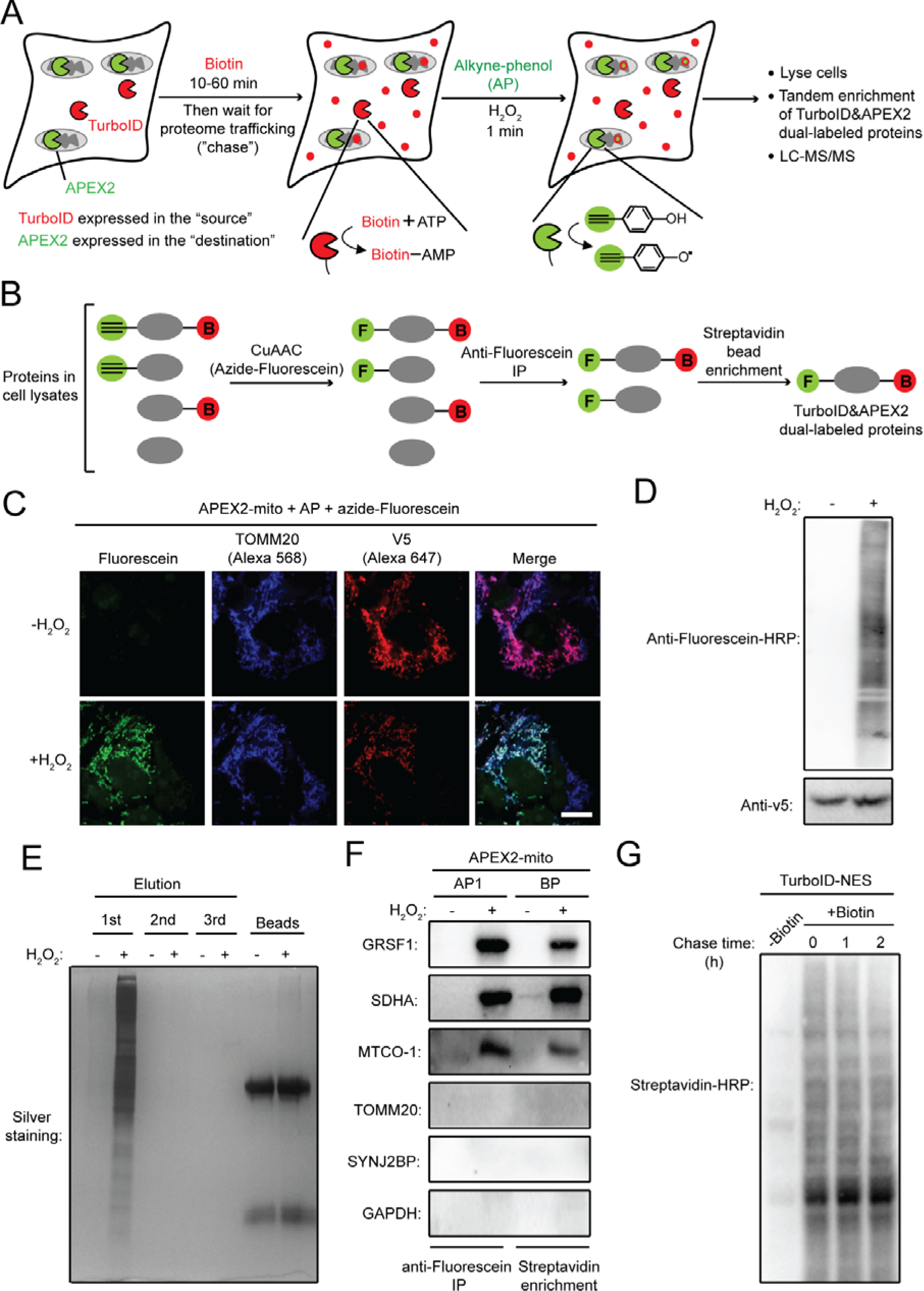
Development of TransitID and characterization of APEX-catalyzed alkyne-phenol labeling. **A.** Scheme showing TransitID in cells with TurboID-catalyzed biotinylation in the “source” location, followed by APEX-catalyzed alkyne tagging in the “destination” location. **B.** Dual enrichment to capture proteins tagged by both TurboID and APEX2. Red B, biotin. Green F, fluorescein. CuAAC, copper-catalyzed azide-alkyne cycloaddition. **C.** Confocal fluorescence imaging of mitochondrial proteins labeled by APEX2 and alkyne-phenol. After cell fixation, click chemistry was performed with fluorescein-azide to visualize alkyne-tagged proteins. TOMM20 is an endogenous mitochondrial marker. V5 is fused to APEX2. Scale bar, 10 μ **D**. Anti-Fluorescein blotting of mitochondrial proteins labeled by APEX2 and alkyne-phenol. Click chemistry was performed on cell lysate with fluorescein-azide. **E**. Anti-fluorescein antibody enrichment of APEX-labeled proteins. All proteins were captured in the first elution from beads. **F**. Blotting for protein markers in APEX-labeled, anti-fluorescein antibody-enriched material. GRSF1, SDHA and MT-CO1 are mitochondrial matrix proteins. Negative controls are TOMM20 and SYNJ2BP (outer mitochondrial membrane) and GAPDH (cytosol). For comparison, the same analysis is performed on enriched material from traditional mito-APEX2 + biotin-phenol labeling and streptavidin enrichment. **G**. Biotin washout stops TurboID labeling. HEK 293T cells expressing TurboID-NES were labeled with biotin for 10 minutes, then washed for 0, 1, or 2 hours before cell lysis and streptavidin blot analysis. No increase in proteome biotinylation extent was observed under the 1- or 2-hour wash conditions.

In this work, we develop this methodology, called TransitID, validate it using nuclear-encoded mitochondrial proteins, and explore three distinct intracellular applications: mapping local versus cytosolic translation of mitochondrial proteins; mapping cytosol-to-nucleus proteome shuttling under stress; and mapping proteome trafficking between the nucleolus and stress granules during the cellular stress response. Specific hits from each dataset are validated and explored, leading to discoveries such as a role for stress granules in protecting the JUN transcription factor from aggregation and degradation following oxidative stress. Finally, we demonstrate that TransitID can be used to capture endogenous proteins secreted by tumor cells and taken up by macrophages, and vice versa, and distinguish between trafficking mechanisms.

### Development of TransitID

The PL enzymes APEX and TurboID use different labeling chemistries and are natural candidates for multiplexing. However, both enzymes use biotinylated substrates, which cannot be distinguished during proteome enrichment. Because APEX binds to its substrate in an open, solvent-exposed cleft, we reasoned that it might be more likely than TurboID to accept alternative, non-biotin substrates. We tested three different alkyne-phenol substrates as alternatives to biotin-phenol (BP), APEX’s original substrate (**Figure S1A**). AP1 showed high labeling efficiency – 5-fold higher than that of biotin-phenol – and minimal cell toxicity (**Figure S1B-D**). The superior labeling efficiency of AP1 may be due to greater membrane permeability, as the labeling efficiency of AP1 and BP are comparable in cell lysates (**Figure S1E**).

The alkyne handle on AP1 enables Click-based derivatization of APEX2-tagged proteins by azide-containing affinity tags. We tested multiple derivatization and enrichment strategies and encountered problems of low recovery with azide-agarose beads (**Figure S1F**) and incomplete removal of azide-FLAG which interfered with protein enrichment (**Figure S1G**). However, we obtained high recovery and low background using Click with azide-fluorescein (FAM), followed by immunoprecipitation (IP) with anti-FAM antibody^13^ (**Figure S1G, H**). We used microscopy (**Figure 1C**) and Western blot detection of on-target and off-target protein markers (**Figure 1D-F**) to verify the spatial specificity of proteome tagging and enrichment by APEX/AP1 and anti-FAM IP.

Using AP1, we tested the orthogonality of TurboID- and APEX2-catalyzed PL reactions. We confirmed that TurboID does not recognize AP1, while APEX2 does not use TurboID’s substrate, biotin (**Figure S1I, J)**. To test both enzymes in the same biological sample, we generated HEK 293T cells expressing TurboID in the cytosol (TurboID-NES) and APEX2 in the mitochondrial matrix (mito-APEX2). We performed TurboID labeling first, due to its non-toxic labeling conditions, and stopped labeling after 1 hour by washing out excess biotin (**Figure 1G**). After a 24 hour “chase”, we initiated APEX labeling in the mitochondrion by adding H_2_O_2_ to the cells for 1 minute (**Figure 2A**). Samples were immediately lysed and analyzed by Western blotting. **Figure 2B** shows promiscuous protein biotinylation in whole cell lysates of samples treated with biotin, and promiscuous FAM-labeling of samples treated with FAM and H_2_O_2_.

**Figure 2:**
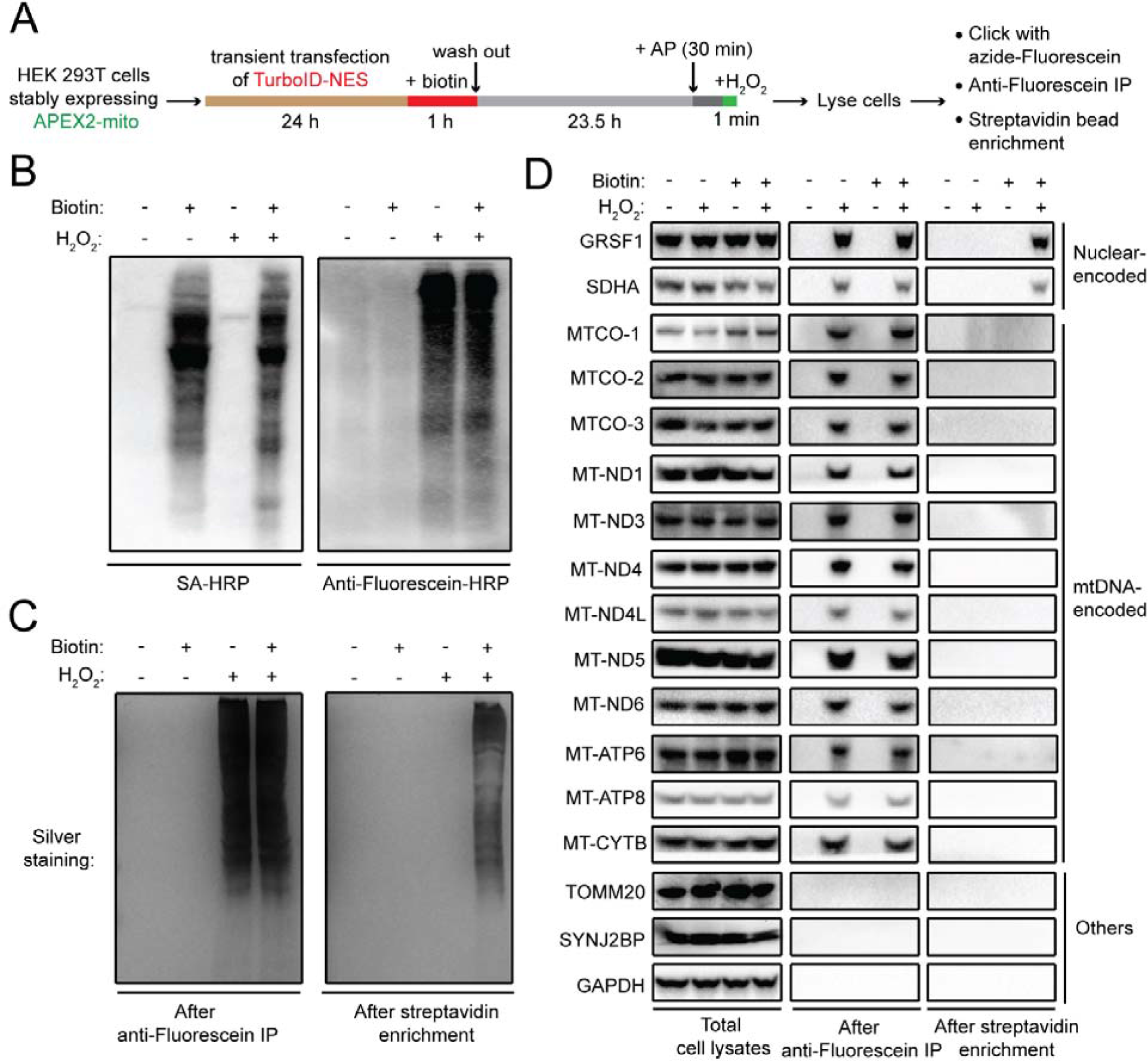
Validation of TransitID for cytosol to mitochondrial matrix proteome trafficking. **A**. Labeling protocol used. NES, nuclear export signal. **B.** Streptavidin and anti-fluorescein blotting of whole cell lysates from (**A**) along with negative controls omitting biotin or H_2_O_2_. **C**. Silver staining of enriched proteins after first anti-fluorescein IP (left) and after second streptavidin bead enrichment (right). **D**. Western blot detection of specific protein markers in samples from (A) after cell lysis (left), after anti-fluorescein IP (middle), and after the second streptavidin enrichment (right). GRSF1 and SDHA are true-positive nuclear-encoded mitochondrial proteins that are translated in the cytosol. mtDNA-encoded proteins are translated in the mitochondrion. TOMM20 and SYNJ2BP (OMM proteins) and GAPDH (cytosolic protein) are true negatives.

To check the spatial specificity of this tandem labeling procedure, we performed dual enrichment on these samples: Click chemistry with azide-FAM, followed by anti-FAM IP and streptavidin bead enrichment (**Figure 2C**). Proteins remaining after both enrichment steps should be those originating from the cytosol and trafficking into the mitochondrial matrix, namely nuclear genome-encoded mitochondrial proteins that are translated by cytosolic ribosomes before import into the mitochondrion. On the other hand, proteins remaining after just the first anti-FAM enrichment should represent *all* mitochondrial matrix proteins, including those encoded by the mitochondrial genome (mtDNA) and translated by mitochondrial ribosomes.

Western blotting in **Figure 2D** shows that all mitochondrial protein markers were detected after the first anti-FAM IP, as expected due to their proximity to mito-APEX2. However, after dual enrichment, only the nuclear-encoded mitochondrial proteins SDHA and GRSF1 remained, while the 12 mtDNA-encoded proteins were no longer detected, because these proteins are translated within the mitochondrion and were never proximal to cytosolic TurboID. In both single-enriched and dual-enriched samples, we did not detect the cytosolic protein GAPDH nor the outer mitochondrial membrane proteins SYNJ2BP or TOMM20, as expected. These data demonstrate the high specificity of TransitID and its ability to distinguish mitochondrial proteins based on their compartment of origin.

### TransitID resolves the spatial origin of nuclear-encoded mitochondrial proteins

Having validated the specificity of TransitID, we next asked whether we could use TransitID in a proteomic experiment to dissect the spatial origins of nuclear genome-encoded mitochondrial proteins (**Figure 3A**). All such proteins are translated in the cytosol, but previous studies have suggested that a portion are translated “locally”, near the outer mitochondrial membrane (OMM), to facilitate co-translational import and insertion of protein products^14, 15^. We prepared HEK 293T cells stably expressing APEX2 in the mitochondrial matrix and introduced either cytosolic TurboID-NES or TurboID-OMM (TurboID anchored to the outer mitochondrial membrane, facing the cytosol) by transient transfection (**Figure 3B**). TransitID was performed as in **Figure 3A** with a 24-hour chase period. We also prepared negative controls with H_2_O_2_ omitted (to prevent APEX labeling) or TurboID omitted. For each sample, on-bead trypsin digestion and TMT labeling were performed before pooling and analysis by liquid chromatography-tandem mass spectrometry (LC-MS/MS).

**Figure 3.**
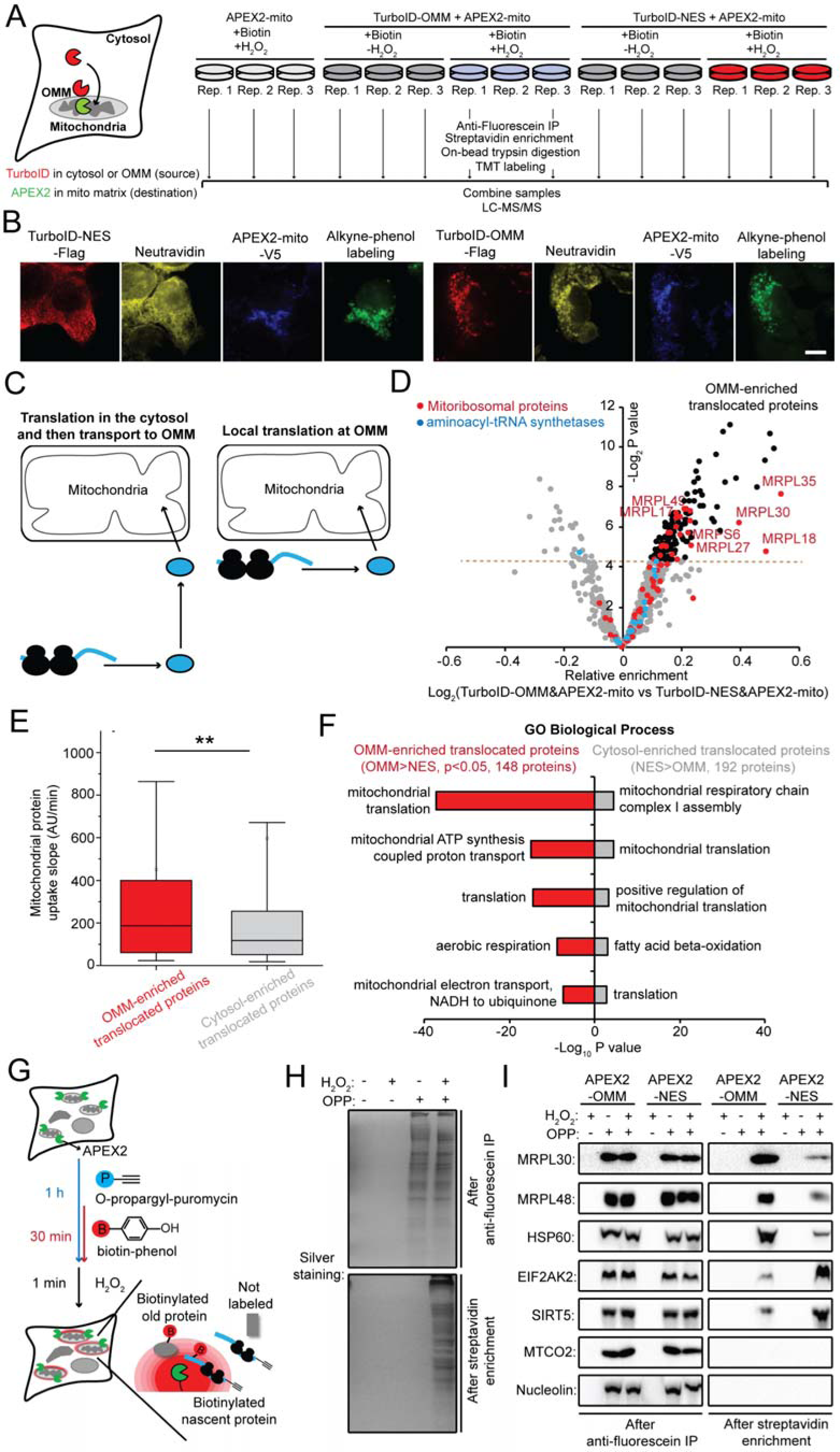
TransitID distinguishes mitochondrial matrix proteins by compartment of origin. **A.** Design of 15-plex TMT proteomic experiment. **B.** Confocal fluorescence imaging of TurboID- and APEX2-dual labeled samples. Neutravidin detects biotinylated proteins. Scale bar, 10 μ comparing local (at outer mitochondrial membrane, or OMM) versus distal (in cytosol) translation of mitochondrial proteins. **D.** Volcano plot showing relative enrichment of proteins in OMM-to-mito samples versus cytosol-to-mito samples. **E.** Comparison of mitochondrial protein uptake rates according to the mePROD^mt^ dataset^65^. **F.** GO biological process analysis of OMM-enriched translocated proteins and cytosol-enriched translocated proteins. **G.** Assay to detect newly synthesized proteins proximal to the OMM, via puromycin (OPP) tagging of new polypeptides followed by APEX2-OMM-catalyzed biotinylation. Puromycin-tagged proteins are enriched by Click chemistry and anti-fluorescein IP. **H**. Silver staining of samples from G, after anti-fluorescein IP (top) and after the second streptavidin bead enrichment step (bottom). **I**. Western blot detection of known protein markers in enriched samples. MRPL30, MRPL48 and HSP60 are enriched in our OMM-to-mito dataset; EIF2AK2 and SIRT5 are enriched in our cytosol-to-mito dataset. MTCO2 (mtDNA-encoded protein) and nucleolin (nucleocytoplasmic protein) are negative controls.

A total of 4210 proteins were detected with two or more unique peptides from this 15-plex experiment (**Table S1**). High correlation was observed across all biological replicates (**Figure S2A**). To determine the set of proteins that traffick from cytosol to mitochondria, we calculated protein enrichment (TMT ratio) in Turbo-NES/mito-APEX2 samples relative to each control (omit H_2_O_2_ or omit TurboID) (**Figure S2B**). Using a true positive list of known nuclear-encoded mitochondrial proteins and a true negative list of plasma membrane proteins, we plotted receiver operating characteristic (ROC) curves and determined TMT ratio cut-offs that maximized the difference between true positive rate (TPR) and false positive rate (FPR) (**Figure S2C**). Proteins with TMT ratios above the cut-offs as well as significant (*p*-value based) enrichment over controls were retained (**Figure S2D**). We obtained a final list of 670 proteins that traffick either from the cytosol or the OMM into the mitochondrial matrix (**Table S1**).

Analysis of specificity and sensitivity of this dataset showed comparable proteome quality to our previous single-step APEX1-mapped mitochondrial matrix proteome^16^ (**Supplementary Text 1**), with the important difference that all 13 mtDNA-encoded proteins were absent from our list, as expected due to the use of cytosolic or OMM TurboID in the first labeling step. A Venn diagram comparing the two datasets showed that 85 proteins identified by APEX1 were missed by TransitID (**Figure S3F**); interestingly, these proteins have much slower turnover rates on average (**Figure S3F**)^17^, and may therefore have lower flux from cytosol to mitochondrial matrix during the 24-hour chase period. 260 proteins identified by TransitID were missed by single-step mito-APEX1 PL. Within these 260 proteins, 42% have prior mitochondrial annotation, while many of the remaining 58% could be newly discovered mitochondrial proteins (mitochondrial “orphans”). We selected two of these proteins for detection by Western blot in purified mitochondrial fractions and confirmed both EIF2AK2 and SNAPC1. Both proteins were resistant to proteinase K treatment, suggesting that they reside within the mitochondrial matrix like our true positive markers MTCO2 and SDHA (**Figure S3G**).

By comparing protein enrichment scores in cytosol-to-mitochondria samples versus OMM-to-mitochondria samples, we identified 148 proteins that may preferentially originate from the OMM rather than cytosol (**Figure 3C-F**). These mitochondrial proteins are candidates for local translation at the OMM. To further explore this, we treated cells with O-propargyl-puromycin (OPP)^18, 19^ to tag newly synthesized proteins, then performed single-step proximity labeling with OMM-APEX2 or cytosolic APEX2-NES (**Figures 3G and S3H**). Dual enrichment was used to isolate proteins tagged by both OPP and APEX, i.e., newly synthesized proteins proximal to the OMM or in the cytosol (**Figure 3H**). We blotted the enriched material for three protein hits from our candidate list of 148 locally translated mitochondrial proteins. **Figures 3I and S3I** show that MRPL30, MRPL48 and HSP60 are all more abundant in OPP/APEX2-OMM samples than in OPP/APEX2-NES samples, in agreement with our TransitID proteomic data. The opposite pattern was observed for the cytosol-enriched translocated proteins EIF2AK2 and SIRT5. Thus, metabolic labeling of newly synthesized proteins in conjunction with APEX-catalyzed PL provides additional support for local translation of a specific subset of mitochondrial proteins. Additional analysis of this dataset is provided in **Supplementary Text 2**.

### TransitID for profiling stress-inhibited cytosol-to-nuclear protein translocation

Under stress, many proteins relocalize in order to prioritize essential processes such as protein quality control and gene expression. Failure to relocalize can result in cell dysfunction and even apoptosis^20, 21^. For example, cytosolic mislocalization of the nuclear protein TDP-43 under stress is known to drive neuronal dysfunction and serves as a pathological hallmark for several neurodegenerative diseases^22–24^.

To profile alterations in cytosol-to-nucleus proteome trafficking under stress, we prepared HEK 293T cells expressing both cytosolic TurboID-NES and nuclear APEX2-NLS (**Figure 4A**). After 10 minutes of TurboID-catalyzed biotinylation, we performed a 2-hour chase in the absence or presence of sodium arsenite-induced oxidative stress. Samples were finally labeled with APEX2-NLS and AP1 for 1 minute before cell fixation or lysis (**Figure 4B**). Microscopy showed correct localization of both PL enzymes and their labeled proteomes (**Figure 4B**), and orthogonal labeling was demonstrated by streptavidin and anti-FAM blotting (**Figure S4A**). After dual enrichment of biotin and alkyne-tagged proteins (**Figure S4B**), samples were blotted for known markers of nucleocytoplasmic shuttling as well as cytosol-resident markers, confirming specificity (**Figure S4C**).

**Figure 4.**
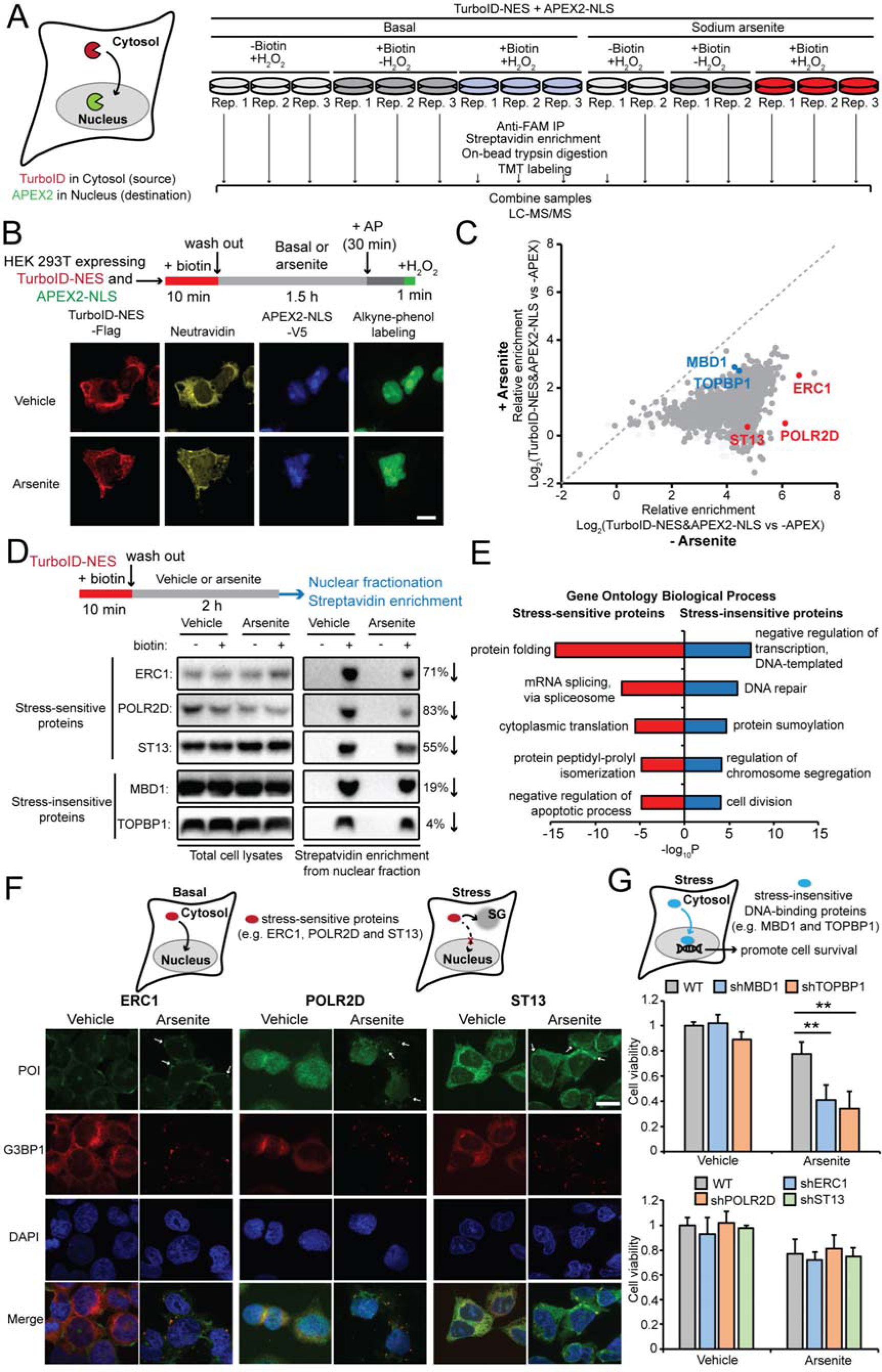
TransitID identifies proteins that traffick from cytosol to nucleus under stress. **A.** Design of 15-plex TMT proteomic experiment. **B.** Confocal fluorescence imaging of dual-labeled samples under basal and arsenite-induced stress conditions. Neutravidin detects biotinylated proteins. Scale bar, 10 μ **C**. Differential enrichment of cytosol-to-nucleus translocated proteins under basal (x axis) versus arsenite-treated stress conditions (y axis). Red-highlighted proteins are stress-sensitive proteins; blue-highlighted proteins are stress-insensitive proteins. **D.** Validation of stress-sensitive and stress-insensitive proteins using TurboID labeling followed by nuclear fractionation. **E.** GO biological process analysis of stress-sensitive and stress-insensitive cytosol-to-nucleus translocated proteins. **F.** Confocal imaging of three stress-sensitive cytosol-to-nucleus translocated proteins, with respect to endogenous G3BP1, a stress granule marker. Arrows point to translocated proteins at stress granules. Scale bars, 10 μ **G.** Knockdown of stress-insensitive translocated proteins (MBD1 or TOPBP1), but not stress-sensitive proteins (ERC1, POLR2D or ST13), impairs cell viability under arsenite stress.

We performed a 15-plex TMT proteomic experiment with three biological replicates each for basal and stressed conditions. Negative controls omitted biotin or H_2_O_2_ (**Figure 4A**). Using ROC-determined cutoffs and *p*-value filtering (**Figure S4D-F**), we obtained a total of 1791 cytosol-to-nucleus translocated proteins under basal conditions (**Table S2**). Nearly all of these proteins showed decreased nuclear import after stress (**Figure 4C**), with 6.6% (127 proteins) showing severe (>85%) inhibition of nuclear import. We validated three such proteins (ERC1, POLR2D and ST13) by performing single-step cytosolic TurboID labeling followed by a 2-hour chase and nuclear fractionation. Blotting of streptavidin-enriched material showed a marked reduction in nuclear abundance of all three proteins after arsenite-induced stress, consistent with our TransitID data (**Figure 4D**). By contrast, the enrichment of MBD1 and TOPBP1, two stress-insensitive proteins according to TransitID, were not obviously affected by arsenite treatment.

Stress-sensitive proteins that are inhibited from nuclear import may contribute to cellular defects under stress, due to loss of their nuclear functions. Gene Ontology analysis of our 127 severely stress-sensitive proteins showed high representation of processes related to protein folding and translation, and low representation of processes related to transcription and DNA repair (**Figure 4E**). Previous studies have suggested that some chaperones and translation factors may relocalize to stress granules (SGs) for protection during stress^25^; we therefore performed imaging of three stress-sensitive proteins from our TransitID dataset that have not previously been linked to SGs (**Figure 4F**). ERC1, POLR2D, and ST13 all colocalized with the SG marker G3BP1 upon arsenite treatment (**Figure 4F**). We also tested knockdown of two DNA-related stress-insensitive proteins (MBD1 or TOPBP1) and three stress-sensitive proteins (ERC1, POLR2D or ST13). We found that knockdown of each stress-insensitive protein impaired cell viability following arsenite stress, whereas knockdown of stress-sensitive proteins did not (**Figure 4G**). Perhaps the stress-insensitive DNA-binding proteins are critical cell regulators during the cell stress response, so that their efficient translocation to the nucleus must be preserved under such conditions.

### Mapping proteome trafficking between SGs and nucleoli during stress and recovery

Stress granules (SGs) are membraneless organelles that sequester mRNAs stalled in translation initiation in the face of cellular stress. Because SGs are dysregulated in numerous diseases^26^, studies have attempted to understand how SGs are formed, disassembled, and what role they play in protecting cells from oxidative, proteotoxic, heat, and UV stress. SGs share many components and characteristics with nucleoli, which also respond to stress by regulating ribosome biogenesis and sequestering nucleoplasmic proteins^27^. Specific proteins such as SGNP and HSP70 have been observed, by imaging, to shuttle between nucleoli and SGs under stress^28–30^. We used TransitID to explore the hypothesis that nucleoli may be a source of SG proteins during the cellular stress response.

To enable greater temporal precision, we used a variant of TurboID called LOV-Turbo (unpublished) that can be reversibly controlled with blue light. LOV-Turbo was targeted to nucleoli by fusion to three tandem copies of the nucleolar targeting sequence of NF-κB-inducing kinase (NIK3x), while APEX2 was targeted to SGs by fusion to the major SG protein G3BP1 (**Figure 5A**). These constructs were expressed in HEK 293T cells lacking endogenous G3BP1 (**Figure S5A**) and imaging showed correct targeting and expected labeling patterns, under both basal and arsenite-treated conditions (**Figure 5C**).

**Figure 5.**
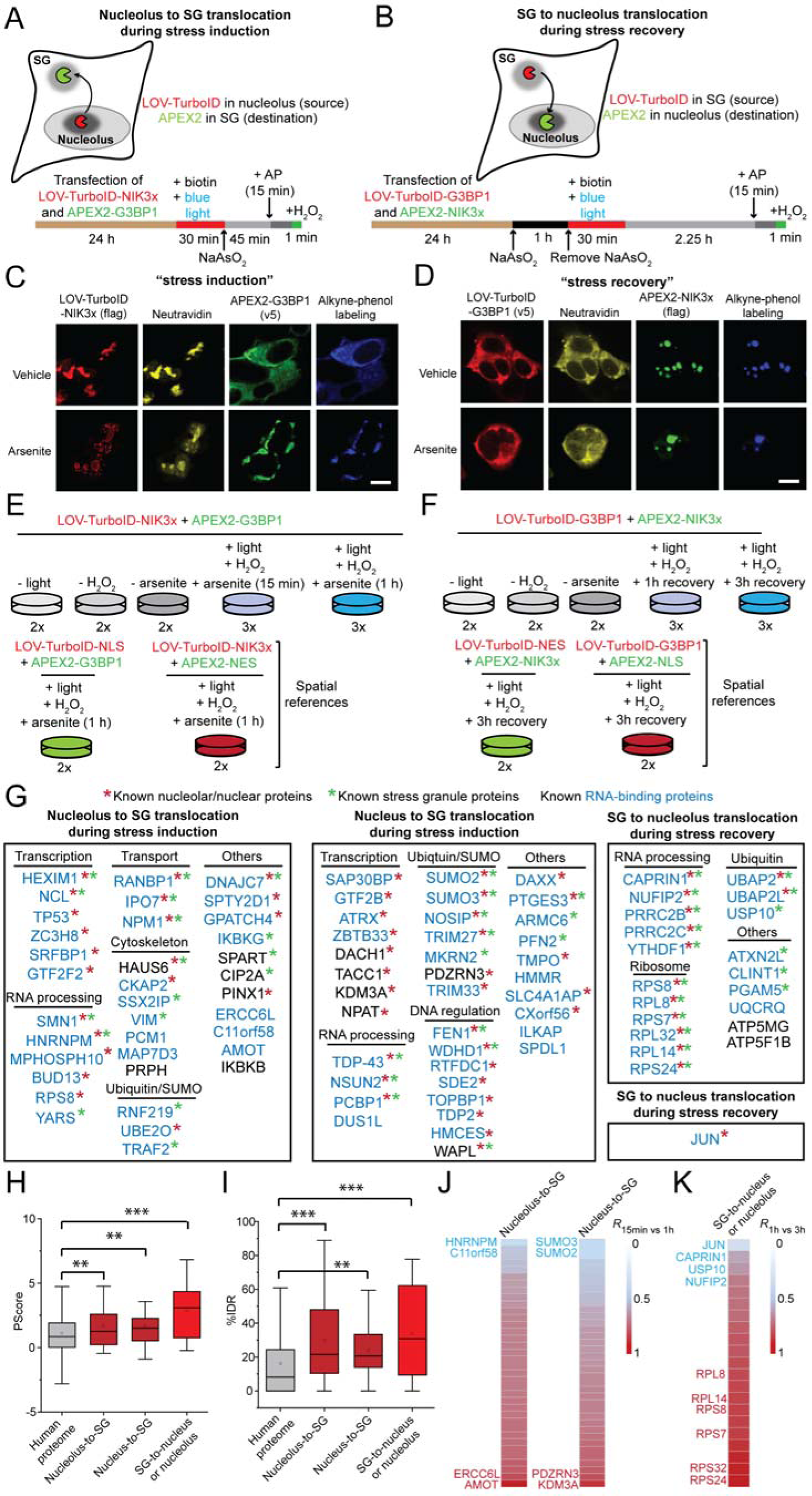
Proteome trafficking between nucleolus and stress granules mapped with TransitID. **A.** TransitID labeling of proteins that originate from the nucleus or nucleolus and traffick to SGs upon arsenite-induced oxidative stress. **B**. TransitID labeling of proteins that traffick from SGs to the nucleus or nucleolus during stress recovery. **C-D**. Confocal fluorescence imaging of dual-labeled samples corresponding to experiments in (**A**) and (**B**), respectively. Neutravidin detects biotinylated proteins. Scale bars, 10 μ **E-F** Design of proteomic samples corresponding to experiments in (**A**) and (**B**), respectively. **G.** Proteins enriched in the 4 indicated datasets. **H.** Phase separation propensity (PScore) of proteins enriched in each dataset in (**G**). **I.** Percent intrinsic disordered regions (%IDR) for proteins in each dataset in (**G**). **J.** Heat map showing proteins enriched in the faster (15 min) versus slower (1 hour) pulse-chase experiment. Red indicates faster-translocating proteins; blue indicates slower-translocating proteins. **K.** Same analysis as in (**J**), for the SG-to-nucleus or nucleolus dataset, comparing enrichment in the 1-hour versus 3-hour pulse-chase experiment. Red indicates faster-translocating proteins.

We performed TransitID as shown in **Figure 5A** by adding blue light and biotin to cells for 30 minutes, then inducing oxidative stress by treating with sodium arsenite for 15 minutes or 1 hour in the dark to suppress LOV-Turbo activity. APEX labeling was initiated with the addition of alkyne-phenol and H_2_O_2_ for 1 minute, then cells were lysed and dual enrichment was performed. Three replicate samples were prepared for proteomic analysis, along with negative controls omitting light, H_2_O_2_, or arsenite, and spatial references with LOV-Turbo expressed throughout the nucleus (LOV-Turbo-NLS) or APEX2 expressed throughout the cytosol (APEX2-NES) (**Figure 5E**). The 16-plex TMT experiment was analyzed by LC-MS/MS and 4822 proteins with 2 or more unique peptides were detected, with good correlation between biological replicates (**Table S3**). We filtered the data as shown in **Figure S5B-F**, using the –light and – H_2_O_2_ controls, basal sample, and the spatial reference samples. We obtained a final list of 36 stress-dependent (1 hour arsenite) nucleolus-to-SG translocated proteins, and 37 stress-dependent (1 hour arsenite) proteins that traffick from the nucleus to SGs (**Figure 5G and Table S3**).

Encouragingly, TDP-43, an ALS-associated protein known to shuttle from the nucleus to SGs during stress ^31^, was detected in our nucleus-to-SG list. We identified many other known SG proteins including RANBP1, PCBP1 and HNRNPM (**Figures 5G and S6D**). The majority of proteins in each list have prior nuclear annotation, and many known nucleolar proteins are present in our nucleolus-to-SG list (**Figure S6E-F**). By crossing with RBPbase (https://rbpbase.shiny.embl.de/), we confirmed that a large fraction of both lists were RNA binding proteins (86% and 84%), as expected based on the known biology of SGs and nucleoli (**Figure S6G**). These biomolecular condensates are also enriched in proteins with intrinsically disordered regions (IDRs) that can undergo phase separation^25^. We therefore analyzed Pscore^32^ and IUPred^33^, measures of phase separation propensity and intrinsic disorder rate, respectively, and found that both were significantly enriched in our datasets compared to the entire human proteome (**Figure 5H and 5I**).

Our proteomic experiment also included samples that were treated with arsenite for a shorter time period - 15 minutes instead of 1 hour. By comparing protein enrichments across these two conditions, we could identify proteins that translocate faster versus slower from nucleolus/nucleus to SGs. We found that proteins originating from the nucleolus generally arrive at SGs faster than those originating from the nucleus (**Figure 5J**). Among the faster trafficking proteins were PDZRN3 and TRIM33, two E3 ubiquitin ligases; perhaps these are involved in the stress-induced ubiquitination of SG constituents that primes SGs for disassembly^34, 35^.

When cells recover from oxidative stress, SGs dissolve on a timescale of 1 to 3 hours^36^. Even less is understood about how SGs disassemble than how they assemble. To explore the hypothesis that some SG proteins may traffick to the nucleus or nucleolus during stress recovery, we prepared HEK 293T cells expressing SG-targeted LOV-Turbo-G3BP1 and nucleolus targeted APEX2-NIK3x (**Figure 5B**). Cells were treated with sodium arsenite for 1 hour to induce SG formation, labeled with biotin in the presence of blue light for 30 minutes immediately after washing out arsenite, then cells were allowed to recover for another 2.5 hours. Finally, APEX2/alkyne-phenol labeling was performed for 1 minute in the nucleolus. Localization of fusion constructs and biotin/alkyne labeling patterns were assessed by imaging (**Figure 5D**). We designed a 15-plex proteomic experiment (**Figure 5F**) and filtered the mass spectrometry data as shown in **Figure S6A-C**. Our final list of 20 proteins that traffick from SGs to the nucleolus, and 1 protein (JUN) that trafficks from SGs to the nucleus during stress recovery are shown in **Figure 5G**.

Analysis performed as above showed that our SG-to-nucleolus dataset is highly enriched in known SG proteins, known nucleolus proteins, known RNA binding proteins, and intrinsically disordered domains (**Figures S6D-G, 5H-I**). Interestingly, three ubiquitin-related proteins, including the ubiquitin hydrolase USP10, were found in this dataset. Previous data suggest that stress-induced nucleolar aggresomes are enriched in ubiquitination^37^, so perhaps these translocated ubiquitin factors are involved in the dynamic regulation of ubiquitination in the nucleolus.

In our “stress recovery” proteomic experiment, we included samples with both shorter (1 hour) and longer (3 hours) chase periods. By comparing enrichment scores across these conditions, we observed that proteins such as CAPRIN1 and USP10 leave SGs more slowly, perhaps because they are known to be direct binding partners of the SG core scaffolding protein G3BP1 (**Figure 5K**)^38, 39^. By contrast, the trafficking of multiple ribosomal proteins is already complete at 1-hour post-stress removal; these may be new or damaged ribosomal subunits that are rapidly relocalized to restore ribosome assembly in the nucleolus.

We selected four proteins from our datasets for follow-up validation by orthogonal methods. By microscopy, we detected all proteins in SGs after arsenite treatment (**Figure S7A**) and heat-induced stress (**Figure S7B**), even though none have prior literature connections to SGs. We also immunoprecipitated the major SG protein G3BP1 and detected all four proteins associated with G3BP1 after arsenite treatment (**Figure S7C**). The interaction of DAXX and SRFBP1 with G3BP1 was abolished by high salt treatment, suggesting that the interaction is electrostatic in nature, while MPP10 and UBE2O’s interaction with G3BP1 showed both salt- and RNA-dependence.

Since DAXX was recently identified as a novel folding enabler^40^, we tested its effect on the aggregation of G3BP1 in vitro. A thermal shift assay showed that G3BP1’s solubility improved at higher temperatures in the presence of DAXX (**Figure S7D**). Perhaps DAXX has a chaperone role in the context of SGs.

### A role for stress granules in regulating the transcription factor JUN during cellular stress

We identified JUN in our TransitID experiment as a protein that trafficks from SGs to the nucleus during stress recovery. JUN is a major component of the transcription factor AP-1 and critical for cell cycle progression and tumor growth ^41^, but it has never previously been linked to SGs. Environmental stresses, such as UV light, upregulate JUN’s transcriptional activity and complexation with its binding partner FOS. Stress is also known to increase JUN phosphorylation by JUN N-terminal kinases (JNKs)^42, 43^, and upregulate JUN’s own transcription via an autocrine amplification loop at its promoter^44^. However, many aspects of JUN’s response to stress are unclear, such as whether JUN’s subcellular localization and biophysical properties change under stress, and how these in turn affect its transcriptional activity.

To validate our proteomic finding, we first performed fluorescence microscopy. HeLa cells treated with sodium arsenite for 1 hour showed cytosolic JUN puncta that overlapped with the endogenous SG marker protein FXR1 (**Figure 6A**). Approximately 20% of total JUN was observed in these puncta. After 3 hours of stress recovery, cytosolic JUN puncta were no longer visible, and JUN was detected exclusively in the nucleus (**Figure 6A**).

**Figure 6.**
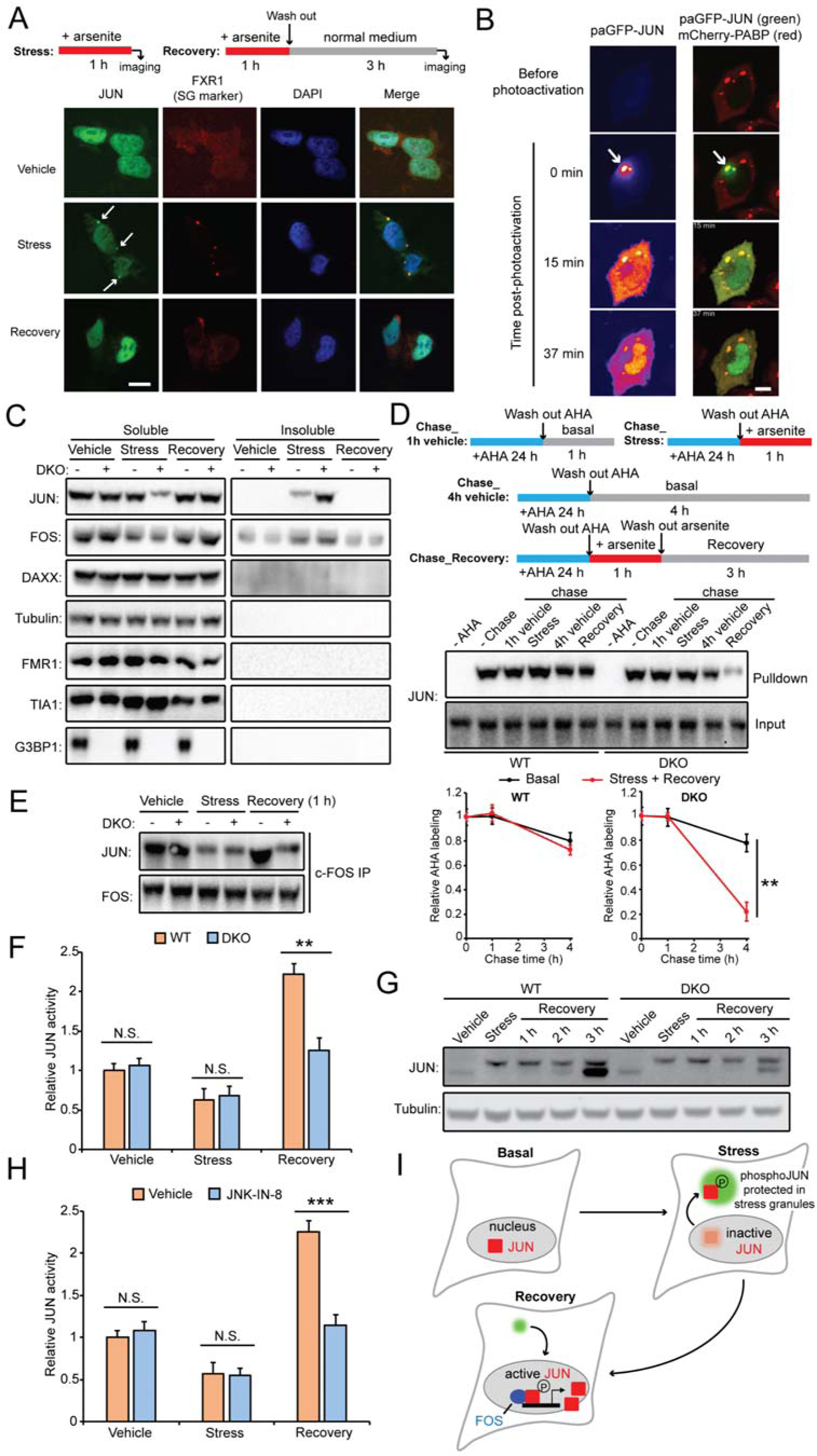
Stress granules protect JUN from degradation and enable rapid recovery from stress. **A.** Confocal fluorescence imaging of endogenous JUN under basal, stress and recovery conditions. Arrows point to JUN at stress granules. Scale bar, 10 m. **B.** Imaging of HeLa cells expressing photoactivatable JUN (paGFP-JUN) and a stress granule ma ^μ^ (mCherry-PABP) during stress recovery. Immediately after arsenite washout, the indicated region (white arrow) was activated by 405-nm laser and GFP was imaged at 200 millisecond intervals. Scale bar, 10 μ **C.** Analysis of JUN aggregation. Total JUN and several other proteins were quantified in soluble versus SDS-resistant insoluble fractions from lysates of wild-type HEK 293T cells and G3BP1&2 double knockout (DKO) cells. **D.** Analysis of JUN degradation by metabolic labeling with AHA followed by western blot detection of AHA-tagged (streptavidin-enriched) JUN. Quantification from three biological replicates shown below. **, p< 0.01. **E.** Detection of JUN complexation with FOS in wild-type HEK 293T versus in G3BP1&2 KO (DKO) cells. **F.** Measurement of JUN activity via ELISA DNA-binding assay. **, p< 0.01. **G.** Western blot detection of total JUN protein in wild-type HEK 293T cells and in G3BP1&2 DKO cells incapable of forming stress granules. Tubulin is a loading control. **H.** Measurement of JUN DNA binding activity by ELISA, with and without inhibition of JNK. **I.** Model for how SGs protect JUN during cellular stress.

Because static images do not rule out the possibility that SG-localized JUN originates from a non-nuclear pool of JUN, we also performed time-lapse microscopy of JUN fused to photoactivatable GFP (paGFP). After irradiation of exclusively nuclear JUN-paGFP under basal conditions, cells were stressed with arsenite and timelapse microscopy was performed. Fluorescent JUN-paGFP puncta were detected in the cytosol within 1 hour (**Figure S8A, supplementary movie 1**). To track JUN movement from SGs to the nucleus during stress recovery, we specifically activated JUN-paGFP within SGs, indicated by co-transfected SG marker mCherry-PABP, immediately after removing arsenite. We observed relocalization of this JUN-paGFP population to the nucleus on a timescale of 15-40 minutes (**Figure 6B, supplementary movie 2**).

We also performed a co-IP experiment to validate JUN localization to SGs. When we pulled down the endogenous SG protein G3BP1, we also detected endogenous JUN post-stress, and the interaction was abolished by treatment with high salt, suggesting that JUN’s interaction with G3BP1 is electrostatic in nature (**Figure S8B**).

Having established that a subpopulation of JUN relocalizes to SGs during stress, we wondered what is the functional purpose of this relocalization? JUN contains a C-terminal leucine zipper domain that it uses to bind to its DNA promoter. Due to the high density of positive charges, this domain is disordered in solution, but forms an α-helical structure upon DNA binding^45^. Disordered cationic proteins are highly prone to aberrant condensation, as seen in neurodegenerative disease^46, 47^. Recent work on TDP-43 suggests that TDP-43 recruitment to SGs protects it from irreversible aggregation in the cytosol^48, 49^. We hypothesized that the same may be true for JUN; perhaps JUN is prone to aggregation during cellular stress, and localization to SGs helps to protect JUN from irreversible aggregation and dysfunction.

To test this, we prepared lysates from basal, stressed, and stress-recovered cells, and partitioned the lysates between soluble and SDS-insoluble fractions by centrifugation. We blotted the fractions for endogenous JUN and its protein-interaction partner FOS. Interestingly, only JUN but not FOS or other control proteins showed significant aggregation upon stress. These JUN aggregates were insensitive to 1,6-hexanediol, which disperses condensates (**Figure S8C**). We repeated the experiment in HEK 293T cells without the ability to form SGs (G3BP1/G3BP2 double knock out or DKO cells). In DKO cells, JUN aggregation post-stress was strongly exacerbated (**Figure 6C**). Hence, JUN may relocalize to SGs to reduce its tendency to aggregate in the face of cellular stress.

We wondered if JUN aggregation may cause it to be more rapidly degraded. To monitor JUN degradation, we performed metabolic labeling of JUN in cells with L-Azidohomoalanine (AHA)^50^. We then chased for 4 hours under basal conditions or with arsenite for the first hour followed by 3 hours of recovery. We detected ∼25% loss of AHA-tagged JUN under both conditions. However, in DKO cells unable to form SGs, JUN degradation was dramatically increased to ∼80% following stress and recovery (**Figure 6D**). These results suggest that SGs play an important role in reducing both the aggregation and degradation of JUN following oxidative cellular stress.

We next probed the role of JUN’s SG localization on its DNA binding and transcriptional activity. Three separate experiments, detailed in **Supplementary Text 3**, show that when cells are unable to form SGs, JUN’s interaction with FOS (**Figure 6E**), DNA binding activity (**Figure 6F**), and ability to upregulate its own synthesis (**Figure 6G**) are all impaired during stress recovery.

Previous work has shown that environmental stresses such as UV light increase JUN phosphorylation at Ser63 and Ser73 via the action of Jun N-terminal kinase (JNK)^42, 43^. We also found that arsenite treatment increased JUN phosphorylation in HEK 293T cells (**Figure 6G**), and addition of the JNK inhibitor JNK-IN-8 prevented this increase (**Figure S8F**). More interestingly, JNK inhibition prevented upregulation of JUN’s protein level and transcriptional activity during stress recovery (**Figures S8F and 6H**). Taken together (**Figure 6I**), our results support a model in which JUN relocalizes to SGs during cellular stress, which may serve the beneficial purpose of alleviating JUN aggregation and degradation, resulting in enhanced restoration of transcriptional activity upon stress recovery.

### Detection of intercellular protein communication between cancer cells and macrophages by TransitID

In addition to mapping intracellular proteome dynamics, we explored the use of TransitID for dissecting intercellular protein signaling. Given that most cells within an organism express largely the same set of proteins, it is very challenging to identify which proteins have originated by intercellular transfer from another cell rather than by translation within the cell type of interest. Yet dissecting such communication is vital to understanding how different cell types signal and cross-regulate one another to achieve tissue and organ function. One setting in which intercellular signaling is both crucial and complex is the tumor microenvironment (TME), where cancer cells and a variety of immune cells interact and extensively influence one another. In particular, tumor-associated macrophages (TAMs) can be hijacked via signals from cancer cells to secrete immunosuppressive signals, facilitating immune evasion by tumors^51^. In the other direction, TAMs secrete signals that promote tumor development^52^. This bidirectional signaling between cancer cells and TAMs is a topic of intense study and has identified TAMs as an important therapeutic target in the treatment of cancer^53^.

Given the lack of methods to systematically probe such intercellular signaling, we sought to apply TransitID to study proteome communication between macrophages and cancer cells. We prepared co-cultures of MC38 colon cancer cells stably expressing cytosolic TurboID-NES with Raw264.7-ASC macrophage cells stably expressing cytosolic APEX2-NES (**Figure 7A**). TransitID was performed with a 4-hour chase period. Fluorescence imaging showed that TurboID and APEX2 remained separate after the experiment (**Figure 7B**) and Western blot analysis revealed no cross-labeling between PL enzymes (**Figure S9A**), indicating that the PL enzymes themselves did not transfer between the two cell types. Western blot analysis of enriched material showed that the proteins CDC42 and PTEN traffick between the tumor cytosol and macrophage cytosol (**Figure 7C-E**). Both proteins have previously been detected in cancer-derived exosomes^54, 55^, but the destination compartment within macrophages was unknown. We found that treatment of cultures with the exosome biogenesis inhibitor GW4869 reduced the transfer of both proteins to macrophage cytosol (**Figure 7E**). As PTEN is a phosphatase, cytosolic access may enable it to reprogram the macrophage phosphorylation signaling as part of tumor-mediated “hijacking” of TAM function^55^.

**Figure 7.**
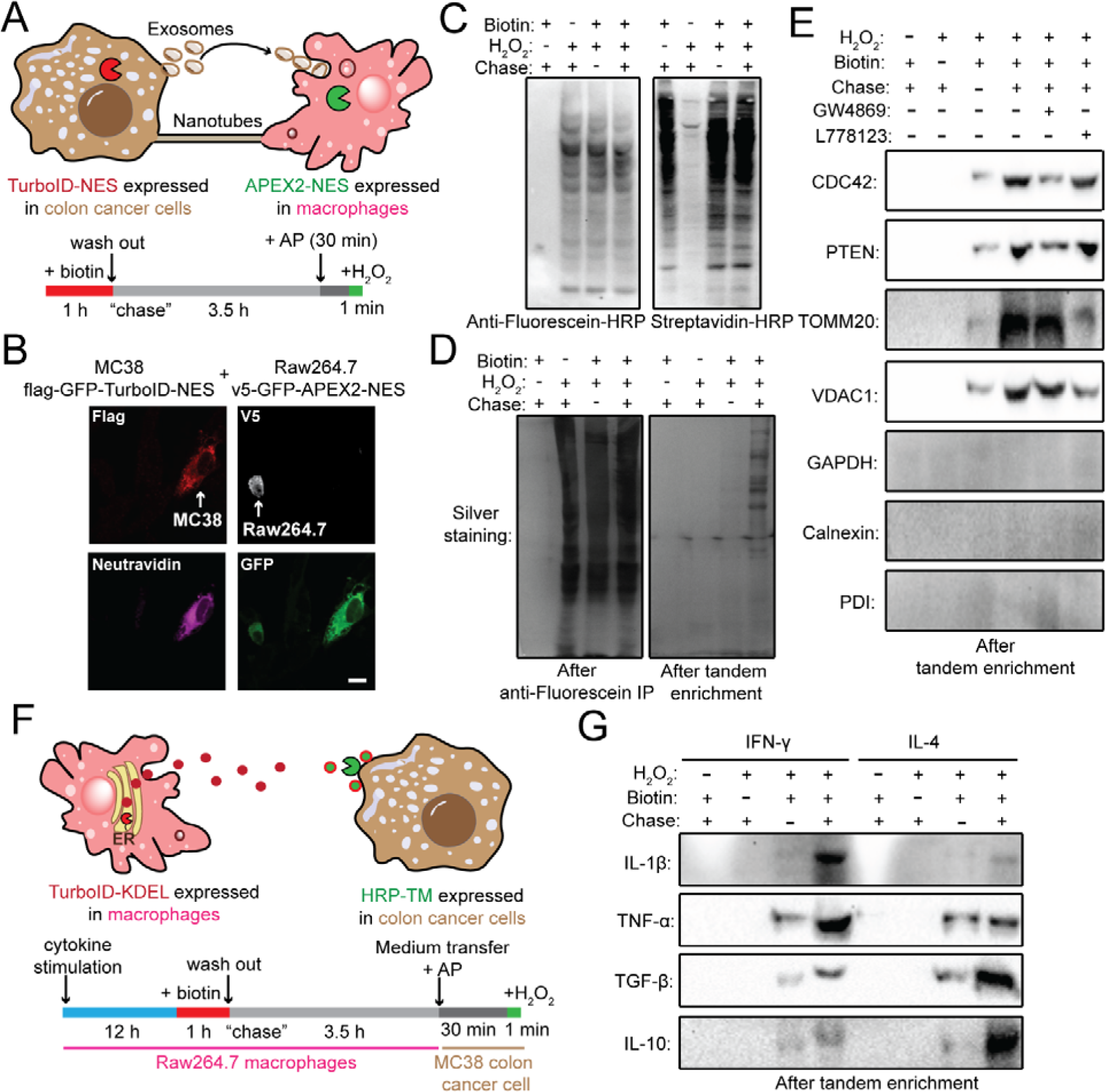
Detection of intercellular protein communication between cancer cells and macrophages by TransitID. **A.** TransitID labeling of proteins that originate from the cytosol of cancer cells and traffick to the cytosol of macrophages. NES, nuclear export sequence. AP, alkyne-phenol. **B.** Confocal fluorescence imaging of co-cultured MC38 cancer cells (expressing Flag-GFP-TurboID-NES) and Raw264.7 macrophages (expressing V5-GFP-APEX2-NES). Neutravidin detects biotinylated proteins. Scale bar, 10 μ **C.** Streptavidin and anti-fluorescein blotting of whole cell lysates from (**A**). **D.** Silver staining of enriched proteins after first anti-fluorescein IP (left) and after second streptavidin bead enrichment (right). **E.** Western blot detection of specific protein markers in samples from (**D**) after tandem enrichment. CDC42 and PTEN are known exosome cargoes^54, 55^ that can activate cytosolic signaling in recipient macrophages. TOMM20 and VDAC2 are mitochondrial proteins that can traffick between cells via nanotubes^57^. Cytosolic GAPDH, ER calnexin, and plasma membrane PDI are negative controls. GW4869 and L778123 are small-molecule inhibitors of exosome biogenesis and tunneling nanotube formation, respectively. **F.** TransitID labeling of proteins that are secreted from macrophages and traffick to the surface of cancer cells following cytokine stimulation. The medium from Raw264.7 cells was collected and added to MC38 cells. HRP-TM, horseradish peroxidase fused to a transmembrane helix. **G.** Western blot detection of specific protein markers in samples from (**F**) after tandem enrichment. IL-1β and TNF-α are cytokines known to be released by M1-type macrophages stimulated by IFN-γ^58^. TGF-β and IL-α re cytokines known to be released by M2-type macrophages stimulated by IL-4^59^.

In addition to exosome-mediated protein transport, we also wondered whether nanotubes, known to form between cancer cells and macrophages^56^, could enable intercellular communication in our co-culture system. We blotted TransitID-enriched material for the outer mitochondrial membrane proteins TOMM20 and VDAC1 both because they are accessible to cytosolic PL enzymes and because mitochondria have been shown to transit through nanotubes^57^. We detected both proteins, but not when the nanotube inhibitor L778123 was applied (**Figure 7E**). Control proteins calnexin (ER) and PDI (cell surface) were not detected as transported proteins.

We also used TransitID to probe proteome communication in the opposite direction, from macrophages to cancer cells. To represent two alternative macrophage states, we treated the cultures with IFN-γ to drive macrophages to a pro-inflammatory “M1” state^58^, or with IL-4 to drive them to an anti-inflammatory “M2” state^59^. TurboID was expressed in the ER lumen of macrophages to label their secreted proteomes, while horseradish peroxidase was expressed on the surface of MC38 colon cancer cells to capture macrophage-derived proteins (**Figure 7F**).

We detected the transfer of inflammatory cytokines IL-1β and TNF-α from M1 macrophages, and the transfer of anti-inflammatory cytokines TGF-β and IL-10 from M2 macrophages. HRP enriched all these cytokines on the surface of cancer cells in both medium-transfer and co-culture systems (**Figure 7G** and **S9B**). Collectively, our results show that TransitID can be used in a versatile manner to investigate intercellular protein trafficking, with subcellular resolution in both source and destination cells.

## Discussion

Proximity labeling has provided a powerful means to capture spatial information in MS experiments. However, until now information about the dynamic movement of proteins from one subcellular compartment to another or from one cell to another has been largely inaccessible by MS. The TransitID methodology introduced here enables the identification of proteins translocating between compartments and cells.

Several recent studies have combined TurboID labeling in “source” tissue with sample dissection and streptavidin enrichment at “destination” regions to map secreted proteome dynamics in living flies^60^ and mice^61, 62^. This methodology works well for long-range proteome communication between tissue regions that can be separated by dissection. However, for short-range proteome trafficking between intermingled cells, or intracellular trafficking between organelles, this approach cannot be applied. Furthermore, published approaches do not enable users to define the destination compartment in recipient cells (e.g., cytosol, cell surface, endosome, etc.). We developed TransitID as a general and flexible solution to the problem of mapping any intercellular or intracellular proteome trafficking, with the ability to precisely define both source and destination compartments of interest.

The main challenge to developing TransitID was the establishment of orthogonal labeling and enrichment chemistry for APEX. The alkyne-phenol AP1 probe and azide-fluorescein-based enrichment developed here are highly efficient and should be useful even for single enzyme PL applications. By avoiding the use of biotin and streptavidin, our protocol circumvents background from endogenous biotinylated proteins, which are especially abundant in vivo and limit signal-to-noise in such PL experiments. Even in cell culture, our side-by-side comparison to conventional BP labeling shows that protein recovery is ∼5-fold higher with AP1/azide-fluorescein, likely due to the increased membrane permeability of AP1.

We demonstrated three intracellular applications of TransitID and one intercellular application. The resulting datasets should serve as valuable resources that can be mined for insights into local translation of mitochondrial proteins, the impact of stress on nucleocytoplasmic protein shuttling, and the dynamics of SG formation and disassembly. Each dataset was accompanied by some follow-up validation, but we performed the most in-depth study on the transcription factor JUN, which we showed unexpectedly relocalizes to SGs in response to cellular stress. We go on to show that this recruitment is essential to prevent JUN aggregation and degradation during cellular stress, thereby enabling faster and more complete restoration of JUN transcriptional function upon stress recovery.

The application of TransitID to study protein communication between macrophages and cancer cells highlights the potential of this technique for dissecting intercellular signaling. If repeated *in vivo* and with full proteome analysis, it might be possible to gain insights into specific mechanisms used by tumor cells to hijack macrophages, and signals used by macrophages to promote tumor development. There is a plethora of other intercellular signaling questions that could be probed with TransitID, such as how activated microglia, reactive astrocytes, and neurons signal to one another in the context of neurodegeneration^63^, and how T cell activation induces tumor cell killing via effector secretion^64^. As scRNA-seq and in situ RNA-seq continue to shed light on the full diversity and organization of cell types in tissue, the need to understand the molecular mechanisms by which these cell types signal to and coordinate with one another will only increase.

With new and improved PL enzymes, it may be possible to further streamline TransitID to achieve faster temporal resolution, greater sensitivity, and a simpler enrichment protocol. Such improvements may make it possible to also combine PL enzymes for simultaneous, rather than tandem, labeling – opening the door to highly specific mapping of membrane contact sites or protein subcomplexes in vivo.

## Acknowledgements

We are grateful to the NIH (U24-CA210986 to S.A.C., U01-CA214125 to S.A.C., R01-DK121409 to S.A.C. and A.Y.T., RC2DK129964 to A.Y.T., R35NS097974 to J.P.T., T32GM007276 to J.S.C.), Stanford Wu Tsai Neurosciences Institute, NSF (GRFP DGE-1656518 to J.S.C., WALII, DBI grant # 2213983 to S.B) and the Stanford Gerald J. Lieberman Fellowship (J.S.C.) for support of this work. We also acknowledge the generous support of the Dr. Miriam and Sheldon G. Adelson Medical Research Foundation to S.A.C. and N.D.U. J.P.T. acknowledges funding from Howard Hughes Medical Institute. S.B. acknowledges funding from CPRIT (RR220094). We thank Dr. Hong Joo Kim for helpful discussions.

## Author contributions

W.Q., J.S.C. and A.Y.T. designed the research and analyzed all the data except where noted. W.Q. and J.S.C performed all experiments except where noted. W.Q., J.S.C., A.Y.T., C.X., N.D.U. and S.A.C. designed the proteomics experiments. W.Q. and J.S.C. prepared the proteomic samples. C.X. processed the proteomic samples and C.X., N.D.U., W.Q., and J.S.C. analyzed the MS data. J.M. and B.D.F. performed photoactivation imaging experiments. S.B. and J.P.T. advised experiments on stress granules. W.Q., J.S.C. and A.Y.T. wrote the paper with input from all authors.

## STAR METHODS

### KEY RESOURCE TABLE

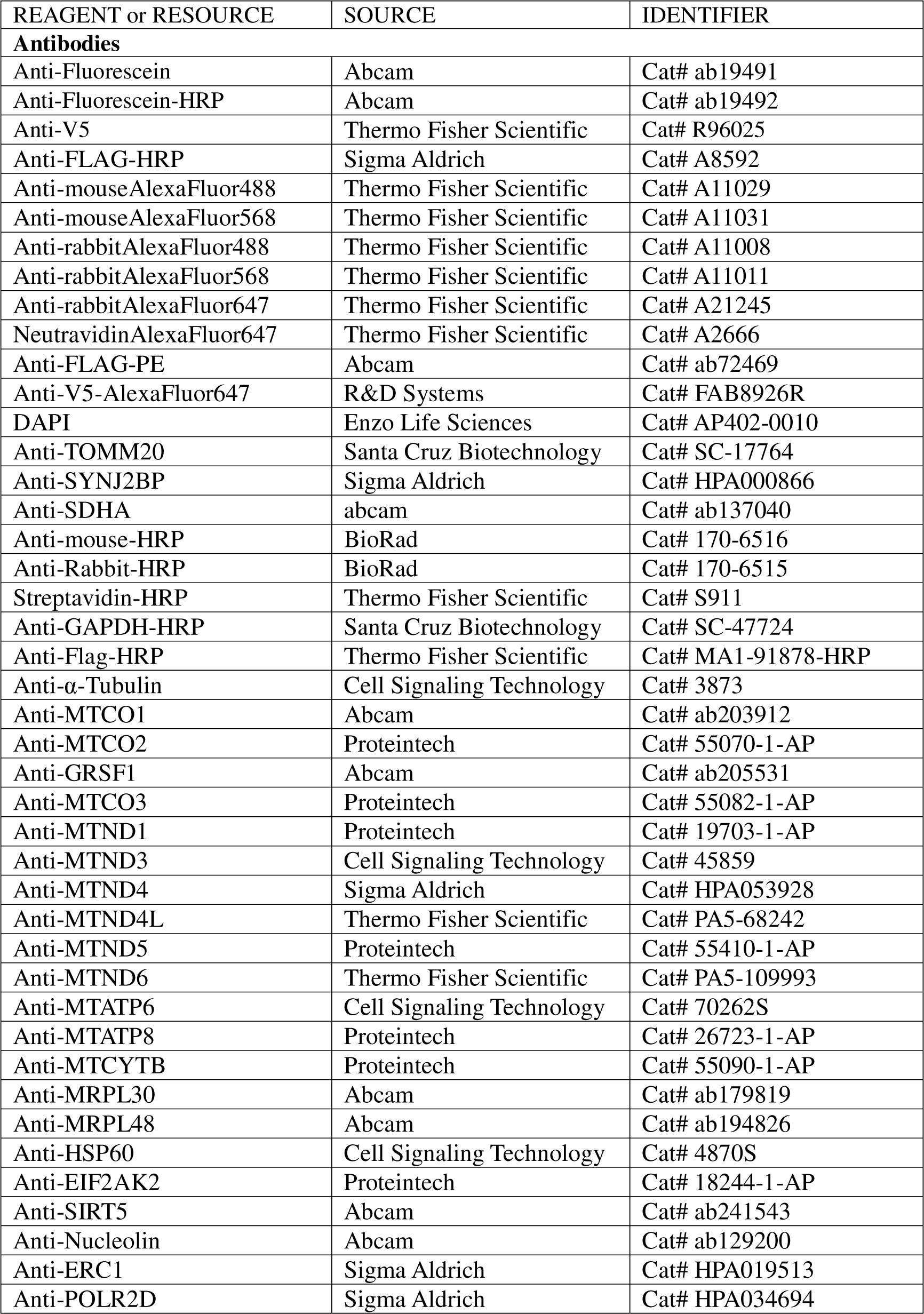
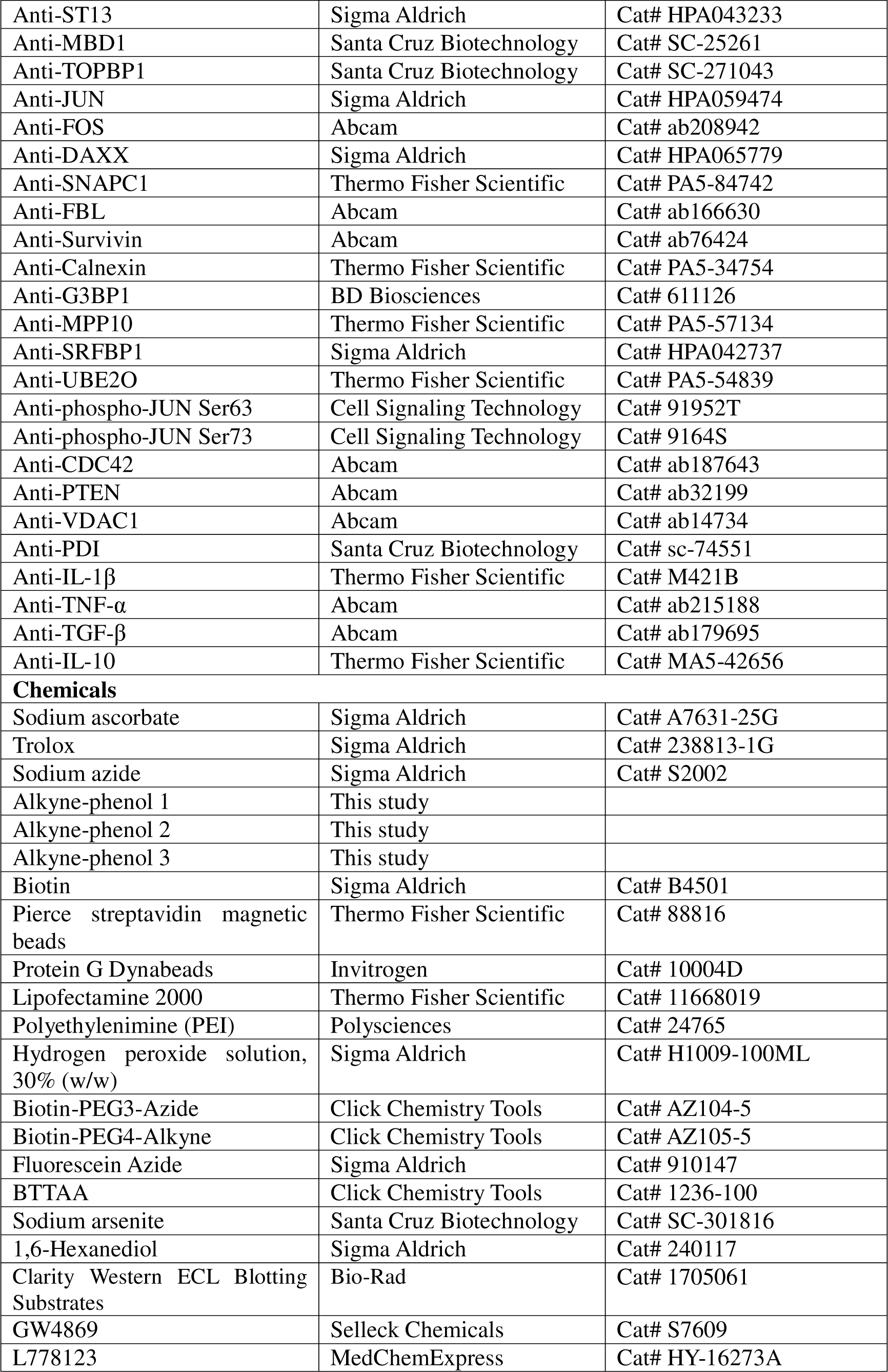
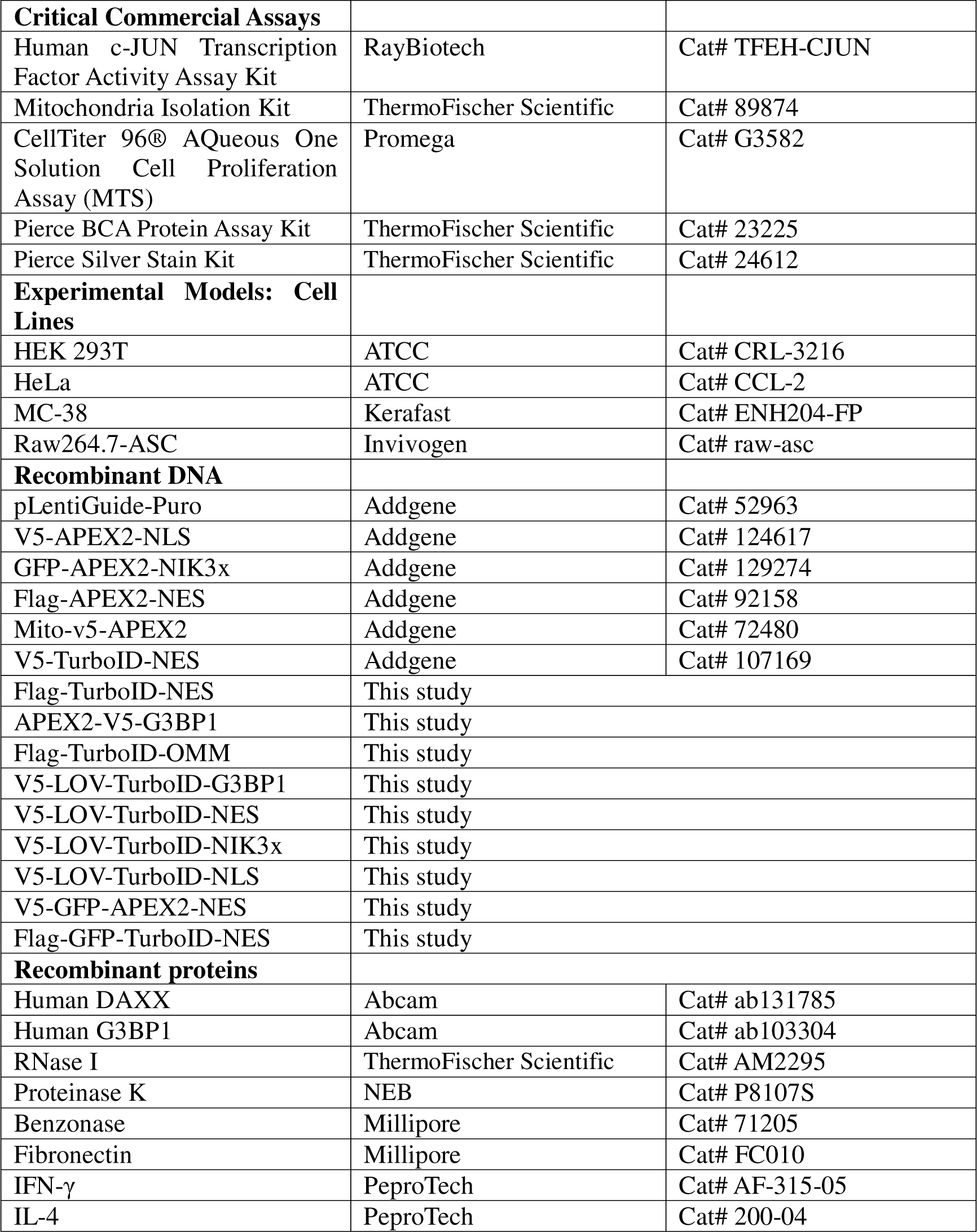

#### Cell Culture

HEK 293T cells from the ATCC (passages <25) were cultured in a 1:1 DMEM/MEM mixture (Cellgro) supplemented with 10% fetal bovine serum, 100 units/mL penicillin, and 100 mg/mL streptomycin at 37 under 5% CO_2_. HeLa (ATCC), MC-38 (Kerafast) and Raw264.7 cells expressing ASC (Invivogen) were cultured in the DMEM medium supplemented with 10% fetal bovine serum, 100 units/mL penicillin, and 100 mg/mL streptomycin at 37 under 5% CO_2_. For fluorescence microscopy imaging experiments, cells were grown on 7 × 7-mm glass coverslips in 24-well plates. For TransitID experiments, cells were grown on 15-cm glass-bottomed Petri dishes (Corning). To improve the adherence of HEK 293T cells, glass slides and plates were pre-treated with 50 mg/mL fibronectin (Millipore) for 20 min at 37 before cell plating and washing three times with Dulbecco’s PBS (DPBS) (pH 7.4). HEK 293T cells stably expressing APEX2-NLS, APEX2-NIK3x, APEX2-OMM and APEX2-NES were generated in our previous studies^7, 66^.

Stable cell lines were generated though infection by lentivirus. To generate lentivirus, HEK 293T cells plated at approximately 70% confluency in a 6-well dish were transfect with 1000 ng of the plasmid of interest, 750 ng of psPAX2, and 250 ng of pMD2G in 160 μL of serum-free DMEM with 12 μL of PEI. After 2 days, the supernatant containing the lentivirus was then syringe-filtered through a 0.45 μm filter. 300 μL of the crude supernatant was then added on to a 6-well dish of approximately 50% confluent HEK 293T. After 2 days, cells were passage and selected with 2 μg/mL of puromycin, 10 μg/mL blasticidin, or 500 μg/mL hygromycin for 1 week.

To generate G3BP1 KO cells, lentivirus was generated using a gRNA against G3BP1 (GTAGTCCCCTGCTGGTCGGGC)^67^ cloned into pLentiGuide-Puro (Addgene) then used to infect HEK 293T cells stably expressing Cas9^68^. After selection with puromycin, clonal cell lines were generated and tested by western blot staining against G3BP1 to identify a complete knockout cell line.

Lentivirus was used to generate stable MC38 and Raw264.7-ASC cells. To generate the lentivirus, HEK 293T cells were plated in a T75 flask to 70% confluency then transfected with 4,000 ng of the plasmid of interest, 3,000 ng of psPAX2, and 1,000 ng of pMD2G in 500 μL of OptiMEM and 30 μL of PEI. The following day, the lentivirus containing supernatant was filtered and stored at 4 °C. 10 ml of media replaced onto the cells. Additionally, 3E5 MC38 or Raw264.7-ASC cells were also plated in a 6-well dish the same day. The next day, the lentivirus containing supernatant was filtered from the transfected cells and combined with the prior day’s supernatant. 2 mL of crude supernatant was added to the MC38 or Raw264.7-ASC plated the day before. The media was replaced with crude lentivirus containing supernatant the following day. The next day, infected cells were passage and selected with the appropriate antibiotic for a week.

#### TransitID cell labeling protocol

For both Western blotting and proteomic analysis, HEK 293T cells stably expressing the APEX2 fusion construct of interest were cultured in a 15-cm dish for 18-24 hours to about 90% confluency. The cells were transiently transfected with the TurboID-fusion construct of interest using lipofectamine 2000 (Thermo Fisher) for 24 hours. TurboID labeling was initiated by adding a final concentration of 50 μM biotin for 10 minutes (or 1 hour where indicated). The labeling was stopped by transferring the cells to ice and washing five times with ice-cold DPBS. The cells were cultured in the normal medium for the indicated “chase” time at 37 APEX labeling was initiated by changing to fresh medium containing 50 μM alkyne-phenol and incubating at 37 under 5% CO_2_ for 30 min. This pre-incubation step is necessary to ensure that cells are loaded with alkyne-phenol but APEX labeling does not begin until H_2_O_2_ (Sigma Aldrich) is added to a final concentration of 1 mM for 1 min, with gentle agitation. The reaction was quenched by replacing the medium with an equal volume of quenching solution (10 mM ascorbate, 5 mM Trolox and 10 mM sodium azide in DPBS). Cells were washed with quenching solution three times to remove excess probe. Cells were washed twice with 10 mL ice-cold DPBS, harvested by scraping, pelleted by centrifugation at 1,400 r.p.m. for 3 min, and either processed immediately or flash frozen in liquid nitrogen and stored at -80°C before further analysis.

For cytosol-to-mitochondria protein translocation, HEK 293T cells stably expressing the APEX2-mito were transiently transfected with TurboID-NES or TurboID-OMM for 24 hours. TurboID labeling was initiated by adding a final concentration of 50 μM biotin for 1 hour, then chase was performed for 24 hours, and followed by 1-minute APEX labeling. For cytosol-to-nucleus protein translocation, HEK 293T cells stably expressing the APEX2-NLS were transiently transfected with TurboID-NES for 24 hours. TurboID labeling was initiated by adding a final concentration of 50 μM biotin for 10 minutes, then chase was performed for 2 hours, and followed by 1-minute APEX labeling. To analyze the dynamics of nuclear import under stress, the cells were treated with 500 μM sodium arsenite during the 2-hour chase.

For stress granules-to-nucleolus or -nucleus protein translocation, HEK 293T cells with G3BP1 knocked out were transfected with either V5-LOV-Turbo1-G3BP1 or V5-LOV-Turbo1-NES from a tetracycline inducible promoter and either GFP-APEX2-NIK3x or V5-APEX2-NLS for 24 hours with 400 ng/mL doxycycline to induce expression of the constructs. For each replicate sample in our proteomic experiment, cells were plated on a 10-cm dish. Cells were treated with 500 μM of sodium arsenite 1 hour followed by 100 μM biotin for LOV-Turbo1 labeling with blue light for 30 minutes. Blue light was administered with an AMUZA blue LED array and LED Array Driver setup. The driver was set at 12 V and the pulse generator was set at 2.3V on a 10% duty cycle (10 ms on/90 ms off). The media was then washed out for a 30-minute or 2.5-hour chase, giving a total recovery period of either 1 hour or 3 hours. 15 minutes prior to APEX labeling, alkyne-phenol was added. APEX labeling was then performed for 1 minute after the chase period.

For nucleolus- or nucleus-to-stress granule protein translocation, HEK 293T cells with G3BP1 knocked out were transfected with either V5-LOV-Turbo1-NIK3x or V5-LOV-Turbo1-NLS from a tetracycline inducible promoter and either V5-APEX2-G3BP1 or flag-APEX2-NES for 24 hours with 400 ng/mL doxycycline to induce expression of the constructs. For each proteomic replicate, cells were plated on a 10-cm dish. Cells were treated with 100 μM biotin and blue light as described earlier for LOV-Turbo1 labeling for 30 minutes. Stress granule formation was then induced with 500 μM of sodium arsenite for 15 minutes or 1 hour. 15 minutes prior to APEX labeling, alkyne phenol was added. APEX labeling was then performed for 1 minute after the induction period.

For intercellular protein trafficking from cancer cell to macrophage cytosol, MC38 cells stably expressing TurboID-NES were mixed with Raw264.7-ASC cells stably expressing APEX2-NES (1:1 cell number) for 12 hours. TurboID labeling was initiated by adding a final concentration of 500 μM biotin for 1 hour, following by 4 hours of chase and 1-minute APEX labeling. To inhibit exosome biogenesis, the co-cultured cells were treated with 5 μM GW4869 for 12 hours. To inhibit nanotube formation, the co-cultured cells were treated with 10 μM L778123 for 12 hours.

For intercellular protein trafficking from macrophage ER to cancer cell surface, Raw264.7-ASC cells stably expressing TurboID-KDEL were treated with 100 ng/ml IFN-γ for 12 hours to drive M1-type polarization or 50 ng/mL IL-4 for 12 hours to drive M2-type polarization. Fo^γ^the medium transfer approach (Figure 7F), the cells were then treated with 500 μM biotin for 1 hour to initiate TurboID labeling, followed by 3.5 hours of chase. The medium was collected and supplemented with 500 μM alkyne-phenol, followed by the incubation with MC38 cells stably expressing HRP-TM. HRP-TM labeling was performed with 1 minute of 1 mM H_2_O_2_ treatment. The medium was removed and MC38 cells were subjected for further steps. For the co-culture system (Figure S9B), the macrophage cells were incubated with MC38 cells. TurboID labeling was initiated by adding a final concentration of 500 μM biotin for 1 hour, followed by 4 hours of chase and 1-minute HRP labeling.

#### TransitID tandem enrichment protocol

The cell pellets prepared as described above were lysed by resuspending in RIPA lysis buffer (50 mM Tris pH 8, 150 mM NaCl, 0.1% SDS, 0.5% sodium deoxycholate, 1% Triton X-100, 1× protease inhibitor cocktail (Sigma-Aldrich), and 1 mM PMSF) by gentle pipetting and incubating for 5 minutes at 4 °C. Lysates were clarified by centrifugation at 20,000 *g* for 10 min at 4 °C. Protein concentration in clarified lysate was estimated with Pierce BCA Protein Assay Kit (Thermo Fisher) and normalized to 2 mg/mL. 1 mL of lysates were reacted with 100 μM fluorescein azide, premixed 2-(4-((bis((1-tertbutyl-1H-1,2,3-triazol-4-yl)methyl)amino)methyl)-1H-1,2,3-triazol1-yl)-acetic acid (BTTAA)-CuSO_4_ complex (500 μM CuSO_4_, BTTAA:CuSO_4_ with a 2:1 molar ratio) and 2.5 mM freshly prepared sodium ascorbate for 2 hours at 37 °C. The resulting lysates were precipitated by 8 mL cold methanol at -80°C overnight and the precipitated proteins were centrifuged at 8000 *g* for 5 minutes at 4 °C. The proteins were washed twice with 1 mL of cold methanol and resuspended in 2 mL of NP-40 buffer (pH 7.5, 50 mM Tris-HCl, 150 mM NaCl, 1% NP-40 and 1% protease inhibitor) with sonication.

For anti-fluorescein IP, 250 μL of protein G dynabeads (Invitrogen) were washed with NP-40 buffer for twice and resuspended in 1 mL of NP-40 buffer. The beads were incubated with 30 μg of anti-fluorescein antibody (Abcam) at 30 °C with rotation for 1 hour. Then the beads were washed with NP-40 buffer twice. For IP, the beads were incubated with 2 mL of protein lysates at 4 °C with rotation overnight. After IP, the beads were washed with NP-40 buffer twice, high salt wash buffer (50 mM Tris-HCl pH 7.4, 1 M NaCl, 1 mM EDTA, 1% NP-40, 0.1% SDS, 0.5% sodium deoxycholate, in H_2_O) twice, finally with NP-40 buffer twice. For elution, the beads were resuspended in 50 μL of high-SDS RIPA buffer (50 mM Tris pH 8, 150 mM NaCl, 1% SDS, 0.5% sodium deoxycholate, 1% Triton X-100, 1× protease inhibitor cocktail, and 1 mM PMSF) and vortexed at 50 °C for 20 minutes. This elution step was repeated once and the eluates were combined. The beads were finally washed with 900 μL of SDS-free RIPA buffer (50 mM Tris pH 8, 150 mM NaCl, 0.5% sodium deoxycholate, 1% Triton X-100, 1× protease inhibitor cocktail, and 1 mM PMSF) and the supernatant was combined with the eluates to get a final SDS concentration of 0.1%.

To enrich biotinylated material from the anti-fluorescein IP eluates, 100 μL of streptavidin-coated magnetic beads (Pierce) were washed twice with RIPA buffer, then incubated with the 1 mL total eluates with rotation at 4 °C overnight. The beads were subsequently washed twice with 1 mL of RIPA lysis buffer, once with 1 mL of 1 M KCl, once with 1 mL of 0.1 M Na_2_CO_3_, once with 1 mL of 2 M urea in 10 mM Tris-HCl (pH 8.0), and twice with 1 mL of RIPA lysis buffer. For Western blotting analysis, the enriched proteins were eluted by boiling the beads in 75 μL of 3× protein loading buffer supplemented with 20 mM DTT and 2 mM biotin. For proteomic analysis, the beads were then resuspended in 1 mL of fresh RIPA lysis buffer and transferred to a new tube. The beads were then washed with 1 mL of washing buffer (75 mM NaCl in 50 mM Tris HCl pH 8.0) twice. The beads were resuspended in 50 μL of washing buffer and shipped to Steve Carr’s laboratory (Broad Institute) on dry ice for further processing and preparation for LC-MS/MS analysis.

#### Gels and Western blots

For all Western blots and silver staining gels, samples were resolved on a 10% SDS-PAGE gel. The silver-stained gels were generated using Pierce Silver Stain Kit (ThermoFisher Scientific). For all Western blots, after SDS-PAGE, the gels were transferred to a PVDF membrane, and then stained by Ponceau S (5 min in 0.1% (w/v) Ponceau S in 5% acetic acid/water). The blots were then blocked in 5% (w/v) BSA in TBS-T (Tris-buffered saline, 0.1% Tween 20) for at least 30 minutes at room temperature. For streptavidin blotting, the blots were stained with 0.3 μg/mL streptavidin-HRP in TBS-T for 1 hour at room temperature. For anti-fluorescein blotting, the blots were stained with anti-fluorescein-HRP (1:2500 dilution, ab19492, abcam) for 1 hour at room temperature. The blots were washed three times with TBS-T for 5 minutes each time before to development. For validation of the specificity of anti-fluorescein IP in Figure 1F, the blots were stained with primary antibodies in TBS-T for 2 hours in room temperature or overnight at 4°C. The primary antibodies include anti-GRSF1 (1:2500 dilution, ab205531, Abcam), anti-SDHA (1:2500 dilution, ab137040, Abcam), anti-MTCO1 (1:2500 dilution, ab203912, Abcam), anti-SYNJ2BP (1:2000 dilution, HPA000866-100UL, Sigma-Aldrich), anti-TOMM20 (1:1000 dilution, SC-17764, Santa Cruz Biotechnology) and anti-GAPDH-HRP (1:5000 dilution, SC-47724, Santa Cruz Biotechnology).

For validation of the specificity of TransitID in Figure 2D, the blots were stained with anti-GRSF1 (1:2500 dilution, ab205531, Abcam), anti-SDHA (1:2500 dilution, ab137040, Abcam), anti-MTCO1 (1:2500 dilution, ab203912, Abcam), anti-MTCO2 (1:2500 dilution, 55070-1-AP, Proteintech), anti-MTCO3 (1:2500 dilution, 55082-1-AP, Proteintech), anti-MTND1 (1:2000 dilution, 19703-1-AP, Proteintech), anti-MTND3 (1:1000 dilution, 45859, Cell Signaling Technology), anti-MTND4 (1:2000 dilution, HPA053928, Sigma Aldrich), anti-MTND4L (1:2000 dilution, PA5-68242, ThermoFisher Scientific), anti-MTND5 (1:2500 dilution, 55410-1-AP, Proteintech), anti-MTND6 (1:2000 dilution, PA5-109993, ThermoFisher Scientific), anti-MTATP6 (1:1000 dilution, 70262S, Cell Signaling Technology), anti-MTATP8 (1:2500 dilution, 26723-1-AP, Proteintech), anti-MTCYTB (1:2500 dilution, 55090-1-AP, Proteintech), anti-SYNJ2BP (1:2000 dilution, HPA000866-100UL, Sigma-Aldrich), anti-TOMM20 (1:1000 dilution, SC-17764, Santa Cruz Biotechnology) and anti-GAPDH-HRP (1:5000 dilution, SC-47724, Santa Cruz Biotechnology) in TBS-T overnight at 4°C.

For the validation of OMM-originated proteins in Figure 3I, the blots were stained with anti-MRPL30 (1:2500 dilution, ab179819, Abcam), anti-MRPL48 (1:2500 dilution, ab194826, Abcam), anti-HSP60 (1:2000 dilution, 4870S, Cell Signaling Technology), anti-EIF2AK2 (1:2000 dilution, 18244-1-AP, Proteintech), anti-SIRT5 (1:2500 dilution, ab241543, Abcam), anti-MTCO2 (1:2500 dilution, 55070-1-AP, Proteintech) and anti-Nucleolin (1:2500 dilution, ab129200, Abcam) in 3% BSA (w/v) in TBS-T for 2 h in room temperature or overnight at 4°C. For the validation of novel mitochondrial proteins in Figure S3G, the blots were stained with anti-EIF2AK2 (1:2000 dilution, 18244-1-AP, Proteintech), anti-SNAPC1 (1:2000 dilution, PA5-84742, ThermoFisher Scientific), anti-FBL (1:2500 dilution, ab166630, Abcam), anti-MTCO2 (1:2500 dilution, 55070-1-AP, Proteintech), anti-SDHA (1:2500 dilution, ab137040, Abcam) and anti-TOMM20 (1:1000 dilution, SC-17764, Santa Cruz Biotechnology) in TBS-T for 2 hours in room temperature or overnight at 4°C. For the validation of stress-inhibited translocated proteins in Figure 4D, the blots were stained with anti-ERC1 (1:2000 dilution, HPA019513, Sigma-Aldrich), anti-POLR2D (1:2000 dilution, HPA034694, Sigma-Aldrich), anti-ST13 (1:2000 dilution, HPA043233, Sigma-Aldrich), anti-MBD1 (1:1000 dilution, SC-25261, Santa Cruz Biotechnology) and anti-TOPBP1 (1:1000 dilution, SC-271043, Santa Cruz Biotechnology) in TBS-T for 2 hours in room temperature or overnight at 4°C. For the validation of TransitID for cytosol-to-nucleus translocation in Figure S4C, the blots were stained with anti-Nucleolin (1:2500 dilution, ab129200, Abcam), anti-Survivin (1:2500 dilution, ab76424, Abcam), anti-SYNJ2BP (1:2000 dilution, HPA000866-100UL, Sigma-Aldrich), anti-TOMM20 (1:1000 dilution, SC-17764, Santa Cruz Biotechnology) and anti-Calnexin (1:5000 dilution, PA5-34754, ThermoFisher Scientific) in TBS-T for 2 hours in room temperature or overnight at 4°C.

For blotting against JUN in Figure 6C-E, 6G, S8B, D, E, the blots were stained with anti-JUN (1:2000 dilution, HPA019513, Sigma-Aldrich) in TBS-T for 2 hours in room temperature or overnight at 4°C. For blotting against phosphorylated JUN in Figure S8E, the blots were stained with anti-phospho-JUN Ser63 (1:1000 dilution, 91952T, Cell Signaling Technology) and anti-phospho-JUN Ser73 (1:1000 dilution, 9164S, Cell Signaling Technology) in TBS-T for 2 hours in room temperature or overnight at 4°C. For blotting against G3BP1 in Figure S5A, S7C-D, the blots were stained with anti-G3BP1 (1:2500 dilution, 611126, BD Biosciences) in TBS-T for 2 hours in room temperature or overnight at 4°C. For blotting against DAXX in Figure 6C, S7C-D, the blots were stained with anti-DAXX (1:2500 dilution, HPA065779, Sigma-Aldrich) in TBS-T for 2 hours in room temperature or overnight at 4°C. For blotting against FOS in Figure 6C, 6E, the blots were stained with anti-FOS (1:2500 dilution, ab208942, Abcam) in TBS-T for 2 hours in room temperature or overnight at 4°C. For blotting against Tubulin in Figure 6C, 6E, S8D-E, the blots were stained with anti-α-Tubulin (1:5000 dilution, 3873, Cell Signaling Technology) in TBS-T for 2 hours in room temperat α or overnight at 4°C. For the validation of novel SG proteins in Figure S7C, the blots were stained with anti-MPP10 (1:2500 dilution, PA5-57134, ThermoFisher Scientific), anti-SRFBP1 (1:2500 dilution, HPA042737, Sigma-Aldrich) and anti-UBE2O (1:2500 dilution, PA5-54839, ThermoFisher Scientific) in TBS-T for 2 hours in room temperature or overnight at 4°C.

For the detection of cytosolic trafficking proteins from cancer cells to macrophages in Figure 7E, the blots were stained with anti-CDC42 (1:2500 dilution, ab187643, Abcam), anti-PTEN (1:2500 dilution, ab32199, Abcam), anti-VDAC1 (1:2500 dilution, ab14734, Abcam), anti-TOMM20 (1:1000 dilution, SC-17764, Santa Cruz Biotechnology), anti-Calnexin (1:5000 dilution, PA5-34754, ThermoFisher Scientific), anti-PDI (1:2000 dilution, SC-74551, Santa Cruz Biotechnology) and anti-GAPDH-HRP (1:5000 dilution, SC-47724, Santa Cruz Biotechnology) in 3% BSA (w/v) in TBS-T for 2 hours in room temperature or overnight at 4°C. For the detection of protein secretion from macrophages to cancer cell surface in Figure 7G and S9B, the blots were stained with anti-IL-1β (1:2500 dilution, M421B, ThermoFisher Scientific), anti-TNF-α (1:2500 dilution, ab215188, Abcam), anti-TGF-β (1:2500 dilution, ab179695, Abcam), anti-IL-10 (1:1000 dilution, MA5-42656, ThermoFisher Scientific) in 3% BSA (w/v) in TBS-T for 2 hours in room temperature or overnight at 4°C. To check the cross-labeling of PL enzymes in the co-culture system (Figure S9A), the blots were stained with anti-v5 (1:2500 dilution, R96025, ThermoFisher Scientific) and anti-Flag-HRP (1:2500 dilution, A8592, Sigma-Aldrich) in 3% BSA (w/v) in TBS-T for 2 hours in room temperature or overnight at 4°C.

After incubating with the primary antibody, the blots were washed with TBS-T for three times (5 minutes for each wash), then stained with secondary antibodies in 3% BSA (w/v) in TBS-T for 2 hours in room temperature. The blots were washed three times with TBS-T for 5 minutes each time before to development with Clarity Western ECL Blotting Substrates (Bio-Rad) and imaging on the ChemiDoc XRS+ System (Bio-Rad).

#### On-bead trypsin digestion of biotinylated proteins

Samples collected and enriched with streptavidin magnetic beads were washed twice with 200 μL of 50 mM Tris-HCl buffer (pH 7.5), transferred into new 1.5 mL Eppendorf tubes, and washed two more times with 200 μL of 50 mM Tris (pH 7.5) buffer. Samples were incubated in 0.4 μg trypsin in 80 μL of 2 M urea/50 mM Tris buffer with 1 mM DTT, for 1 hour at room temperature while shaking at 1000 rpm. Following pre-digestion, 80 μL of each supernatant was transferred into new tubes. Beads were then incubated in 80 uL of the same digestion buffer for 30 minutes while shaking at 1000 rpm. Supernatant was transferred to the tube containing the previous elution. Beads were washed twice with 60 μL of 2 M urea/50 mM Tris buffer, and these washes were combined with the supernatant. The eluates were spun down at 5000 × g for 30 seconds and the supernatant was transferred to a new tube. Samples were reduced with 4 mM DTT for 30 minutes at room temperature, with shaking. Following reduction, samples were alkylated with 10 mM iodoacetamide for 45 minutes in the dark at room temperature. An additional 0.5 μg of trypsin was added and samples were digested overnight at room temperature while shaking at 700 × g. Following overnight digestion, samples were acidified (pH < 3) with neat formic acid (FA), to a final concentration of 1% FA. Samples were spun down and desalted on C18 StageTips as previously described. Eluted peptides were dried to completion and stored at −80 °C.

#### TMTpro labeling and fractionation of peptides

Desalted peptides were labeled with TMTpro reagents (ThermoFisher Scientific). Peptides were resuspended in 80 μL of 50 mM HEPES and labeled with 20 μL 25 mg/mL TMTpro reagents in acetonitrile. Samples were incubated at room temperature for 1 hour with shaking at 1000 rpm. The TMTpro reaction was quenched with 4 μL of 5% hydroxylamine at room temperature for 15 minutes with shaking. TMTpro-labeled samples were combined, dried to completion, reconstituted in 100 μL of 0.1% FA, and desalted on StageTips. Combined TMTpro-labeled peptide sample was then desalted on C18 StageTips as previously described and dried to completion. For the cytosol-to-mitochondrial matrix translocation experiment and the cytosol-to-nucleus translocation experiments, 1/3 of the peptide sample was run as a singleshot and the remaining 2/3 was reserved for basic reverse phase (bRP) fractionation. For all other experiments, all of the peptide sample was bRP fractionated.

TMTpro labeled peptide sample was fractionated by bRP fractionation. StageTips packed with two disks of SDB-RPS (Empore) material. StageTips were conditioned with 100 μL of 100% MeOH, followed by 100 μL of 50% MeCN/0.1% FA and two washes with 100 μL of 0.1% FA. Peptide samples were resuspended in 200 μL of 1% FA (pH<3) and loaded onto StageTips. 6 step-wise elutions were carried out in 100 μL of 20 mM ammonium formate buffer with increasing concentration of 5%, 10%, 15%, 20%, 25%, and 45% MeCN. Eluted fractions were dried to completion.

#### Liquid chromatography and mass spectrometry

Peptide samples were separated with an online nanoflow Proxeon EASY-nLC 1200 UHPLC system (Thermo Fisher Scientific). The cytosol-to-mitochondrial matrix translocation peptide samples were analyzed on a Q-Exactive HFX mass spectrometer (Thermo Fisher Scientific) and all other samples were analyzed on an Orbitrap Exploris 480 mass spectrometer (Thermo Fisher Scientific). In this set up, the LC system, column, and platinum wire used to deliver electrospray source voltage were connected via a stainless-steel cross (360 mm, IDEX Health & Science, UH-906x). The column was heated to 50 °C using a column heater sleeve (Phoenix-ST). Each sample was injected onto an in-house packed 27 cm x 75 μm internal diameter C18 silica picofrit capillary column (1.9 mm ReproSil-Pur C18-AQ beads, Dr. Maisch GmbH, r119.aq; PicoFfrit 10 μm tip opening, New Objective, PF360-75-10-N-5).

For the experiments analyzed on the Orbitrap Exploris 480, mobile phase flow rate was 200 nL/min, comprised of 3% acetonitrile/0.1% formic acid (Solvent A) and 90% acetonitrile/0.1% formic acid (Solvent B). The 154-min LC– MS/MS method used the following gradient profile: (min:%B) 0:2;2:6; 122:35; 130:60; 133:90; 143:90; 144:50; 154:50 (the last two steps at 500 nL/min flow rate). Data acquisition was done in the data-dependent mode acquiring HCD MS/MS scans (r = 45,000) after each MS1 scan (r = 60,000) on the top 12 most abundant ions using a normalized MS1 AGC target of 100% and an MS2 AGC target of 50%. The maximum ion time utilized for MS/MS scans was 120 ms; the HCD-normalized collision energy was set to 32; the dynamic exclusion time was set to 20 s, and the peptide match and isotope exclusion functions were enabled. Charge exclusion was enabled for charge states that were unassigned, 1 and >6.

For the experiments analyzed on the Q-Exactive HFX, mobile phase flow rate was 200 nL/min, comprised of 3% acetonitrile/0.1% formic acid (Solvent A) and 90% acetonitrile/0.1% formic acid (Solvent B). bRP fractions were analyzed using a 110-min LC– MS/MS method used the following gradient profile: (min:%B) 0:2;1:6; 85:30; 94:60; 95:90; 100:90; 101:50; 110:50 (the last two steps at 500 nL/min flow rate). Data acquisition was done in the data-dependent mode acquiring HCD MS/MS scans (r = 45,000) after each MS1 scan (r = 60,000) on the top 12 most abundant ions using a normalized MS1 AGC target of 3e6 and an MS2 AGC target of 1e5. The maximum ion time utilized for MS/MS scans was 105 ms; the HCD-normalized collision energy was set to 29; the dynamic exclusion time was set to 15 s, and the peptide match and isotope exclusion functions were enabled. Charge exclusion was enabled for charge states that were unassigned, 1 and >6. Singleshot peptide sample was analyzed using a 154-min LC-MS/MS method with the same gradient profile as the method used on the Orbitrap Exploris 480. Data acquisition uses the same parameters as previously described for the 110 min method excepting that dynamic exclusion was set to 20 s.

#### Mass spectrometry data processing

Mass spectrometry data was processed using Spectrum Mill (proteomics.broadinstitute.org). For all samples, extraction of raw files retained spectra within a precursor mass range of 600-6000 Da and a minimum MS1 signal-to-noise ratio of 25. MS1 spectra within a retention time range of +/-60 seconds, or within a precursor m/z tolerance of +/-1.4 m/z were merged. MS/MS searching was performed against a human Uniprot database with a release date of December 28, 2017. Digestion parameters were set to “trypsin allow P” with an allowance of 4 missed cleavages. The MS/MS search included fixed modification of carbamidomethylation on cysteine. TMTpro was searched using the full-mix function. Variable modifications were acetylation and oxidation of methionine. Restrictions for matching included a minimum matched peak intensity of 30% and a precursor and product mass tolerance of +/-20 ppm. Peptide spectrum matches (PSMs) were validated using a maximum false discovery rate (FDR) threshold of 1.2% for precursor charges 2 through 6 within each LC-MS/MS run. Protein polishing autovalidation was further applied to filter the PSMs using a target protein score threshold of 0. TMTpro reporter ion intensities were corrected for isotopic impurities in the Spectrum Mill protein/peptide summary module using the afRICA correction method which implements determinant calculations according to Cramer’s Rule. We required fully quantified unique human peptides with 2 unique peptides and 2 ratio counts for protein quantification. We used the Proteomics Toolset for Integrative Data Analysis (Protigy, v1.0.4, Broad Institute, https://github.com/broadinstitute/protigy) to calculate moderated t-test P values for regulated proteins.

#### Analysis of proteomic data for cytosol to mitochondrial matrix translocation

The original identified proteins are shown in Table S1. To determine the cutoff ratio for each comparison, a receiver operating characteristic (ROC) analysis was performed. The true-positives (TPs) were known mitochondrial matrix proteins annotated by GO: 0005759; and not known as mtDNA-encoded proteins or not annotated with the following GO terms: GO:0005741 for OMM, GO:0005758 for IMS, GO:0005743 for IMM. The false-positives (FPs) were plasma membrane proteins annotated by GO:0005886. For each comparison, the proteins were first ranked in a descending order according to the mean of fold change. For each protein on the ranked list, the accumulated true-positive count and false-positive count above its fold change ratio were calculated. A receiver operating characteristic (ROC) curve was plotted accordingly for each comparison (Figure S2C). The cutoff was set where true-positive rate – false-positive rate (TPR-FPR) maximized. After further filtering with p values (*p* < 0.05) (Figure S2D), post-cutoff proteomic lists of all comparisons were intersected to obtain the final cytosol-to-mitochondrial matrix and OMM-to-mitochondrial matrix translocated protein list, respectively (Figure S2B, Table S1).

For the analysis of mitochondria specificity of the translocated proteins (Figure S3B), a list of mitochondrial proteins was collected from the MitoCarta 3.0 database, the MitoCarta 2.0 database, GOCC terms containing mitochondrial annotations, the mitochondrial matrix proteome identified by APEX profiling^16^. The number and percentage of mitochondrial proteins in human proteome, cytosol-to-mitochondrial matrix translocated proteome and OMM-to-mitochondrial matrix translocated proteome was determined. For the analysis of sub-mitochondria specificity of the translocated proteins (Figure S3C), the number and percentage of proteins for each submitochondrial compartment was determined in human proteome, cytosol-to-mitochondrial matrix translocated proteome and OMM-to-mitochondrial matrix translocated proteome. To analyze the depth of coverage (Figure S3E), cytosol-to-mitochondrial matrix and OMM-to-mitochondrial matrix translocated proteins were combined and crossed with five groups of well-established mitochondrial matrix proteins (i–v). The combined translocated proteins were also compared against the mitochondrial matrix proteome identified by APEX profiling^16^. The translocated proteins covered or not covered by the single APEX-mito profiling were subjected to the analysis of protein half-lives according to a previous study^17^.

To assign mitochondrial matrix proteins that specifically originate from the OMM, the enrichment of TurboID-OMM&APEX2-mito TransitID labeling was compared to the TurboID-NES&APEX2-mito TransitID labeling. The translocated proteins that are significantly more enriched by the TurboID-OMM TransitID labeling (fold change > 1, *p* < 0.05) are considered as OMM-locally translated proteins (Figure 3D). The mitochondrial uptake rate for OMM-locally translated proteins were obtained from a previous study^65^.

#### Analysis of proteomic data for cytosol to nucleus translocation

The original identified proteins are shown in Table S2. To determine the cutoff ratio for each comparison, a receiver operating characteristic (ROC) analysis was performed. The true-positives (TPs) were gold standard nuclear proteins assembled in our previous study^7^. The false-positives (FPs) were known mitochondrial matrix proteins annotated by GO: 0005759 and not annotated with the following GO terms: GO:0005741 for OMM, GO:0005758 for IMS, GO:0005743 for IMM. For each comparison, the proteins were first ranked in a descending order according to the mean of fold change. For each protein on the ranked list, the accumulated true-positive count and false-positive count above its fold change ratio were calculated. A receiver operating characteristic (ROC) curve was plotted accordingly for each comparison (Figure S4D). The cutoff was set where true-positive rate – false-positive rate (TPR-FPR) maximized. After further filtering with p values (*p* < 0.05), post-cutoff proteomic lists of all comparisons were intersected to obtain the final cytosol-to-nucleus translocated protein list (Figure S4F, Table S2). To evaluate the impact of arsenite on cytosol-to-nucleus translocation, the TurboID-NES&APEX2-NLS TransitID labeling under arsenite was compared to the TransitID labeling under basal condition (Figure 4C).

#### Analysis of proteomic data for SG-nucleolus/nucleus communications

The original identified proteins for nucleolus/nucleus-to-SG translocation during stress induction are shown in Table S3. For comparison of TransitID labeling against -light or -H_2_O_2_ negative controls, true positive proteins (TP1) were known nucleolar proteins annotated by GO:0005730 and false-positives (FP1) were known mitochondrial matrix proteins annotated by GO: 0005759 and not annotated with the following GO terms: GO:0005741 for OMM, GO:0005758 for IMS, GO:0005743 for IMM. For each comparison, the proteins were first ranked in a descending order according to the mean of fold change. For each protein on the ranked list, the accumulated true-positive count and false-positive count above its fold change ratio were calculated. A receiver operating characteristic (ROC) curve was plotted accordingly for each comparison (Figure S5C). The cutoff was set where true-positive rate – false-positive rate (TPR-FPR) maximized. After further filtering with p values (*p* < 0.05) (Figure S2D), post-cutoff proteomic lists of both comparisons were intersected to obtain the 553 dual labeled proteins (Figure S5B, Table S3). For comparison of TransitID labeling under arsenite against dual labeling under basal condition, 109 dual labeled proteins with log_2_Fold change > 0.5 and p value < 0.05 were assigned as stress-dependent translocated proteins (Figure S5B, Table S3). For comparison of LOV-TurboID-NIK3x&APEX2-G3BP1 TransitID labeling vs LOV-TurboID-NIK3x&APEX2-NES TransitID labeling, true positive proteins (TP2) were known stress granule proteins annotated by GO:0010494 and false-positives (FP2) were known OMM proteins assembled in our previous study^68^. After filtering with the ROC cutoff and p value, the post-cutoff list of this comparison was crossed with the stress-dependent translocated proteins to obtain the 73 stress-dependent translocated proteins specifically to stress granules (Figure S5B, Table S3). For comparison of LOV-TurboID-NIK3x&APEX2-G3BP1 TransitID labeling vs LOV-TurboID-NLS&APEX2-G3BP1 TransitID labeling, true positive proteins (TP3) were known nucleolar proteins annotated by GO:0005730 and false-positives (FP3) were known nuclear pore proteins annotated by GO:0005643. After filtering with the ROC cutoff and p value, the post-cutoff list of this comparison was crossed with the 73 stress-dependent translocated proteins specifically to stress granules to obtain the 36 nucleolus-to-SG and 37 nucleus-to-SG translocated proteins during stress induction (Figure S5B, Table S3). To differentiate fast translocated proteins from slower ones (Figure 5J), TransitID enrichment with 15-minute arsenite treatment was compared to 1-hour treatment for the 73 translocated proteins.

The original identified proteins for SG-to-nucleolus/nucleus translocation during stress recovery are shown in Table S4. For comparison of TransitID labeling against -light or -H_2_O_2_ negative controls, true positive proteins (TP1) were known stress granule proteins annotated by GO:0010494 and false-positives (FP1) were known mitochondrial matrix proteins annotated by GO: 0005759 and not annotated with the following GO terms: GO:0005741 for OMM, GO:0005758 for IMS, GO:0005743 for IMM. For each comparison, the proteins were first ranked in a descending order according to the mean of fold change. For each protein on the ranked list, the accumulated true-positive count and false-positive count above its fold change ratio were calculated. A receiver operating characteristic (ROC) curve was plotted accordingly for each comparison (Figure S5B). The cutoff was set where true-positive rate – false-positive rate (TPR-FPR) maximized. After further filtering with p values (*p* < 0.05) (Figure S2D), post-cutoff proteomic lists of both comparisons were intersected to obtain the 549 dual labeled proteins (Figure S5A, Table S4). For comparison of TransitID labeling under arsenite against TransitID labeling under basal condition, 29 dual labeled proteins with log_2_Fold change > 0.5 and p value < 0.05 were assigned as stress-dependent translocated proteins (Figure S5A, Table S4). For comparison of LOV-TurboID-G3BP1&APEX2-NIK3x TransitID labeling vs LOV-TurboID-NES&APEX2-NIK3x TransitID labeling, true positive proteins (TP2) were known stress granule proteins annotated by GO:0010494 and false-positives (FP2) were known OMM proteins assembled in our previous study^68^. After filtering with the ROC cutoff and p value, the post-cutoff list of this comparison was crossed with the stress-dependent translocated proteins to obtain the 21 stress-dependent translocated proteins specifically from stress granules (Figure S5A, Table S4). For comparison of LOV-TurboID-G3BP1&APEX2-NIK3x dual labeling vs LOV-TurboID-G3BP1&APEX2-NLS TransitID labeling, true positive proteins (TP3) were known nucleolar proteins annotated by GO:0005730 and false-positives (FP3) were known nuclear envelop proteins annotated by GO:0005635. After filtering with the ROC cutoff and p value, the post-cutoff list of this comparison was crossed with the 21 stress-dependent translocated proteins specifically from stress granules to obtain the 20 nucleolus-to-SG and 1 nucleus-to-SG translocated protein during stress recovery (Figure S5A, Table S4). To differentiate fast translocated proteins from slower ones (Figure 5K), TransitID enrichment with 1-hour recovery was compared to 3-hour recovery for the 21 translocated proteins.

For the analysis of stress granule specificity (Figure S6D), the final lists were crossed with a combined list of known stress granule proteins assembled in a previous study^69^. For the analysis of nuclear specificity (Figure S6E), a list of known nuclear proteins were collected from human proteins annotated with annotated with the following Gene Ontology terms: GO:0016604, GO:0031965, GO:0016607, GO:0005730, GO:0001650, GO:0005654, GO:0005634. The number and percentage of known nuclear proteins in human proteome and different translocated proteomes was determined. For the analysis of nucleolar specificity (Figure S6F), the final lists were crossed with known nucleolar proteins annotated by GO:0005730 or proteins labeled by at least two nucleolus-localized BioID baits in a previous study^8^. For the analysis of RNA binding specificity (Figure S6G), the final lists were crossed with known RNA-binding proteins assembled in RBPbase (https://rbpbase.shiny.embl.de/). For the analysis of phase separation propensity of those translocated proteins (Figure 6H), Pscore values of translocated proteins in the final lists were compared to Pscore values of the human proteome according to a previous study^32^. For the analysis of the percentage of intrinsic disorder regions (IDRs) in those translocated proteins (Figure 6I), the IDR percentage of translocated proteins in the final lists were compared to the IDR percentage of the human proteome according to a previous study^33^.

#### Validation of novel mitochondrial proteins by mitochondrial fractionation

To validate novel mitochondrial proteins in the final cytosol-to-mitochondrial matrix and OMM-to-mitochondrial matrix translocated protein lists, mitochondria isolation was performed, followed by the proteinase K treatment. Mitochondria isolation was performed according to the manufacturer’s protocol (ThermoFischer Scientific). Briefly, 2e^7^ HEK 293T cells were resuspended in 800 μL of mitochondria isolation reagent A and homogenized in Dounce Tissue Grinder on ice. 800 μL of Mitochondria Isolation Reagent C and 200 μL of Mitochondria Isolation Reagent A were added into the lysates. After mixing, the lysates were centrifuged at 700 *g* for 10 minutes at 4°C and the supernatant was transferred to a new tube and centrifuged at 3,000 *g* for 15 minutes at 4°C. The supernatant was saved as the cytosolic fraction. The pellets were resuspended in 500 μL of Mitochondria Isolation Reagent C, followed by centrifugation at 12,000 *g* for 5 minutes. The supernatant was discarded, and the pellets are the mitochondrial fraction. The pellets were resuspended in 100 μL PBS buffer containing different concentrations of proteinase K for 15 minutes at 4°C. The incubation was stopped by adding 7 mM PMSF and subjected to Western blot analysis.

#### Capture of localized nascent polypeptides

To capture OMM-localized newly synthesized polypeptides (Figure 3G-I), APEX2-OMM stable cells were treated with 15 μM of O-propargyl-puromycin (OPP) for 30 minutes. Then the medium was replaced with that containing 15 μM of O-propargyl-puromycin (OPP) and 500 μM of biotin-phenol for another 30 minutes. H_2_O_2_ was then added to a final concentration of 1 mM and the plate was gently agitated for 1 minute. The reaction was quenched by replacing the medium with an equal volume of quenching solution (10 mM ascorbate, 5 mM Trolox and 10 mM sodium azide in DPBS). Cells were washed with quenching solution for three times. Cells were washed twice with 10 mL ice-cold DPBS, harvested by scraping, pelleted by centrifugation at 1,400 rpm for 3 minutes, and either processed immediately or flash frozen in liquid nitrogen and stored at -80 °C before further analysis. To enrich biotinylated newly synthesized polypeptides, tandem enrichment was performed as described above. The enriched materials were subjected to analysis by silver staining and Western blot.

#### TurboID-NES labeling following by nuclear fractionation

In order to validate the impact of arsenite on cytosol-to-nucleus translocation (Figure 4D), TurboID-NES stable cells cultured in T75 flasks were treated with 50 μM of biotin for 10 minutes. After removing biotin-containing medium, the cells were washed with ice-cold DPBS for three times. Then the cells were cultured for another 2 hours with or without 500 μM sodium arsenite treatment. Cells were washed with 10 mL ice-cold DPBS for three times, harvested by scraping, pelleted by centrifugation at 1,400 rpm for 3 minutes.

Nuclear fractionations were performed following the protocols and suggestions published by a previous study^70^. Cells were collected in DPBS with some aliquoted for whole cell lysate samples then pelleted at 500 *g* at 4 °C for 2 minutes. 1 mL of hypotonic buffer (HLB: 20 mM Tris (pH7.5), 5 mM KCl, 3 mM MgCl_2_, 10% glycerol, 0.5% NP-40 and protease inhibitor cocktail) was used to resuspend 75 mg of cells and incubated on ice for 10 minutes. Cells were briefly vortexed and centrifuged at 500 *g* for 5 minutes at 4°C. 870 μL of the supernatant was transferred to a new tube and combined with 25 μL of 5 M NaCl to generate the cytoplasmic fraction. The remaining supernatant was discarded and the pellet was resuspended in 1 mL HLB. The nuclear fraction was pelleted at 500 *g* for 2 minutes then washed in cold isotonic wash buffer (IWB: 20 mM Tris-HCl (pH 7.5), 100 mM KCl, 3 mM MgCl_2_, 10% glycerol, 0.6% NP40, 0.5 mM DTT (DTT should be freshly added). This washing step with the IWB was performed once more. Then, the nuclear pellet was lysed in 1 mL RIPA buffer containing benzonase at 1,000 U/mL (Millipore).

To enrich biotinylated proteins from the nuclear fraction, 100 μL streptavidin-coated magnetic beads (Pierce) were washed twice with RIPA buffer, then incubated with the 1 mL nuclear fraction with rotation at 4 °C overnight. The beads were subsequently washed twice with 1 mL of RIPA lysis buffer, once with 1 mL of 1 M KCl, once with 1 mL of 0.1 M Na_2_CO_3_, once with 1 mL of 2 M urea in 10 mM Tris-HCl (pH 8.0), and twice with 1 mL of RIPA lysis buffer. The enriched proteins were eluted by boiling the beads in 75 μL of 3× protein loading buffer supplemented with 20 mM DTT and 2 mM biotin, following by Western blot analysis using antibodies targeting protein of interest.

#### Immunofluorescence staining and fluorescence microscopy

To verify novel stress granule proteins in Figure 4F, 6A and Supplementary Figure 7A, HEK 293T cells were plated on human fibronectin (EMD Millipore) coated glass coverslips. Cells were then treated with or without 500 μM sodium arsenite for 1 hour then fixed for 10 minutes with 4% paraformaldehyde at room temperature. Cells were then permeabilized with cold methanol for 10 minutes at -20 °C and blocked in 5% BSA dissolved in DPBS for 1 hour. The coverslips were incubated with primary antibodies against a stress granule marker of either G3BP1 or FXR1 and candidate proteins ERC1, POLR2D, ST13, JUN, DAXX, MPP10, SRFBP1, or UBE2O for 2 hours at room temperature. Coverslips were washed three times in DPBS with 5 minutes for each washed then incubated in secondary antibodies conjugated to either AlexaFluor-488, 568, or 647 and DAPI for 1 hour. Coverslips were washed thrice more then mounted on glass slides.

To verify stress granule localization from heat stress in Supplementary Figure 7B, U2OS cells were plated on human fibronectin coated glass coverslips and treated with or without heat shock at 45 °C for 2 hours. Cells were then fixed, permeabilize, and blocked, then incubated with primary antibodies for 2 hours followed by secondary antibodies for 1 hour and mounted onto slides.

To verify the localization of APEX2-mito in Figure 1C, stable HEK 293T cells expressing APEX2-mito-V5 on human fibronectin coated coverslips were used for APEX labeling with alkyne phenol and 1 minute hydrogen peroxide. Cells were fixed and permeabilized. Alkyne labeled proteins were then reacted with 100 μM azide-AlexaFluor488 in a copper catalyzed click reaction containing premixed 2-(4-((bis((1-tertbutyl-1H-1,2,3-triazol-4-yl)methyl)amino)methyl)-1H-1,2,3-triazol1-yl)-acetic acid (BTTAA)-CuSO_4_ complex (500 μM CuSO4, BTTAA:CuSO_4_ with a 2:1 molar ratio) and 2.5 mM freshly prepared sodium ascorbate for 2 hours at room temperature. Coverslips were then blocked for, incubated with primary antibodies against TOMM20 and V5, washed three times with DPBS, incubated with secondary antibodies, washed, and mounted on glass slides.

To verify dual labeling of cytosol and outer mitochondrial membrane to mitochondrial matrix translocation in Figure 3B, stable HEK293T cells expressing APEX2-mito-V5 were transfected with TurboID-NES-flag or TurboID-OMM-flag. Cells were induced to label with 50 μM biotin for 1 hour followed by a 24-hour chase. Alkyne phenol was added to the media 30 minutes prior to the end of the chase period. Cells were then labeled with 1 minute of hydrogen peroxide treatment. Cells were then fixed and permeabilized. Alkyne labeled proteins were then reacted with 100 μM azide-AlexaFluor405 by click reaction described above. Coverslips were then blocked for 1 hour, incubated with primary antibodies against the Flag epitope directly conjugate to phycoerythrin and V5 epitope directly conjugated to AlexFluore647 as well as neutravidin conjugated to AlexaFluor488. Coverslips were washed three times with DPBS and mounted on glass slides.

To verify dual labeling of cytosol to nucleus translocation under basal or sodium arsenite stress in Figure 4B, stable HEK 293T cells expressing APEX2-NLS-V5 were transfected with TurboID-NES-flag. Cells were induced to label with 50 μM biotin for 10 minutes followed by a 2-hour chase with or without 500 μM sodium arsenite. Alkyne phenol was added to the media 30 minutes prior to the end of the chase period. Cells were then labeled with 1 minute of hydrogen peroxide treatment. Cells were then fixed and permeabilized. Alkyne labeled proteins were then reacted with 100 μM azide-AlexaFluor405 by click reaction described above. Coverslips were then blocked for 1 hour, incubated with primary antibodies against the Flag epitope directly conjugate to phycoerythrin and V5 epitope directly conjugated to AlexFluore647 as well as neutravidin conjugated to AlexaFluor488. Coverslips were washed three times with DPBS and mounted on glass slides.

To verify dual labeling of nucleolus to stress granule translocation in Figure 5C, G3BP1 knocked-out HEK 293T cells were transfected with APEX-V5-G3BP1 and flag-LOV-Turbo1-NIK3x. Cells were induced to label with 100 μM biotin and blue light for 30 minutes followed by a 1-hour chase with or without 500 μM sodium arsenite. Alkyne phenol was added to the media 15 minutes prior to the end of the chase period. Cells were then labeled with 1 minute of hydrogen peroxide treatment. Cells were then fixed and permeabilized. Alkyne labeled proteins were then reacted with 100 μM azide-AlexaFluor405 by click reaction described above. Coverslips were then blocked for 1 hour, incubated with primary antibodies against the Flag epitope for 2 hours, washed three times with DPBS, incubated with anti-mouse-AlexaFluor568 secondary antibody for 1 hour, washed three times with DPBS, incubated with anti-V5-AlexaFluor647 and neutravidin-AlexaFluor488 for 2 hours, washed three times with DPBS, and mounted on glass slides.

To verify dual labeling of stress granule to nucleolus translocation during stress recovery, G3BP1 knocked-out HEK293T cells were transfected with GFP-APEX2-NIK3x and V5-LOV-Turbo1-G3BP1. Cells were stressed with 500 μM sodium arsenite for 1 hour or kept at basal condition, then induced to label with 100 μM biotin and blue light for 30 minutes in media without sodium arsenite, followed by a 2.5-hour chase. Alkyne phenol was added to the media 15 minutes prior to the end of the chase period. Cells were then labeled with 1 minute of hydrogen peroxide treatment. Cells were then fixed and permeabilized. Alkyne labeled proteins were then reacted with 100 μM azide-AlexaFluor405 by click reaction described above. Coverslips were then blocked for 1 hour, incubated with anti-V5-AlexaFluor647 and neutravidin-488 for 2 hours, washed three times with DPBS, and mounted on glass slides.

To verify dual labeling of MC38 colon cancer cells to Raw264.7 macrophages in Figure 7B, stable cells of MC38 expressing flag-GFP-TurboID-NES and Raw264.7 expressing V5-GFP-APEX2-NES were plated in equal ratio on human fibronectin coated coverslips. Cells were induced to label with 50 μM biotin for 1 hour followed by a 4-hour chase. Alkyne phenol was added to the media 30 minutes prior to the end of the chase period. Cells were then labeled with 1 minute of hydrogen peroxide treatment. Cells were then fixed and permeabilized. Coverslips were then blocked for 1 hour, incubated with primary antibody against the V5 epitope for 2 hours, washed three times with DPBS, incubated with anti-mouse-AlexaFluor405 secondary antibody for 1 hour, washed three times with DPBS, incubated with anti-Flag-phycoerythrin and neutravidin-AlexaFluor647 for 2 hours, washed three times with DPBS, and mounted on glass slides.

Fluorescence confocal microscopy was performed with a Zeiss AxioObserver microscope with 60× oil immersion objectives, outfitted with a Yokogawa spinning disk confocal head, Cascade II:512 camera, a Quad-band notch dichroic mirror (405/488/568/647), and 405 (diode), 491 (DPSS), 561 (DPSS) and 640 nm (diode) lasers (all 50 mW). DAPI (405 laser excitation, 445/40 emission), Alexa Fluor488 (491 laser excitation, 528/38 emission) and AlexaFluor647 (640 laser excitation, 700/75 emission) and differential interference contrast (DIC) images were acquired through a 60x oil-immersion lens. Acquisition times ranged from 100 to 2,000 ms. All images were collected and processed using SlideBook 6.0 software (Intelligent Imaging Innovations).

#### Photoactivation

HeLa cells were seeded into a four-well Lab-tek chambered cover glass (Nunc) cultured with DMEM supplemented with 10% FBS. The day after seeding, the cells were co-transfected with paGFP-JUN and mCherry-PABP constructs using ViaFect (Promega) for 24 hours. Prior to imaging, media was replaced with Fluorobrite DMEM media supplemented with 10% FBS and 4 mM L-glutamine.

Photoactivation experiments were performed on a Yokogawa CSU W1 spinning disk attached to a Nikon Ti2 eclipse with a Photometrics Prime 95B camera using Nikon Elements software (version 5.21.02). The light path was split between the port for the spinning disk/acquisition laser and the Photostimulation lasers, enabling Photoactivation with the 405nm laser to occur simultaneously while imaging. All Photoactivation imaging was taken on a Nikon Plan Apo 60× 1.40 NA oil objective, with Immersol 518 F/37C (Zeiss; refractive index 1.518) with Perfect Focus 2.0 engaged. During imaging, cells were maintained at 37°C and supplied with 5% CO_2_ using a Bold Line Cage Incubator (Okolabs) and an objective heater (Bioptechs). To induce stress granules, cells were incubated with 500 µM sodium arsenite for 60 min, and then was washed out to recover from stress. Photoactivation with the 405nm photostimulation laser in the indicated location occurred over 100 ms once during a 10 s continuous capture where 488nm and 561nm channels were imaged every 200 ms. After each photoactivation acquisition, the xy stage position was recorded for each position. Either after arsenite addition or 15 minutes after arsenite washout, each recorded location was imaged every 2 minutes until completion of the experiment.

#### Co-immunoprecipitation

To validate the novel SG proteins identified by TransitID (Figure S7C and S8A), immunoprecipitation of G3BP1 was performed by using a G3BP1 antibody (BD Biosciences). HEK 293T cells were exposed to 500 μM sodium arsenite for 1 hour at 37°C. After treatment, cells were washed three times with DPBS and lysed in 500 μL of NP-40 buffer (pH 7.5, 50 mM Tris-HCl, 150 mM NaCl, 1% NP-40 and 1% protease inhibitor) with sonication. The protein concentration was determined by BCA and normalized to 1 mg/mL. 500 μL of lysates were treated with 4 U/ml RNase I (ThermoFischer Scientific) or 1 M NaCl for 15 minutes at 4°C. Then the lysates were incubated with 125 μL protein G dynabeads (pre-incubated with 15 μg anti-G3BP1 antibody) at 4 °C with rotation overnight. The beads were then washed with NP-40 buffer for three times and boiled in protein loading buffer for Western blot analysis with antibodies indicated.

To evaluate the interaction between JUN and FOS (Figure 6E), immunoprecipitation of FOS was performed by using a FOS antibody (Abcam). For “stress” samples, HEK 293T cells or G3BP1&2 double knockout (DKO) cells were treated with 500 μM sodium arsenite for 1 hour at 37°C. For “recovery” samples, HEK 293T cells or DKO cells were firstly treated with 500 μM sodium arsenite for 1 hour at 37°C, and then cultured in normal medium for 1 hour after washing the cells with DPBS for three times. The cells were washed three times with PBS and lysed in 500 μL NP-40 buffer (pH 7.5, 50 mM Tris-HCl, 150 mM NaCl, 1% NP-40 and 1% protease inhibitor) with sonication. The protein concentration was determined by BCA and normalized to 1 mg/mL. 500 μL lysates were incubated with 125 μL protein G dynabeads (pre-incubated with 15 μg anti-FOS antibody) at 4 °C with rotation overnight. The beads were then washed with NP-40 buffer for three times and boiled in protein loading buffer for Western blot analysis using anti-JUN antibody (Sigma Aldrich).

#### Protein solubility assay

To evaluate the impact of stress granules on the solubility of JUN (Figure 6C), HEK 293T cells or DKO cells were treated with 500 μM sodium arsenite for 1 hour at 37°C. For “recovery” samples, HEK 293T cells or DKO cells were firstly treated with 500 μM sodium arsenite for 1 hour at 37°C, and then cultured in normal medium for 1 hour after washing the cells with PBS for three times. The cells were washed three times with PBS and lysed in cold DPBS containing 1% protease inhibitor with sonication. The cell lysates were centrifuged at 20,000 *g* for 30 minutes at 4 °C and the supernatant was collected as the soluble fraction. The pellets were resuspended in 4% SDS/PBS and centrifuged at 20,000 *g* for 30 minutes at 4 °C. The supernatant was discarded and the pellets were resuspended in protein loading buffer with boiling as the insoluble fraction. The soluble and insoluble fractions were subjected to Western blot analysis.

#### Thermal shift assay

To evaluate the impact of DAXX on the thermal stability of G3BP1 (Figure S7D), 1 μM of human recombinant G3BP1 protein (Abcam) was incubated with 0.5 μM of human recombinant DAXX protein (Abcam) for 30 minutes at 37 °C. The samples were divided into 6 aliquots and transferred into 0.2-mL polymerase chain reaction (PCR) tubes. Each sample was heated in parallel for 3 minutes to the respective temperature (range: 37–62 °C). Subsequently, the samples were centrifuged at 20,000 g for 20 minutes at 4°C and supernatant was collected for Western blot analysis using anti-G3BP1 and anti-DAXX antibodies.

#### Quantification of protein degradation by Azidohomoalanine pulse chase labeling

To evaluate the impact of stress granules on the degradation of JUN (Figure 6D), HEK 293T cells or DKO cells were cultured in medium containing 1 mM Azidohomoalanine (AHA) for 24 hours at 37°C. Then the cells were washed with DPBS for three times and chased into the normal medium with different treatments. For “stress” samples, the cells were treated with 500 μM sodium arsenite during the 1-hour chase and a control sample is 1-hour chase under basal condition. For “recovery” samples, the cells were treated with 500 μM sodium arsenite for 1 hour and then cultured in normal medium for 3 hours after washing the cells with PBS for three times. A control sample is 4-hour chase under basal condition. Cells were washed with 10 mL ice-cold DPBS for three times, harvested by scraping, pelleted by centrifugation at 1,400 rpm for 3 minutes, and either processed immediately or flash frozen in liquid nitrogen and stored at -80 °C before further analysis.

Cells were lysed in RIPA buffer and protein concentration was normalized to 2 mg/mL. 1 mL lysates were reacted with 100 μM biotin-PEG_4_-alkyne, premixed BTTAA-CuSO_4_ complex (500 μM CuSO_4_, BTTAA:CuSO_4_ with a 2:1 molar ratio) and 2.5 mM freshly prepared sodium ascorbate for 2 hours at room temperature. The resulting lysates were precipitated by 8 mL methanol at -80°C overnight and the precipitated proteins were centrifuged at 8,000 *g* for 5 minutes at 4 °C. The proteins were washed twice with 1 mL cold methanol and resuspended in 1 mL RIPA buffer with sonication. The biotinylated proteins were further captured by 200 μL streptavidin magnetic beads for 2 hours. The beads were washed as described above and proteins were eluted by boiling the beads in 75 μL of 3× protein loading buffer supplemented with 20 mM DTT and 2 mM biotin. The resulting samples were analyzed by Western bloting with anti-JUN antibody.

#### Quantification of protein synthesis by Azidohomoalanine labeling

To evaluate the impact of stress granules on protein synthesis of JUN (Figure S7C), cells were cultured in methionine-free medium supplemented with 1 mM azidohomoalanine (AHA) along with different treatments. For “stress” samples, the cells were treated with 500 μM sodium arsenite during the 1 hour of AHA labeling and a control sample is 1-hour labeling under basal condition. For “recovery” samples, the cells were treated with 500 μM sodium arsenite for 1 hour and then labeled under basal condition for 3 hours after washing the cells with DPBS for three times. A control sample is 4-hour labeling under basal condition. Cells were washed with 10 mL of ice-cold DPBS for three times, harvested by scraping, pelleted by centrifugation at 1,400 rpm for 3 minutes, and either processed immediately or flash frozen in liquid nitrogen and stored at -80 °C before further analysis. The sample processing including click reaction and streptavidin enrichment was carried out as described above.

#### JUN transcriptional activity assay

To evaluate the impact of stress granules on the transcription activity of JUN by genetic knockout (Figure 6F), HEK 293T cells or DKO cells were treated with 500 μM sodium arsenite for 1 hour at 37°C. For “recovery” samples, HEK 293T cells or DKO cells were firstly treated with 500 μM sodium arsenite for 1 hour at 37°C, and then cultured in normal medium for 3 hours after washing the cells with DPBS for three times. To evaluate the impact of stress granules on the transcription activity of JUN by small molecule inhibition (Figure S6B), HEK 293T cells were treated with 1 μM of GSK2606414 for 30 minutes and then treated with 500 μM of sodium arsenite and 1 μM of GSK2606414 for 1 hour. For “recovery” samples, the cells were cultured in normal medium for 3 hours after washing the cells with DPBS for three times.

JUN transcription activity was determined using a commercial JUN transcription factor activity assay kit (RayBiotech) by following the manufacturer’s protocol. The cells were lysed in cold PBS containing 1% protease inhibitor with sonication. The cell lysates were centrifuged at 20,0300 g for 30 minutes at 4 °C and the supernatant was collected. The cell lysates were incubated with the 96-well plate coated with double strand DNA containing a JUN binding sequence for 2 hours at room temperature with gentle shaking. After the binding, the lysates were removed and the plates were washed with 1x wash buffer for four times. Then each well was incubated with 100 μL of prepared TF Activity Assay c-JUN Primary Antibody for 1 hour at room temperature with gentle shaking. After four times of washes, each well was further incubated with 100 μL of prepared TF Activity Assay HRP-conjugated Secondary Antibody for 1 hour at room 3 with gentle shaking. After four times of washes, each well was further incubated with 100 μL of TMB One-Step Substrate Reagent for 30 minutes at room temperature in the dark with gentle shaking. The reaction was stopped by adding 50 μL of stop solution to each well and the plate was read at 450 nm immediately to quantify relative JUN transcription activity.

#### Cell proliferation assays

In order to determine the cellular toxicity of alkyne-phenol (Figure S1D), 1E4 HEK 293T cells per well were plated in 96-well plates with 100 μL fresh medium per well and cultured for 24 hours. Then the cells were treated with 50 or 500 μM of alkyne-phenol or biotin-phenol for 30 minutes at 37°C. In order to determine the effect of stress-sensitive (ERC1, POLR2D and ST13) and stress-insensitive (MBD1 and TOPBP1) proteins on cell viability under stress (Figure 4G), 1e^4^ HEK 293T cells stably expressing shRNA targeting each protein were plated in 96-well plates with 100 μL fresh medium per well and cultured for 24 hours. Then the cells were treated with 500 μM sodium arsenite for 1 hour at 37°C.

For all the experiments above, cell viability was determined by adding 20 μL of CellTiter 96 AQueous One Solution Reagent (Promega) into each well. After incubation for 4 hours, the absorbance at 490 nm was recorded using a 96-well plate reader. Each biological experiment has five technical replicates and three biological replicates were performed.

#### Statistical analysis

For comparison between two groups, P values were determined using two-tailed Student’s t tests, *P < 0.05; **P < 0.01; ***P < 0.001; N.S. not significant. For all box plots (Figure 3E, 5H, 5I, Supplementary Figure 3F), P values were calculated with Wilcoxon rank sum by R (*P < 0.05; **P < 0.01; ***P < 0.001). Error bars represent means ± SD.

#### Data availability

Additional data beyond that provided in the Figures and Supplementary Information are available from the corresponding author upon request.

The original mass spectra, spectral library, and the protein sequence database used for searches have been deposited in the public proteomics repository MassIVE (http://massive.ucsd.edu) and are accessible at ftp://MSV000090740@massive.ucsd.edu. when providing the dataset password: trafficking. If requested, also provide the username: MSV000090740. These datasets will be made public upon acceptance of the manuscript.

## References

1. Cook, K.C., and Cristea, I.M. (2019). Location is everything: protein translocations as a viral infection strategy. Curr Opin Chem Biol 48, 34–43. 10.1016/j.cbpa.2018.09.021.

2. Jones, S., and Rappoport, J.Z. (2014). Interdependent epidermal growth factor receptor signalling and trafficking. Int J Biochem Cell Biol 51, 23–28. 10.1016/j.biocel.2014.03.014.

3. Qin, W., Cho, K.F., Cavanagh, P.E., and Ting, A.Y. (2021). Deciphering molecular interactions by proximity labeling. Nat Methods 18, 133–143. 10.1038/s41592-020-01010-5.

4. Lam, S.S., Martell, J.D., Kamer, K.J., Deerinck, T.J., Ellisman, M.H., Mootha, V.K., and Ting, A.Y. (2015). Directed evolution of APEX2 for electron microscopy and proximity labeling. Nat Methods 12, 51–54. 10.1038/nmeth.3179.

5. Roux, K.J., Kim, D.I., Raida, M., and Burke, B. (2012). A promiscuous biotin ligase fusion protein identifies proximal and interacting proteins in mammalian cells. J Cell Biol 196, 801–810. 10.1083/jcb.201112098.

6. Choi-Rhee, E., Schulman, H., and Cronan, J.E. (2004). Promiscuous protein biotinylation by Escherichia coli biotin protein ligase. Protein Sci 13, 3043–3050. 10.1110/ps.04911804.

7. Branon, T.C., Bosch, J.A., Sanchez, A.D., Udeshi, N.D., Svinkina, T., Carr, S.A., Feldman, J.L., Perrimon, N., and Ting, A.Y. (2018). Efficient proximity labeling in living cells and organisms with TurboID. Nat Biotechnol 36, 880–887. 10.1038/nbt.4201.

8. Go, C.D., Knight, J.D.R., Rajasekharan, A., Rathod, B., Hesketh, G.G., Abe, K.T., Youn, J.-Y., Samavarchi-Tehrani, P., Zhang, H., Zhu, L.Y., et al. (2021). A proximity-dependent biotinylation map of a human cell. Nature 595, 120–124. 10.1038/s41586-021-03592-2.

9. Youn, J.-Y., Dunham, W.H., Hong, S.J., Knight, J.D.R., Bashkurov, M., Chen, G.I., Bagci, H., Rathod, B., MacLeod, G., Eng, S.W.M., et al. (2018). High-Density Proximity Mapping Reveals the Subcellular Organization of mRNA-Associated Granules and Bodies. Mol Cell 69, 517–532.e11. 10.1016/j.molcel.2017.12.020.

10. Markmiller, S., Soltanieh, S., Server, K.L., Mak, R., Jin, W., Fang, M.Y., Luo, E.-C., Krach, F., Yang, D., Sen, A., et al. (2018). Context-Dependent and Disease-Specific Diversity in Protein Interactions within Stress Granules. Cell 172, 590–604.e13. 10.1016/j.cell.2017.12.032.

11. Lobingier, B.T., Hüttenhain, R., Eichel, K., Miller, K.B., Ting, A.Y., von Zastrow, M., and Krogan, N.J. (2017). An Approach to Spatiotemporally Resolve Protein Interaction Networks in Living Cells. Cell. 10.1016/j.cell.2017.03.022.

12. Paek, J., Kalocsay, M., Staus, D.P., Wingler, L., Pascolutti, R., Paulo, J.A., Gygi, S.P., and Kruse, A.C. (2017). Multidimensional Tracking of GPCR Signaling via Peroxidase-Catalyzed Proximity Labeling. Cell. 10.1016/j.cell.2017.03.028.

13. Miki, T., Awa, M., Nishikawa, Y., Kiyonaka, S., Wakabayashi, M., Ishihama, Y., and Hamachi, I. (2016). A conditional proteomics approach to identify proteins involved in zinc homeostasis. Nat Methods 13, 931–937. 10.1038/nmeth.3998.

14. Williams, C.C., Jan, C.H., and Weissman, J.S. (2014). Targeting and plasticity of mitochondrial proteins revealed by proximity-specific ribosome profiling. Science 346, 748–751. 10.1126/science.1257522.

15. Fazal, F.M., Han, S., Parker, K.R., Kaewsapsak, P., Xu, J., Boettiger, A.N., Chang, H.Y., and Ting, A.Y. (2019). Atlas of Subcellular RNA Localization Revealed by APEX-Seq. Cell 178, 473–490.e26. 10.1016/j.cell.2019.05.027.

16. Rhee, H.-W., Zou, P., Udeshi, N.D., Martell, J.D., Mootha, V.K., Carr, S.A., and Ting, A.Y. (2013). Proteomic mapping of mitochondria in living cells via spatially restricted enzymatic tagging. Science 339, 1328–1331. 10.1126/science.1230593.

17. McShane, E., Sin, C., Zauber, H., Wells, J.N., Donnelly, N., Wang, X., Hou, J., Chen, W., Storchova, Z., Marsh, J.A., et al. (2016). Kinetic Analysis of Protein Stability Reveals Age-Dependent Degradation. Cell 167, 803–815.e21. 10.1016/j.cell.2016.09.015.

18. Liu, J., Xu, Y., Stoleru, D., and Salic, A. (2012). Imaging protein synthesis in cells and tissues with an alkyne analog of puromycin. Proc Natl Acad Sci U S A 109, 413–418. 10.1073/pnas.1111561108.

19. Zhu, Y., Willems, L.I., Salas, D., Cecioni, S., Wu, W.B., Foster, L.J., and Vocadlo D.J. (2020). Tandem Bioorthogonal Labeling Uncovers Endogenous Cotranslationally O-GlcNAc Modified Nascent Proteins. J Am Chem Soc 142, 15729–15739. 10.1021/jacs.0c04121.

20. Freibaum, B.D., Lu, Y., Lopez-Gonzalez, R., Kim, N.C., Almeida, S., Lee, K.-H., Badders, N., Valentine, M., Miller, B.L., Wong, P.C., et al. (2015). GGGGCC repeat expansion in C9orf72 compromises nucleocytoplasmic transport. Nature 525, 129–133. 10.1038/nature14974.

21. Pakos-Zebrucka, K., Koryga, I., Mnich, K., Ljujic, M., Samali, A., and Gorman, A.M. (2016). The integrated stress response. EMBO Rep 17, 1374–1395. 10.15252/embr.201642195.

22. Neumann, M., Sampathu, D.M., Kwong, L.K., Truax, A.C., Micsenyi, M.C., Chou, T.T., Bruce, J., Schuck, T., Grossman, M., Clark, C.M., et al. (2006). Ubiquitinated TDP-43 in frontotemporal lobar degeneration and amyotrophic lateral sclerosis. Science 314, 130–133. 10.1126/science.1134108.

23. Meneses, A., Koga, S., O’Leary, J., Dickson, D.W., Bu, G., and Zhao, N. (2021). TDP-43 Pathology in Alzheimer’s Disease. Mol Neurodegener 16, 84. 10.1186/s13024-021-00503-x.

24. Ma, X.R., Prudencio, M., Koike, Y., Vatsavayai, S.C., Kim, G., Harbinski, F., Briner, A., Rodriguez, C.M., Guo, C., Akiyama, T., et al. (2022). TDP-43 represses cryptic exon inclusion in the FTD-ALS gene UNC13A. Nature 603, 124–130. 10.1038/s41586-022-04424-7.

25. Protter, D.S.W., and Parker, R. (2016). Principles and Properties of Stress Granules. Trends Cell Biol 26, 668–679. 10.1016/j.tcb.2016.05.004.

26. Wolozin, B., and Ivanov, P. (2019). Stress granules and neurodegeneration. Nat Rev Neurosci 20, 649–666. 10.1038/s41583-019-0222-5.

27. Boulon, S., Westman, B.J., Hutten, S., Boisvert, F.-M., and Lamond, A.I. (2010). The nucleolus under stress. Mol Cell 40, 216–227. 10.1016/j.molcel.2010.09.024.

28. Zhu, C.-H., Kim, J., Shay, J.W., and Wright, W.E. (2008). SGNP: an essential Stress Granule/Nucleolar Protein potentially involved in 5.8s rRNA processing/transport. PLoS One 3, e3716. 10.1371/journal.pone.0003716.

29. Alastalo, T.-P., Hellesuo, M., Sandqvist, A., Hietakangas, V., Kallio, M., and Sistonen, L. (2003). Formation of nuclear stress granules involves HSF2 and coincides with the nucleolar localization of Hsp70. J Cell Sci 116, 3557–3570. 10.1242/jcs.00671.

30. Mahboubi, H., and Stochaj, U. (2014). Nucleoli and stress granules: connecting distant relatives. Traffic 15, 1179–1193. 10.1111/tra.12191.

31. Suk, T.R., and Rousseaux, M.W.C. (2020). The role of TDP-43 mislocalization in amyotrophic lateral sclerosis. Mol Neurodegener 15, 45. 10.1186/s13024-020-00397-1.

32. Vernon, R.M., Chong, P.A., Tsang, B., Kim, T.H., Bah, A., Farber, P., Lin, H., and Forman-Kay, J.D. (2018). Pi-Pi contacts are an overlooked protein feature relevant to phase separation. Elife 7. 10.7554/eLife.31486.

33. Erdős, G., Pajkos, M., and Dosztányi, Z. (2021). IUPred3: prediction of protein disorder enhanced with unambiguous experimental annotation and visualization of evolutionary conservation. Nucleic Acids Res 49, W297–W303. 10.1093/nar/gkab408.

34. Maxwell, B.A., Gwon, Y., Mishra, A., Peng, J., Nakamura, H., Zhang, K., Kim, H.J., and Taylor, J.P. (2021). Ubiquitination is essential for recovery of cellular activities after heat shock. Science 372, eabc3593. 10.1126/science.abc3593.

35. Gwon, Y., Maxwell, B.A., Kolaitis, R.-M., Zhang, P., Kim, H.J., and Taylor, J.P. (2021). Ubiquitination of G3BP1 mediates stress granule disassembly in a context-specific manner. Science 372, eabf6548. 10.1126/science.abf6548.

36. Wheeler, J.R., Matheny, T., Jain, S., Abrisch, R., and Parker, R. (2016). Distinct stages in stress granule assembly and disassembly. Elife 5. 10.7554/eLife.18413.

37. Latonen, L. (2019). Phase-to-Phase With Nucleoli - Stress Responses, Protein Aggregation and Novel Roles of RNA. Front Cell Neurosci 13, 151. 10.3389/fncel.2019.00151.

38. Yang, P., Mathieu, C., Kolaitis, R.-M., Zhang, P., Messing, J., Yurtsever, U., Yang, Z., Wu, J., Li, Y., Pan, Q., et al. (2020). G3BP1 Is a Tunable Switch that Triggers Phase Separation to Assemble Stress Granules. Cell 181, 325–345.e28. 10.1016/j.cell.2020.03.046.

39. Sanders, D.W., Kedersha, N., Lee, D.S.W., Strom, A.R., Drake, V., Riback, J.A., Bracha, D., Eeftens, J.M., Iwanicki, A., Wang, A., et al. (2020). Competing Protein-RNA Interaction Networks Control Multiphase Intracellular Organization. Cell 181, 306–324.e28. 10.1016/j.cell.2020.03.050.

40. Huang, L., Agrawal, T., Zhu, G., Yu, S., Tao, L., Lin, J., Marmorstein, R., Shorter, J., and Yang, X. (2021). DAXX represents a new type of protein-folding enabler. Nature 597, 132–137. 10.1038/s41586-021-03824-5.

41. Shaulian, E., and Karin, M. (2002). AP-1 as a regulator of cell life and death. Nat Cell Biol 4, E131–6. 10.1038/ncb0502-e131.

42. Dérijard, B., Hibi, M., Wu, I.H., Barrett, T., Su, B., Deng, T., Karin, M., and Davis, R.J. (1994). JNK1: a protein kinase stimulated by UV light and Ha-Ras that binds and phosphorylates the c-Jun activation domain. Cell 76, 1025–1037. 10.1016/0092-8674(94)90380-8.

43. Behrens, A., Sibilia, M., and Wagner, E.F. (1999). Amino-terminal phosphorylation of c-Jun regulates stress-induced apoptosis and cellular proliferation. Nat Genet 21, 326–329. 10.1038/6854.

44. Angel, P., Hattori, K., Smeal, T., and Karin, M. (1988). The jun proto-oncogene is positively autoregulated by its product, Jun/AP-1. Cell 55, 875–885. 10.1016/0092-8674(88)90143-2.

45. Liu, J., Perumal, N.B., Oldfield, C.J., Su, E.W., Uversky, V.N., and Dunker, A.K. (2006). Intrinsic disorder in transcription factors. Biochemistry 45, 6873–6888. 10.1021/bi0602718.

46. Lee, K.-H., Zhang, P., Kim, H.J., Mitrea, D.M., Sarkar, M., Freibaum, B.D., Cika, J., Coughlin, M., Messing, J., Molliex, A., et al. (2016). C9orf72 Dipeptide Repeats Impair the Assembly, Dynamics, and Function of Membrane-Less Organelles. Cell 167, 774–788.e17. 10.1016/j.cell.2016.10.002.

47. Boeynaems, S., Bogaert, E., Kovacs, D., Konijnenberg, A., Timmerman, E., Volkov, A., Guharoy, M., de Decker, M., Jaspers, T., Ryan, V.H., et al. (2017). Phase Separation of C9orf72 Dipeptide Repeats Perturbs Stress Granule Dynamics. Mol Cell 65, 1044–1055.e5. 10.1016/j.molcel.2017.02.013.

48. Mann, J.R., Gleixner, A.M., Mauna, J.C., Gomes, E., DeChellis-Marks, M.R., Needham, P.G., Copley, K.E., Hurtle, B., Portz, B., Pyles, N.J., et al. (2019). RNA Binding Antagonizes Neurotoxic Phase Transitions of TDP-43. Neuron 102, 321–338.e8. 10.1016/j.neuron.2019.01.048.

49. McGurk, L., Gomes, E., Guo, L., Mojsilovic-Petrovic, J., Tran, V., Kalb, R.G., Shorter, J., and Bonini, N.M. (2018). Poly(ADP-Ribose) Prevents Pathological Phase Separation of TDP-43 by Promoting Liquid Demixing and Stress Granule Localization. Mol Cell 71, 703–717.e9. 10.1016/j.molcel.2018.07.002.

50. Dieterich, D.C., Link, A.J., Graumann, J., Tirrell, D.A., and Schuman, E.M. (2006). Selective identification of newly synthesized proteins in mammalian cells using bioorthogonal noncanonical amino acid tagging (BONCAT). Proc Natl Acad Sci U S A 103, 9482–9487. 10.1073/pnas.0601637103.

51. Balta, E., Wabnitz, G.H., and Samstag, Y. (2021). Hijacked Immune Cells in the Tumor Microenvironment: Molecular Mechanisms of Immunosuppression and Cues to Improve T Cell-Based Immunotherapy of Solid Tumors. Int J Mol Sci 22. 10.3390/ijms22115736.

52. Zhou, J., Tang, Z., Gao, S., Li, C., Feng, Y., and Zhou, X. (2020). Tumor-Associated Macrophages: Recent Insights and Therapies. Front Oncol 10, 188. 10.3389/fonc.2020.00188.

53. Mantovani, A., Allavena, P., Marchesi, F., and Garlanda, C. (2022). Macrophages as tools and targets in cancer therapy. Nat Rev Drug Discov. 10.1038/s41573-022-00520-5.

54. Wei, X., Ye, J., Pei, Y., Wang, C., Yang, H., Tian, J., Si, G., Ma, Y., Wang, K., and Liu, G. (2022). Extracellular vesicles from colorectal cancer cells promote metastasis via the NOD1 signalling pathway. J Extracell Vesicles 11, e12264. 10.1002/jev2.12264.

55. Putz, U., Howitt, J., Doan, A., Goh, C.-P., Low, L.-H., Silke, J., and Tan, S.-S. (2012). The tumor suppressor PTEN is exported in exosomes and has phosphatase activity in recipient cells. Sci Signal 5, ra70. 10.1126/scisignal.2003084.

56. Hanna, S.J., McCoy-Simandle, K., Leung, E., Genna, A., Condeelis, J., and Cox, D. (2019). Tunneling nanotubes, a novel mode of tumor cell-macrophage communication in tumor cell invasion. J Cell Sci 132. 10.1242/jcs.223321.

57. Saha, T., Dash, C., Jayabalan, R., Khiste, S., Kulkarni, A., Kurmi, K., Mondal, J., Majumder, P.K., Bardia, A., Jang, H.L., et al. (2022). Intercellular nanotubes mediate mitochondrial trafficking between cancer and immune cells. Nat Nanotechnol 17, 98–106. 10.1038/s41565-021-01000-4.

58. Wu, C., Xue, Y., Wang, P., Lin, L., Liu, Q., Li, N., Xu, J., and Cao, X. (2014). IFN-γ primes macrophage activation by increasing phosphatase and tensin homolog via downregulation of miR-3473b. J Immunol 193, 3036–3044. 10.4049/jimmunol.1302379.

59. Luzina, I.G., Keegan, A.D., Heller, N.M., Rook, G.A.W., Shea-Donohue, T., and Atamas, S.P. (2012). Regulation of inflammation by interleukin-4: a review of “alternatives”. J Leukoc Biol 92, 753–764. 10.1189/jlb.0412214.

60. Droujinine, I.A., Meyer, A.S., Wang, D., Udeshi, N.D., Hu, Y., Rocco, D., McMahon, J.A., Yang, R., Guo, J., Mu, L., et al. (2021). Proteomics of protein trafficking by in vivo tissue-specific labeling. Nat Commun 12, 2382. 10.1038/s41467-021-22599-x.

61. Wei, W., Riley, N.M., Yang, A.C., Kim, J.T., Terrell, S.M., Li, V.L., Garcia-Contreras, M., Bertozzi, C.R., and Long, J.Z. (2021). Cell type-selective secretome profiling in vivo. Nat Chem Biol 17, 326–334. 10.1038/s41589-020-00698-y.

62. Kim, K.-E., Park, I., Kim, J., Kang, M.-G., Choi, W.G., Shin, H., Kim, J.-S., Rhee, H.-W., and Suh, J.M. (2021). Dynamic tracking and identification of tissue-specific secretory proteins in the circulation of live mice. Nat Commun 12, 5204. 10.1038/s41467-021-25546-y.

63. Liddelow, S.A., Guttenplan, K.A., Clarke, L.E., Bennett, F.C., Bohlen, C.J., Schirmer, L., Bennett, M.L., Münch, A.E., Chung, W.-S., Peterson, T.C., et al. (2017). Neurotoxic reactive astrocytes are induced by activated microglia. Nature 541, 481–487. 10.1038/nature21029.

64. van der Leun, A.M., Thommen, D.S., and Schumacher, T.N. (2020). CD8+ T cell states in human cancer: insights from single-cell analysis. Nat Rev Cancer 20, 218– 232. 10.1038/s41568-019-0235-4.

65. Schäfer, J.A., Bozkurt, S., Michaelis, J.B., Klann, K., and Münch, C. (2022). Global mitochondrial protein import proteomics reveal distinct regulation by translation and translocation machinery. Mol Cell 82, 435–446.e7. 10.1016/j.molcel.2021.11.004.

66. Qin, W., Myers, S.A., Carey, D.K., Carr, S.A., and Ting, A.Y. (2021). Spatiotemporally-resolved mapping of RNA binding proteins via functional proximity labeling reveals a mitochondrial mRNA anchor promoting stress recovery. Nat Commun 12, 4980. 10.1038/s41467-021-25259-2.

67. Fischer, J.W., Busa, V.F., Shao, Y., and Leung, A.K.L. (2020). Structure-Mediated RNA Decay by UPF1 and G3BP1. Mol Cell 78, 70–84.e6. 10.1016/j.molcel.2020.01.021.

68. Hung, V., Lam, S.S., Udeshi, N.D., Svinkina, T., Guzman, G., Mootha, V.K., Carr, S.A., and Ting, A.Y. (2017). Proteomic mapping of cytosol-facing outer mitochondrial and ER membranes in living human cells by proximity biotinylation. Elife 6. 10.7554/eLife.24463.

69. Marmor-Kollet, H., Siany, A., Kedersha, N., Knafo, N., Rivkin, N., Danino, Y.M., Moens, T.G., Olender, T., Sheban, D., Cohen, N., et al. (2020). Spatiotemporal Proteomic Analysis of Stress Granule Disassembly Using APEX Reveals Regulation by SUMOylation and Links to ALS Pathogenesis. Mol Cell 80, 876–891.e6. 10.1016/j.molcel.2020.10.032.

70. Senichkin, V. v, Prokhorova, E.A., Zhivotovsky, B., and Kopeina, G.S. (2021). Simple and Efficient Protocol for Subcellular Fractionation of Normal and Apoptotic Cells. Cells 10. 10.3390/cells10040852.

