## Supplementary Information for "Dynamic mapping of proteome trafficking within and between living cells by TransitID"

^1^Departments of Biology, Genetics, and Chemistry, Stanford University, Stanford, CA, USA; ^2^Chan Zuckerberg Biohub, San Francisco, CA, USA; ^3^The Broad Institute of MIT and Harvard, Cambridge, MA, USA; ^4^Department of Cell and Molecular Biology, St. Jude Children’s Research Hospital, Memphis, TN, USA; ^5^Department of Molecular and Human Genetics, Therapeutic Innovation Center, Center for Alzheimer’s and Neurodegenerative Diseases, and Dan L. Duncan Comprehensive Cancer Center, Baylor College of Medicine, Houston, TX, USA; ^6^Jan and Dan Duncan Neurological Research Institute, Texas Children’s Hospital, Houston, TX, USA; ^7^Current address: School of Pharmaceutical Sciences, Tsinghua-Peking Center for Life Sciences, Tsinghua University, Beijing, China.

This file includes Supplementary texts 1-3, Supplementary figures 1-10, Supplementary tables 1-4 and Supplementary movies 1-2.

**Supplementary texts**

**Supplementary text 1. Analysis of combined list of 670 proteins that traffick from the cytosol or OMM to the mitochondrial matrix**

For our combined list of 670 proteins that traffick from the cytosol or OMM to the mitochondrial matrix in HEK 293T cells, we observed high enrichment of nuclear-encoded mitochondrial matrix proteins (>80%, **Figures S3A-S3B**). Analysis of the sub-mitochondrial specificity of this dataset showed primarily mitochondrial matrix and inner mitochondrial membrane (IMM) proteins, with a lack of IMS and OMM-resident proteins (**Figure S3C**). Consistent with this, only matrix-accessible subunits of the TOM/TIM/PAM protein-import complex were enriched in the combined dataset (**Figure S3D**). None of the 13 mtDNA-encoded proteins were enriched, as expected due to the use of cytosolic or OMM TurboID in the first labeling step. This is also consistent with our Western blot validation in **Figure 2D**.

To evaluate depth of coverage, we checked the representation of well-established mitochondrial matrix protein groups in our dataset of 670 proteins. More than 80% of “true positive” proteins were detected in each group (**Figure S3E**). We conclude that our dataset has comparable sensitivity to our previous single-step mito-APEX1-mapped mitochondrial matrix proteome^1^.

**Supplementary text 2. Analysis of 148 proteins that preferentially translocate from the OMM to the mitochondrial matrix**

To further analyze our list of 148 proteins that preferentially translocate from the OMM (rather than cytosol) to the mitochondrial matrix, we crossed our dataset with the mePRODmt database^2^, which quantifies the kinetics of mitochondrial protein uptake. We found that OMM-to-mito matrix proteins are generally imported faster (**Figure 3E**), which may result from their local synthesis and proximity to the OMM. Gene Ontology (GO) analysis showed that OMM-enriched translocated proteins are enriched in the biological processes of mitochondrial translation and oxidative phosphorylation (OXPHOS) (**Figure 3F**). Interestingly, the mitochondrial ribosome and OXPHOS are the two mitochondrial complexes that require coordinated assembly from nuclear and mitochondrial genomes^3^. Perhaps local translation at the OMM provides an efficient mechanism to coordinate the syntheses of nuclear-encoded and mitochondrial genome-encoded components of these two essential macromolecular complexes.

**Supplementary text 3. Probing JUN transcriptional activity in the context of SGs.**

We performed several assays to evaluate the role of SGs in JUN’s post-stress DNA binding and transcriptional activity. First, we used co-IP to measure JUN’s interaction with its endogenous binding partner FOS, and found, as expected, that stress reduced the interaction, while interaction was restored post stress recovery. We repeated the same experiment in DKO cells that are unable to form SGs, and found that while basal and post-stress levels of JUN-FOS interaction were unchanged, the extent of FOS interaction with JUN during stress recovery was markedly decreased (**Figure 6E**). This is consistent with our earlier observation that SGs reduce JUN aggregation and degradation upon stress, leaving more soluble JUN available to form complexes with FOS.

Second, we probed JUN’s DNA binding activity by performing an ELISA binding assay to JUN’s double stranded DNA promoter sequence. We found that JUN’s DNA binding decreased under stress, and increased upon stress recovery, but not in DKO cells (**Figure 6F**). A similar effect on JUN’s DNA binding activity was observed when cells were treated with GSK260641435^4^, a PERK inhibitor known to disrupt SG assembly (**Figure S8D**).

Third, we measured JUN’s ability to upregulate its own synthesis post-stress. **Figure 6G** shows that JUN protein levels increased dramatically at 3-hour post-stress removal, while this phenomenon was not observed in DKO cells. The timing of JUN induction may correlate with the restart of cellular protein synthesis ~ 2 hours after stress removal. To test this, we used AHA pulse labeling^5^ to tag newly synthesized JUN protein, and observed 2.5-fold more new JUN 3 hours after stress removal compared to basal conditions. By contrast, synthesis of new JUN was inhibited in DKO cells (**Figure S8E**). Collectively, these assays indicate that JUN relocalization to SGs is important for full restoration of its FOS interaction, DNA binding, and transcriptional activity during stress recovery.

**Supplementary figures**

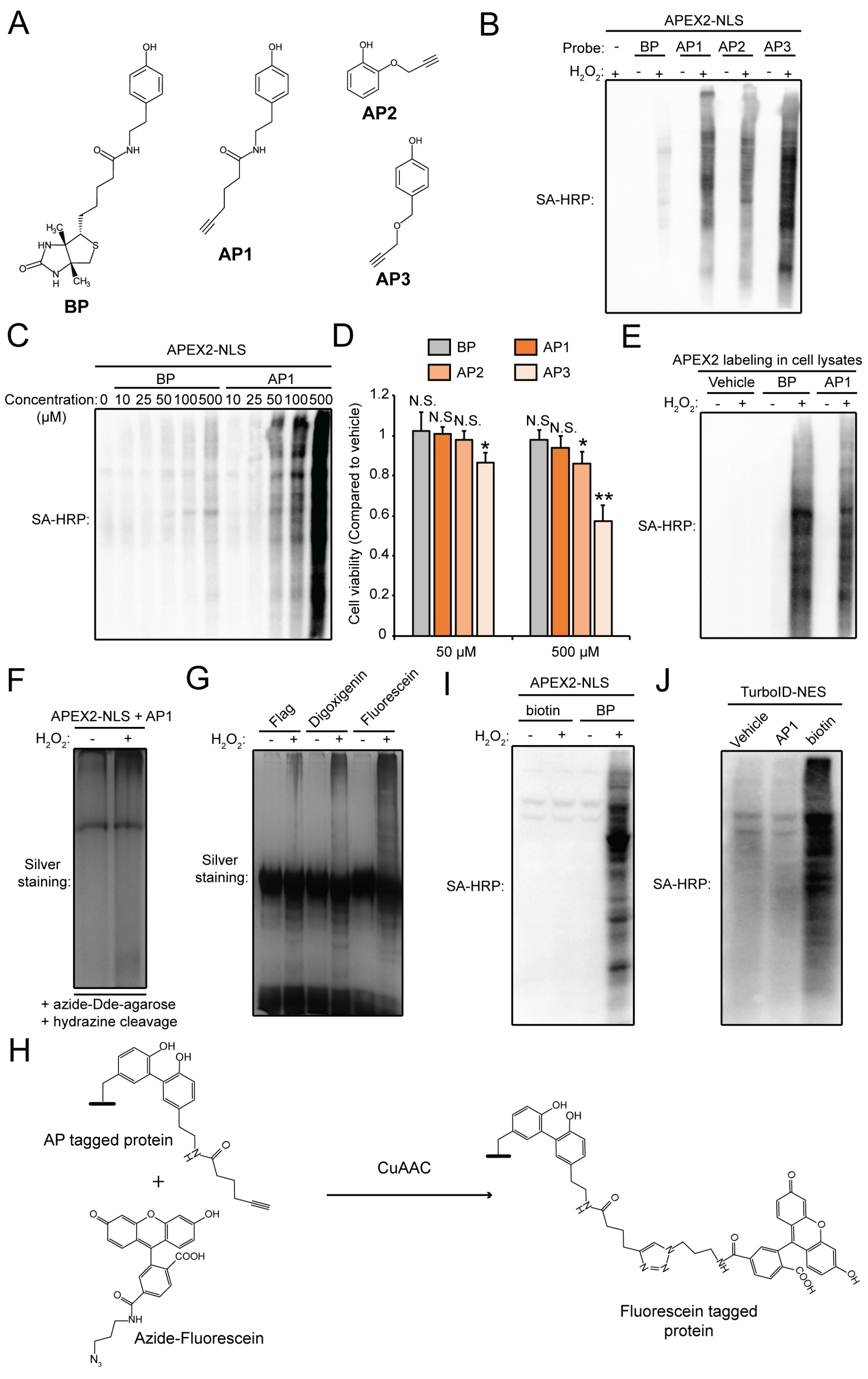

**Figure S1. Development of TransitID for mapping proteome translocation, related to Figure 1. A.** Chemical structures of biotin-phenol (BP) and alkyne-phenols (AP1-3). **B**. Streptavidin blot of lysates from HEK cells expressing nuclear APEX2 and labeled with the indicated probes for 1 minute. AP-labeled samples were derivatized with azide-biotin before analysis. **C**. Concentration-dependent labeling by AP1 and BP in HEK. **D**. Cell toxicity of AP and BP probes in HEK 293T cells. **E**. Streptavidin blot of APEX labeling in cell lysates with indicated probes. **F**. Enrichment of AP-labeled proteins by Click reaction with azide-Dde [N-1-(4,4-dimethyl-2,6-dioxocyclohex-1-ylidene)-3-ethyl]-agarose beads followed by hydrazine cleavage. **G**. Enrichment of AP-labeled proteins by Click reaction with azide conjugated to the indicated affinity tags (Flag, digoxigenin, and fluorescein (FAM)), followed by IP by their corresponding antibodies. **H**. Click reaction between AP-labeled proteins and azide-fluorescein. **I**. APEX2 does not use biotin as a substrate. **J**. TurboID does not use AP1 as a substrate.

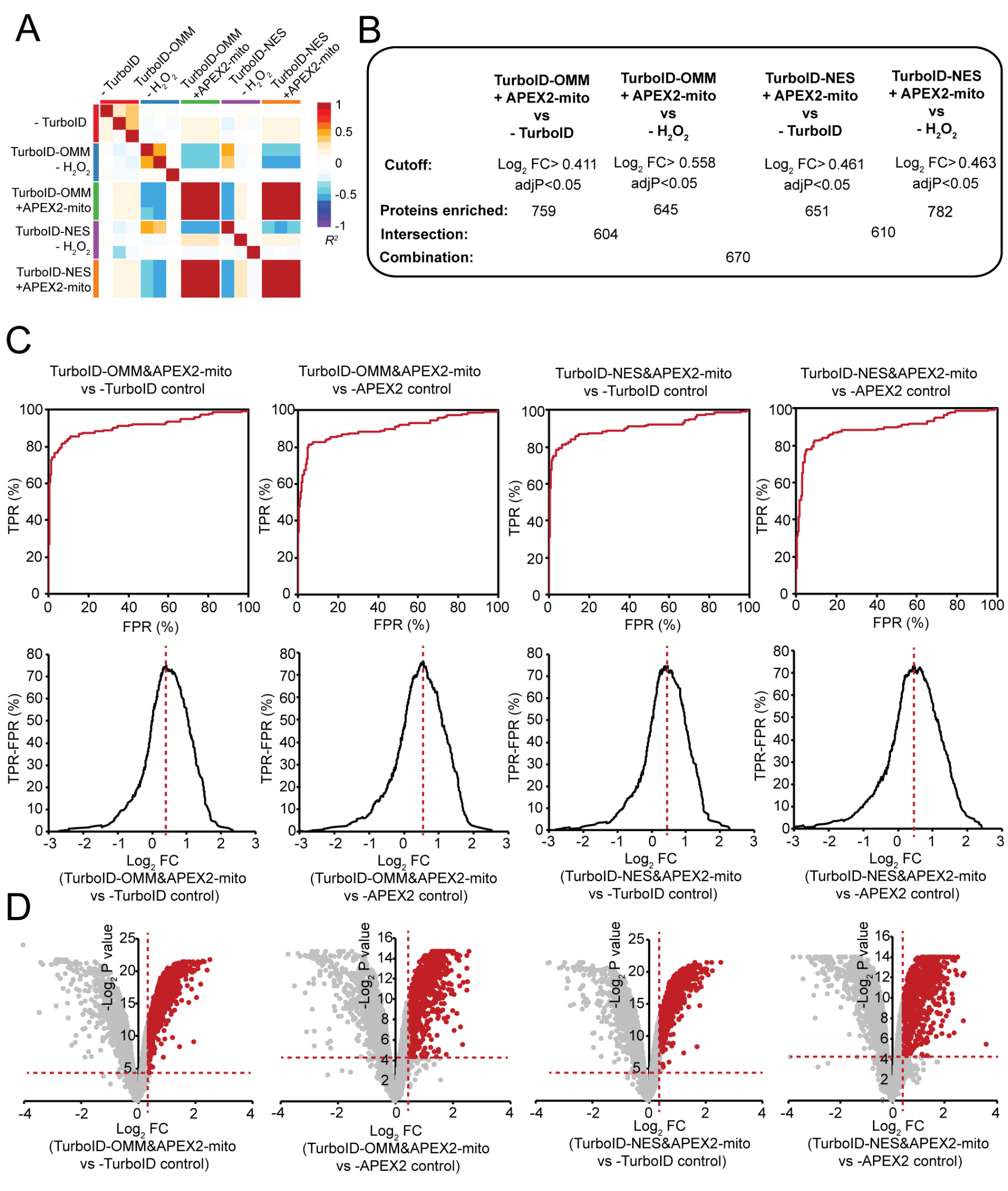

**Figure S2. Analysis of proteomic data for cytosol/OMM-to-mitochondrial matrix trafficking experiment shown in Figure 3. A.** Correlation between biological replicates in the TMT experiment. **B.** Filtering protocol for the mass spec data. Number of proteins remaining after each filtering step is given. **C.** Receiver operating characteristic (ROC) curves for the indicated TMT ratios (across top). Proteins were ranked in descending order based on TMT ratio. True positives are known nuclear-encoded mitochondrial proteins. False positives are annotated plasma membrane proteins. **D.** Filtering of each comparison by ROC cutoffs and p-values.

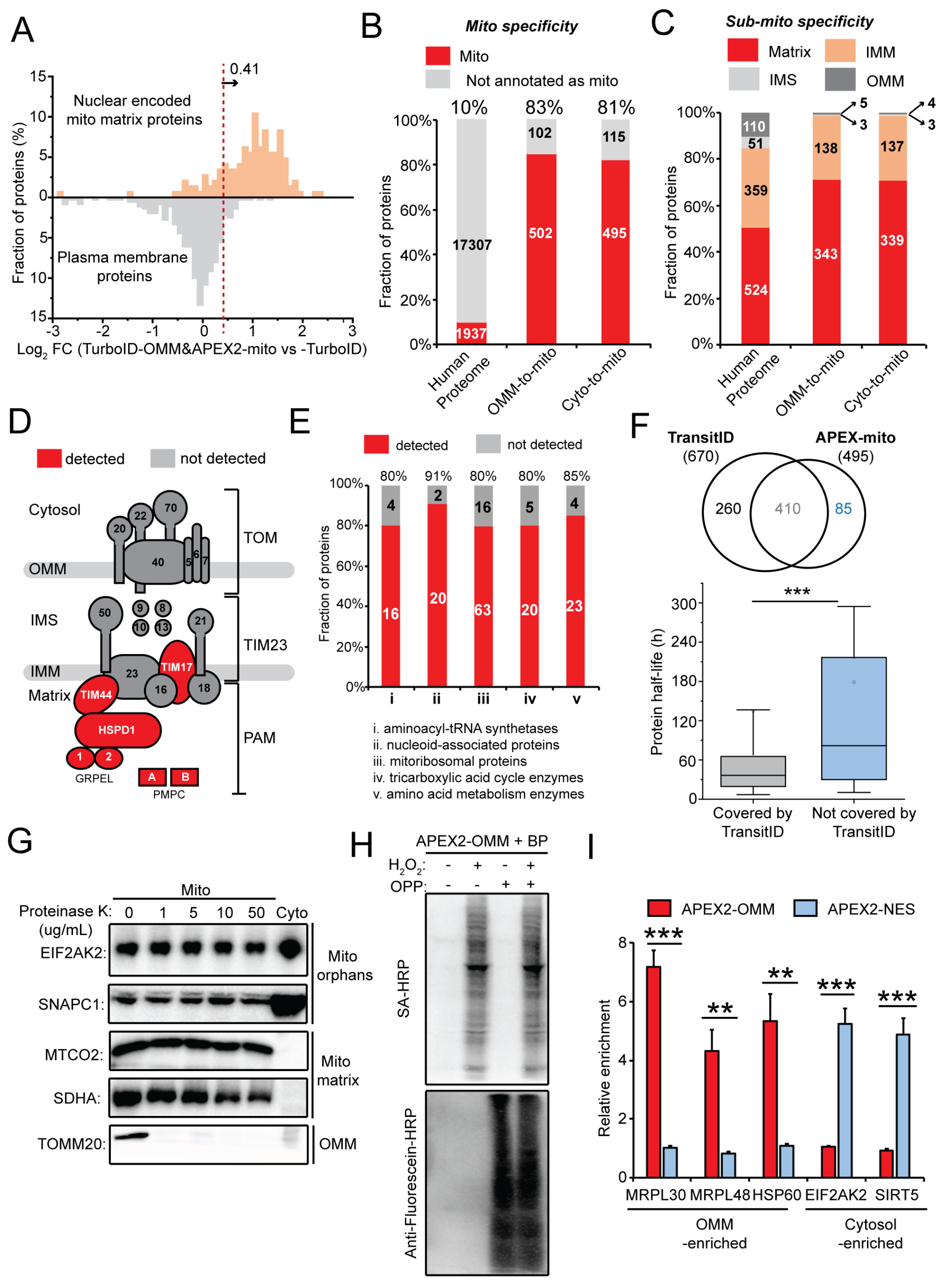

**Figure S3. Further analysis of cytosol/OMM-to-mitochondrial matrix proteomes. Related to Figure 3. A.** Sample histogram showing how the ROC-determined cutoff was applied. **B-C**. Mitochondrial (**B**) and sub-mitochondrial (**C**) specificity of cyto-to-mito and OMM-to-mito datasets. **D.** The components of TOM/TIM/PAM complex identified in either dataset. **E.** Coverage/sensitivity analysis for TransitID datasets. **F.** Overlap between our TransitID combined dataset and previous single PL (APEX) mapping of the mitochondrial matrix^1^. Distribution of protein half-lives according to *McShane et al., 2016* ^6^. **G.** Validation of mitochondrial orphans by mitochondrial fractionation with proteinase K digestion. Positive and negative control protein markers (MTCO2, SDHA and TOMM20) are shown. **H.** Puromycin tagging of newly synthesized polypeptides followed by APEX2-OMM-catalyzed biotinylation, as in **Figure 3G**. Streptavidin and anti-fluorescein blotting of whole cell lysates shown. **I.** Quantification of western blot data shown in **Figure 3I** from three biological replicates.

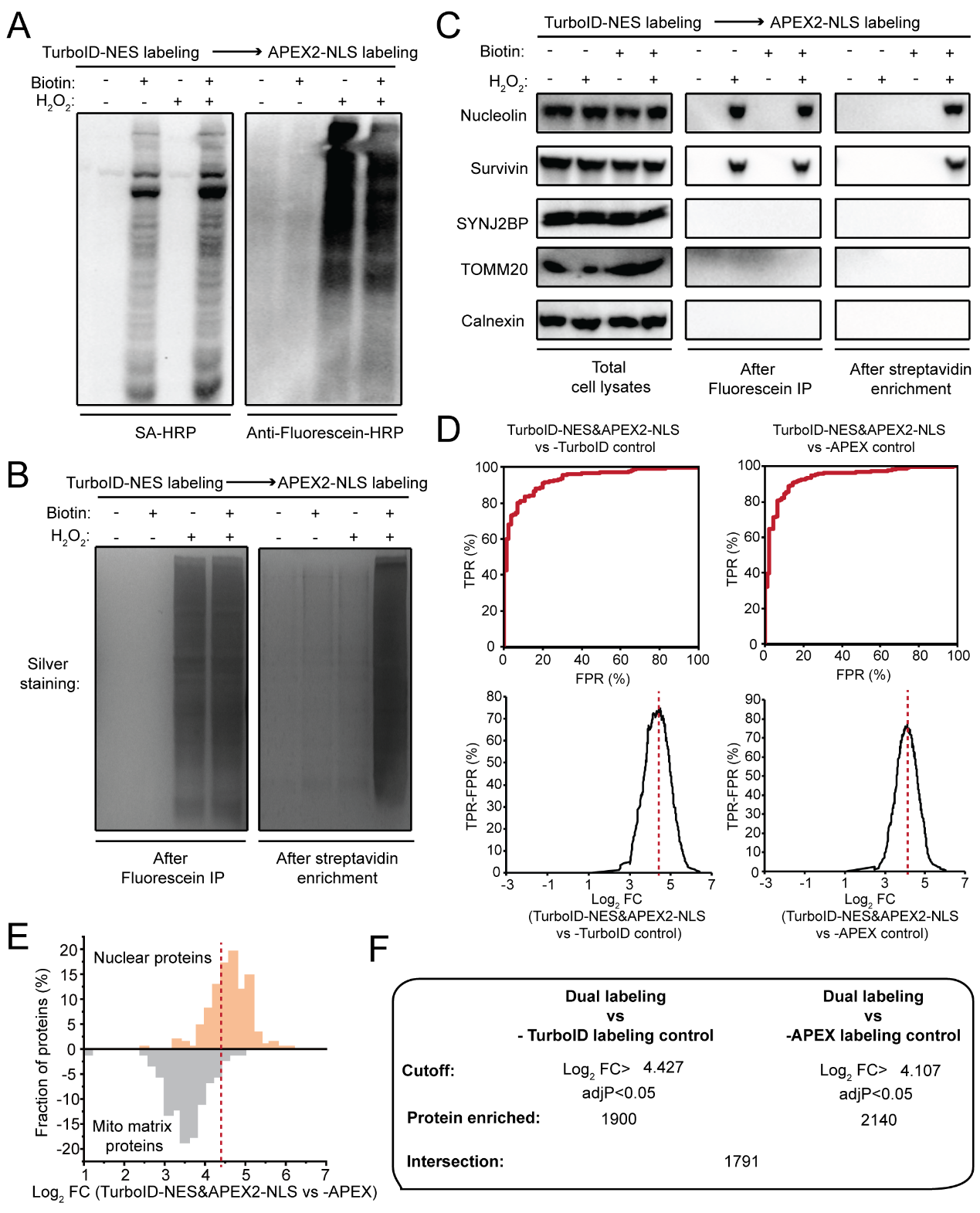

**Figure S4. Additional characterization of TransitID samples for cytosol-to-nucleus trafficking under stress. Related to Figure 4. A**. Streptavidin and anti-fluorescein blotting of cells labeled as in **Figure 4B**, along with negative controls omitting biotin or H_2_O_2_. **B**. Silver staining of enriched proteins after anti-fluorescein IP (left) and after the second streptavidin bead enrichment (right). **C**. Western blot detection of protein markers in samples from (A) after cell lysis (left), after anti-fluorescein IP (middle), and after the second streptavidin enrichment (right). Nucleolin and survivin are true-positive cytosol to nucleus translocated proteins. TOMM20 and SYNJ2BP (OMM proteins) and Calnexin (ERM protein) are true negatives. **D.** Receiver operating characteristic (ROC) curves for the indicated TMT ratios. Proteins were ranked in descending order by TMT ratio. True positives are known nuclear proteins. False positives are mitochondrial matrix proteins. **E.** Sample histogram showing how the ROC-derived cutoff was applied. **F.** Number of proteins remaining after each step of filtering the mass spectrometry data.

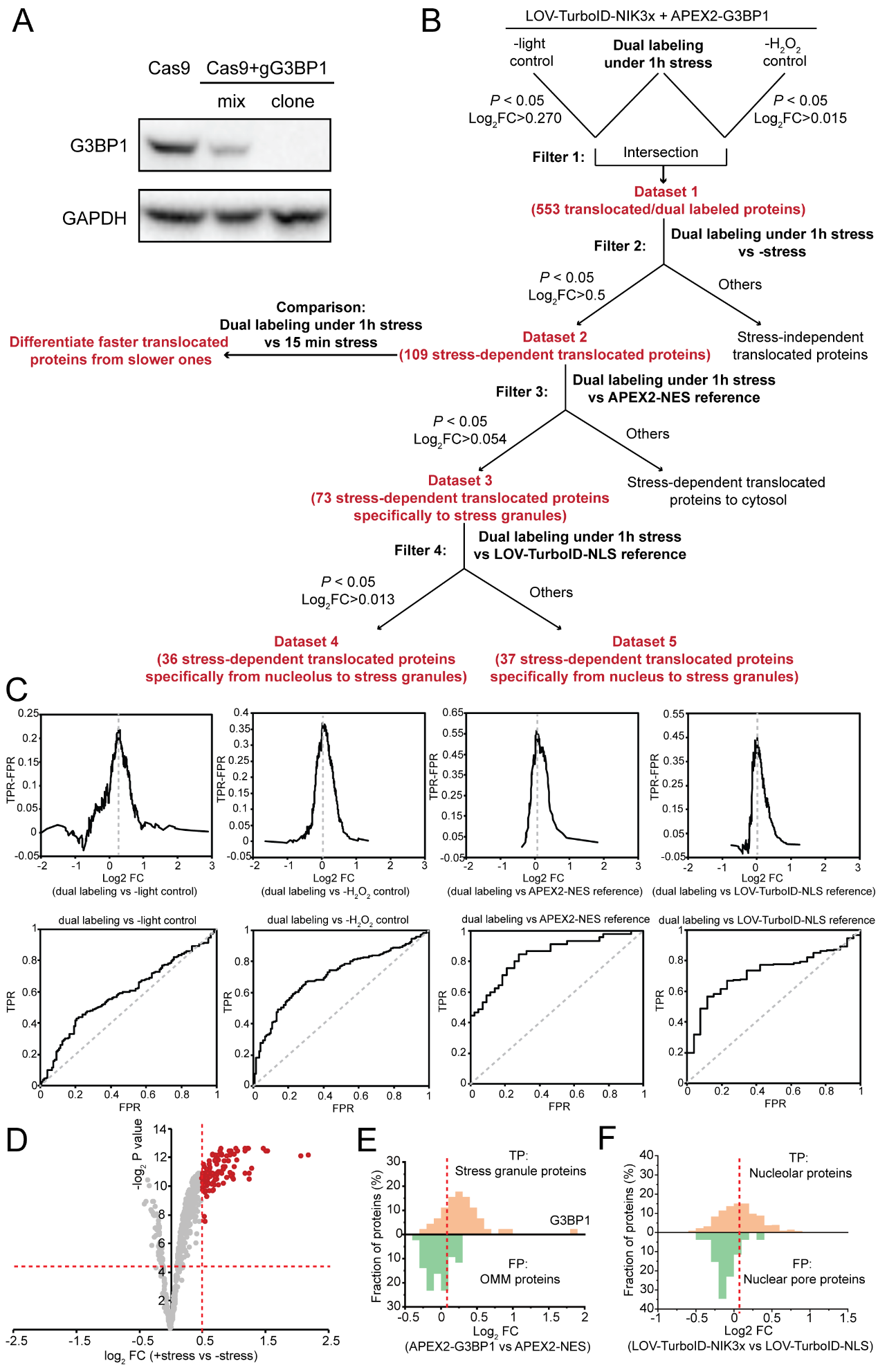

**Figure S5. Analysis of proteomic data for the nucleolus-to-stress granule trafficking experiment shown in Figure 5. A.** Western blot validation of G3BP1 knockout in HEK 293T cells. **B.** Chart showing how the mass spectrometry data was filtered to generate 5 separate proteomic datasets (Supplementary Table 3). **C.** Receiver operating characteristic (ROC) curves of TMT ratios used for assignment of translocated proteins. Proteins were ranked in descending order based on TMT ratio. For the comparisons to omit-light and omit-H_2_O_2_ negative controls, true positives were nucleolar proteins, and false positives were mitochondrial matrix proteins. For the comparison to APEX2-NES spatial reference, true positives were known stress granule proteins and false positives were OMM proteins. For the comparison against LOV-TurboID-NLS reference, true positives were known nucleolar proteins and false positives were nuclear pore proteins. **D**. Volcano plot showing differential enrichment of nucleolus-to-stress granule proteins under basal versus 1 hour arsenite-treated conditions. **E.** Sample histogram showing how the cutoff for the comparison of dual labeling against APEX2-NES reference was applied. **F.** Sample histogram showing how the cutoff for the comparison of dual labeling against TurboID-NLS reference was applied.

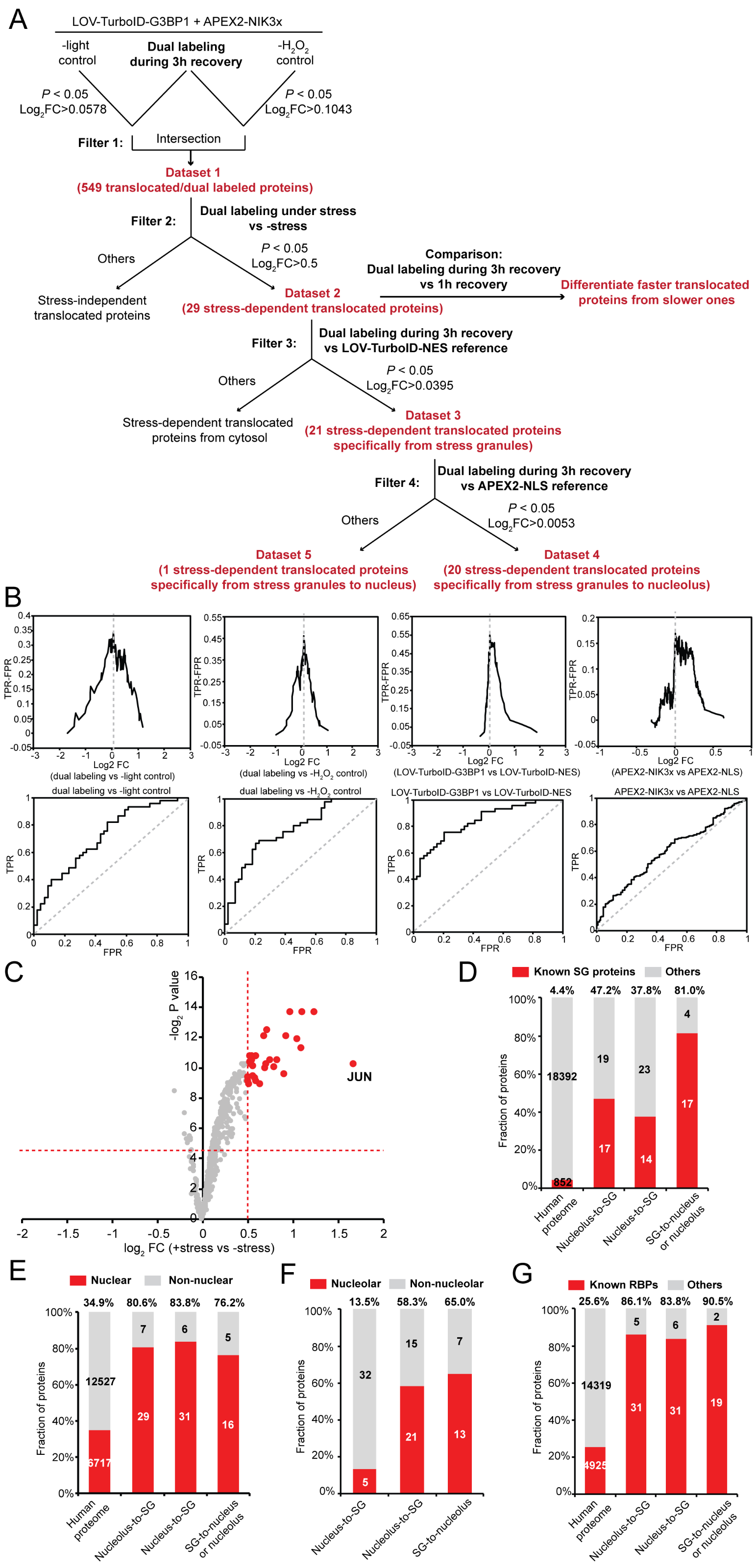

**Figure S6. Analysis of proteomic data for the SG-to-nucleolus trafficking experiment shown in Figure 5. A.** Chart showing how the mass spectrometry data was filtered to generate 5 separate proteomic datasets (Supplementary Table 4). **B.** Receiver operating characteristic (ROC) curves of TMT ratios used for assignment of translocated proteins. Proteins were ranked in descending order based on TMT ratio. For the comparison against omit-light and omit-H_2_O_2_ negative controls, true positives were known stress granule proteins while false positives were mitochondrial matrix proteins. For the comparison against APEX2-NES reference, true positives were known stress granule proteins while false positives were OMM proteins. For the comparison against LOV-TurboID-NLS reference, true positives were known nucleolar proteins and false positives were known nuclear pore proteins. **C.** Volcano plot showing differential enrichment of stress granule-to-nucleolus translocated proteins under basal conditions versus during 3hrs stress recovery. **D-G**. The percentage of known SG proteins (**D**), known nuclear proteins (**E**), known nucleolar proteins (**F**) and known RNA binding proteins (**G**) in each dataset.

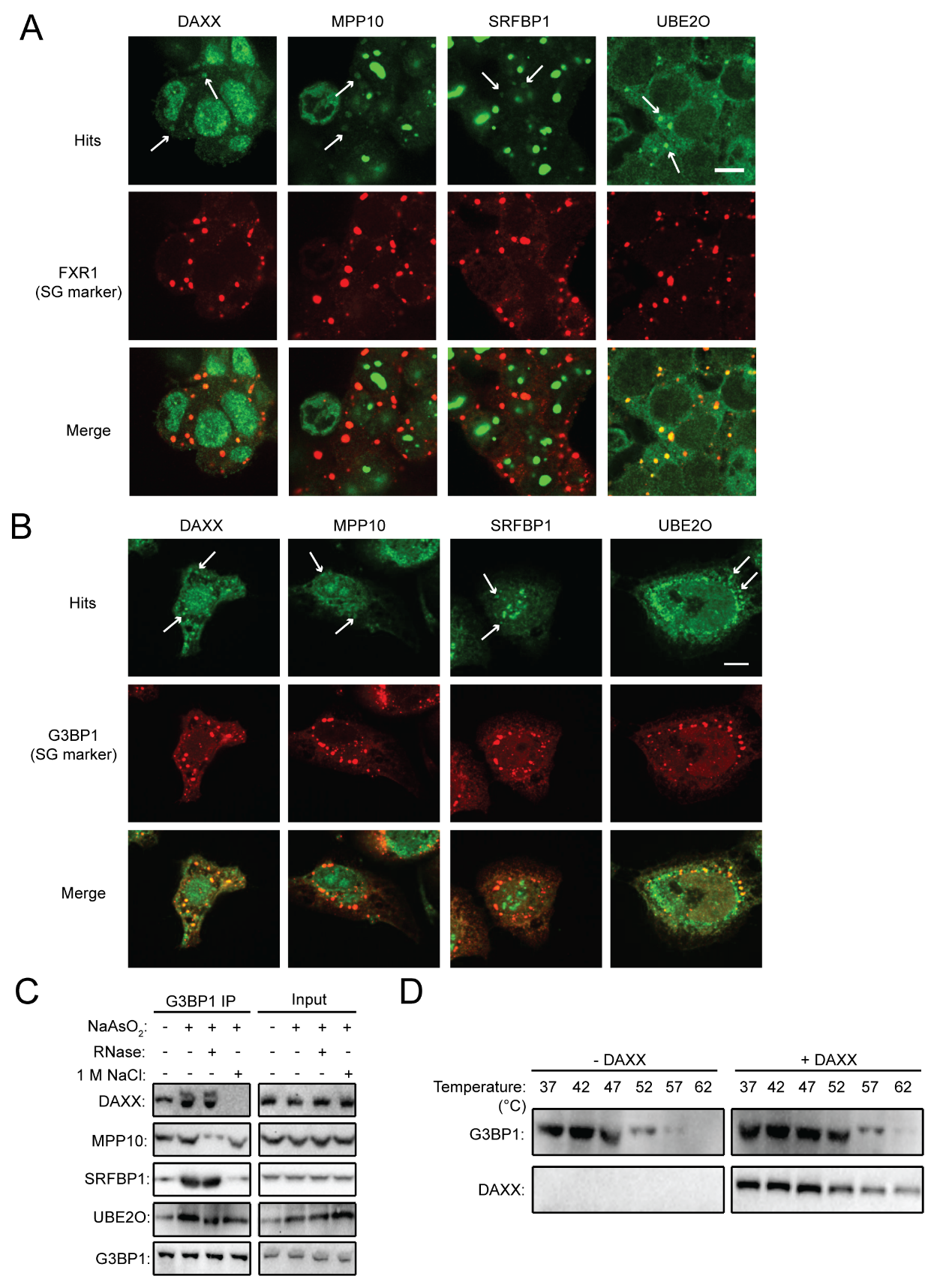

**Figure S7. Follow-up validation for proteins found to traffick between SGs and nucleolus/nucleus. Related to Figure 5. A.** Confocal fluorescence imaging of novel SG proteins identified by TransitID, in HEK 293T cells treated for 1 hour with arsenite. Scale bar, 10 um. **B.** Confocal fluorescence imaging of proteins in (A) in U2OS cells treated with heat (45 ℃) for 2 hours. Scale bar, 10 μm. **C.** Enrichment of novel SG proteins in SGs by anti-G3BP1 immunoprecipitation. IP samples were treated with RNase or high salt to disrupt protein-protein protein-RNA interactions. **D.** Impact of DAXX on the thermal stability of purified G3BP1. Purified G3BP1 were incubated with purified DAXX at 37 ℃ for 30 minutes and treated with different temperature for 3 minutes. After centrifugation to remove aggregates, the remaining soluble fraction was subjected to Western blot analysis.

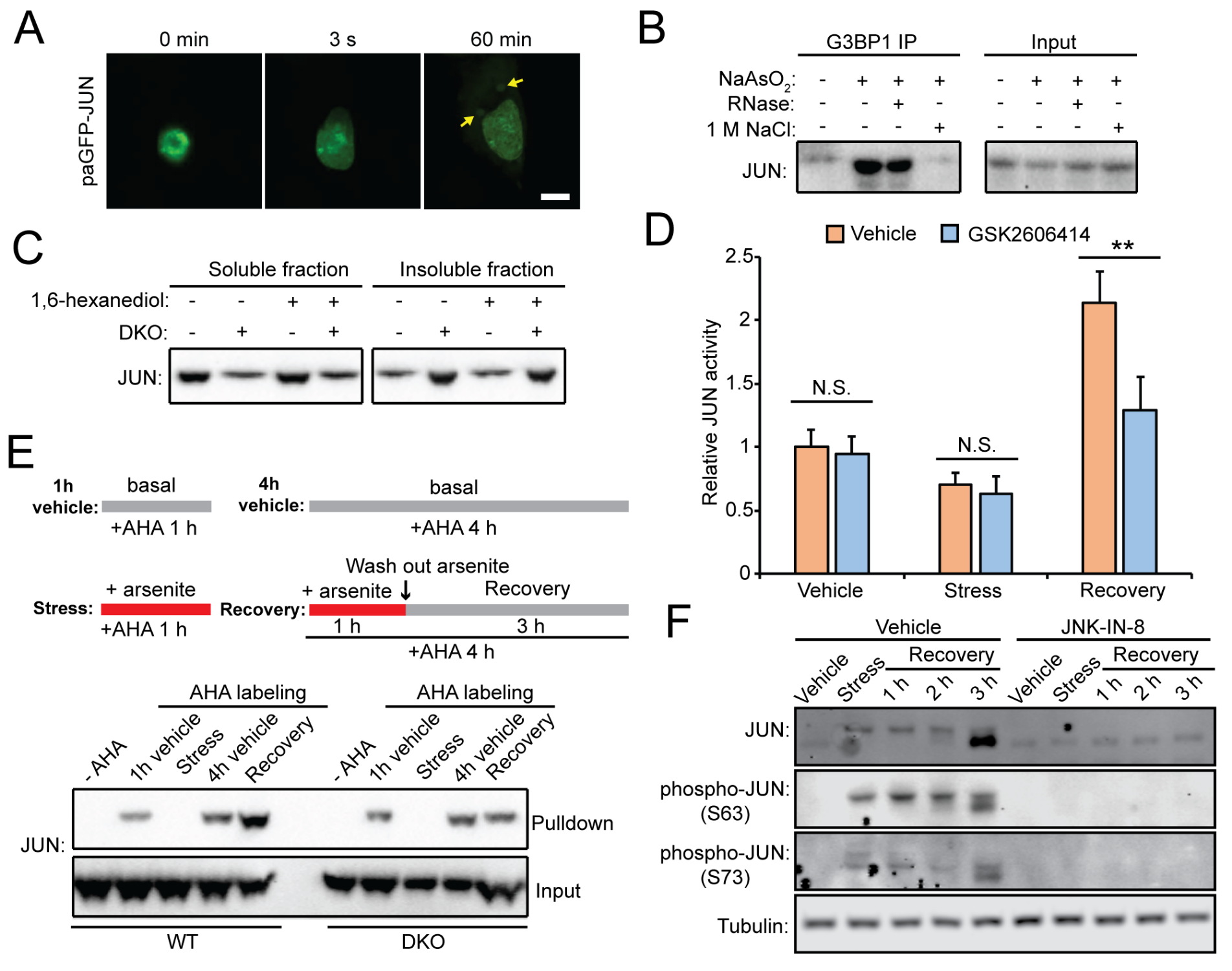

**Figure S8. Stress granules protect JUN from degradation and enable rapid recovery from stress. Related to Figure 6. A.** Imaging of HeLa cells expressing photoactivatable JUN (paGFP-JUN) during stress induction. The nucleus was activated by 405-nm laser immediately after arsenite treatment and GFP signals were monitored with 200-ms intervals. Scale bar, 10 μm. **B.** Detection of JUN-G3BP1 interaction by anti-G3BP1 immunoprecipitation. **C.** Analysis of JUN aggregation in the presence of 1,6-hexanediol. **D.** The impact of stress granule inhibitor GSK2606414 on JUN transcriptional activity under basal, stress and recovery conditions. **E.** Synthesis of JUN under various conditions, determined by AHA labeling. **F.** Total and phosphorylated JUN upon JNK inhibition under various conditions.

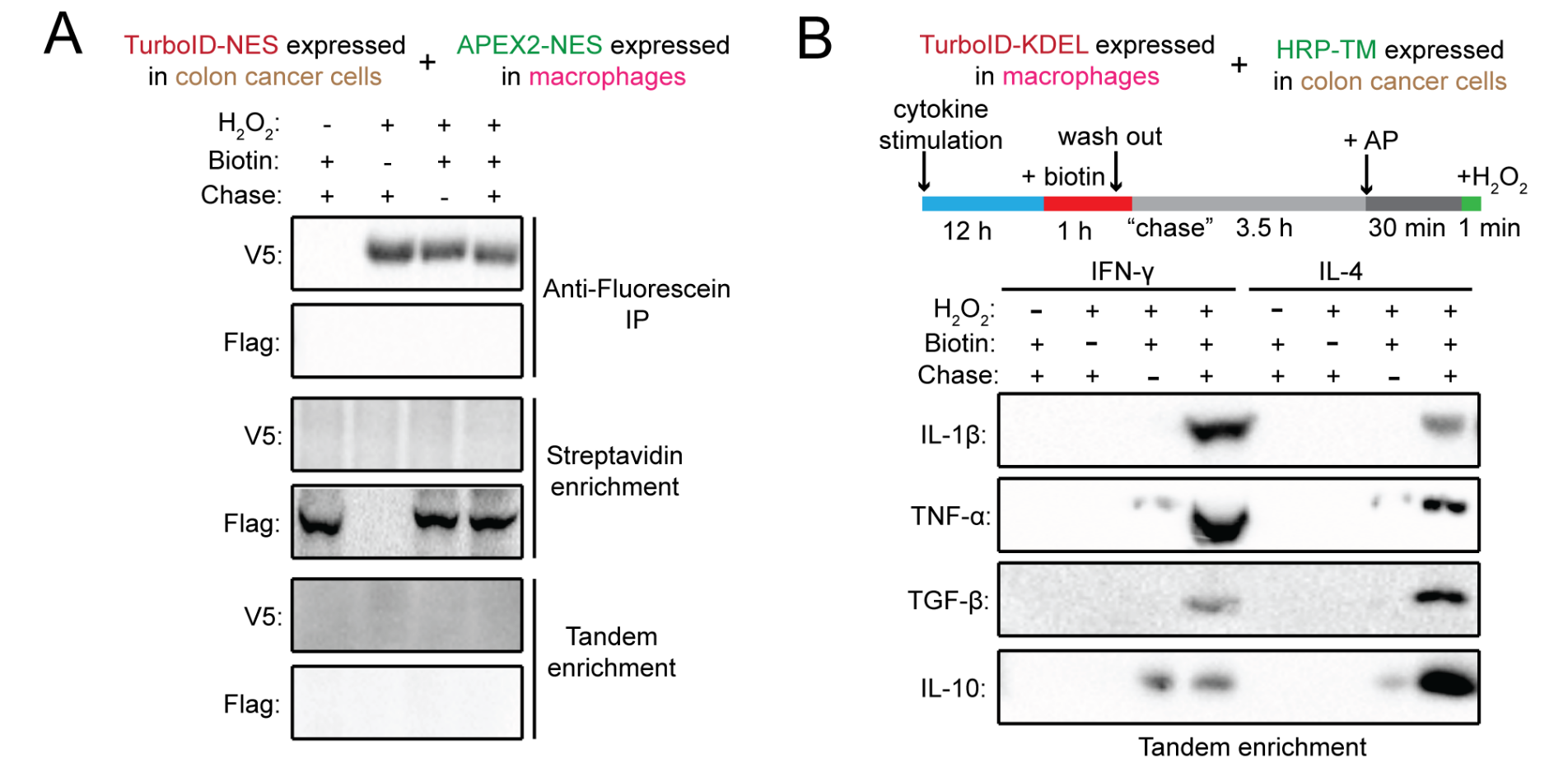

**Figure S9. Detection of intercellular protein communication between cancer cells and macrophages by TransitID. Related to Figure 7. A.** TurboID and APEX2 do not label each other in the co-culture system shown in Figure 7A. **B.** TransitID labeling of proteins that are secreted from macrophages and traffick to the surface of cancer cells following cytokine stimulation in the co-culture system. Western blot detection of specific protein markers after tandem enrichment.

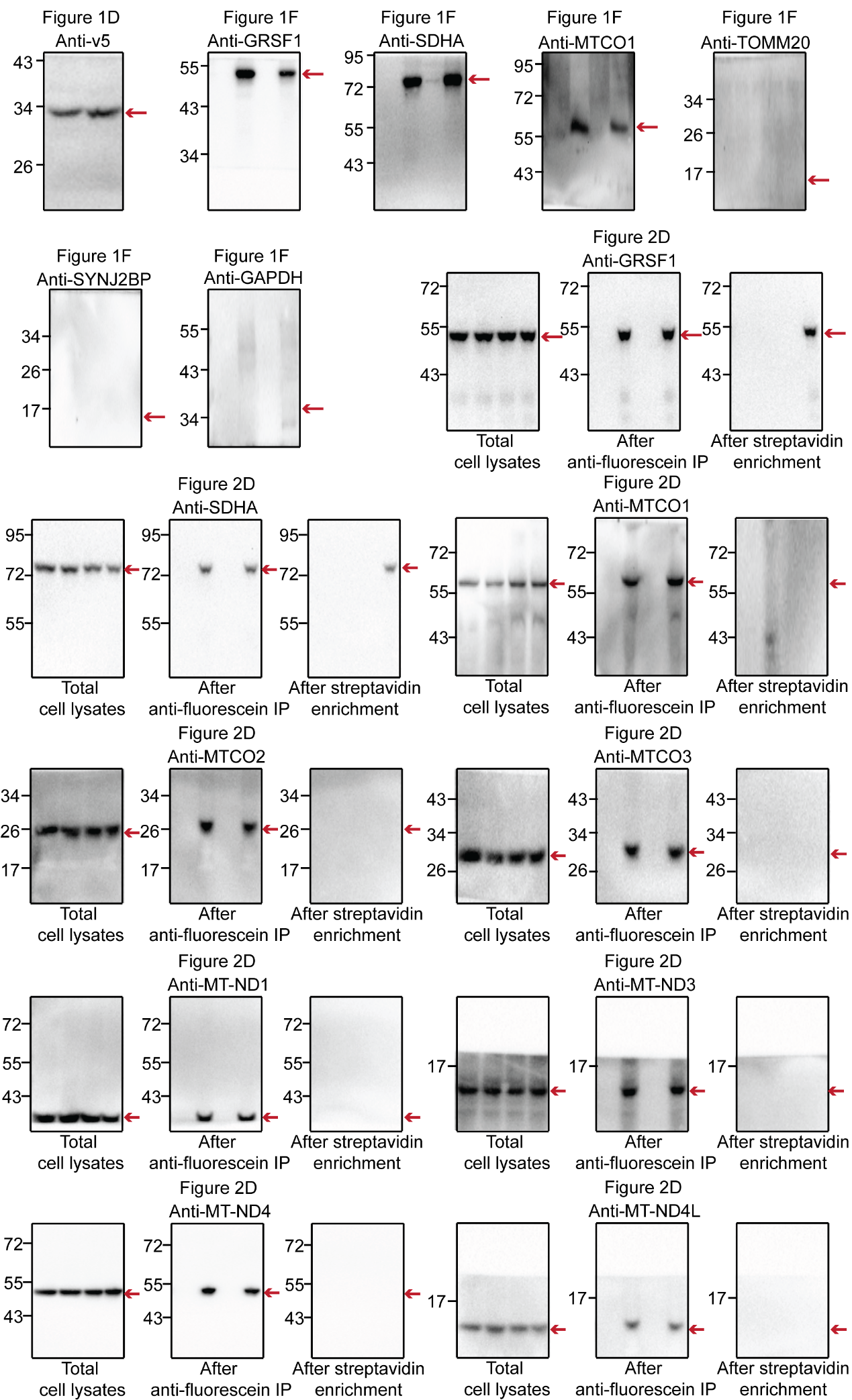

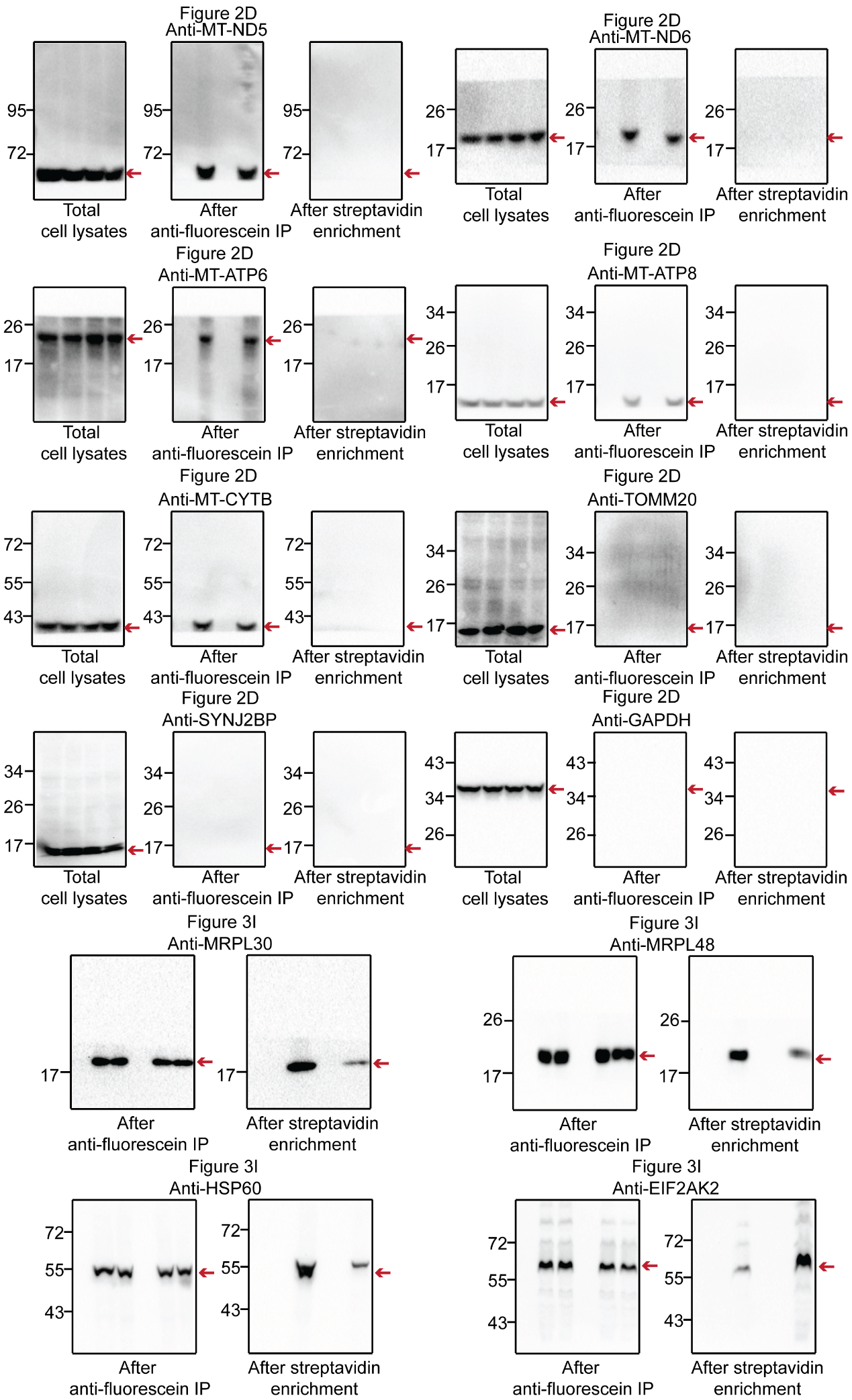

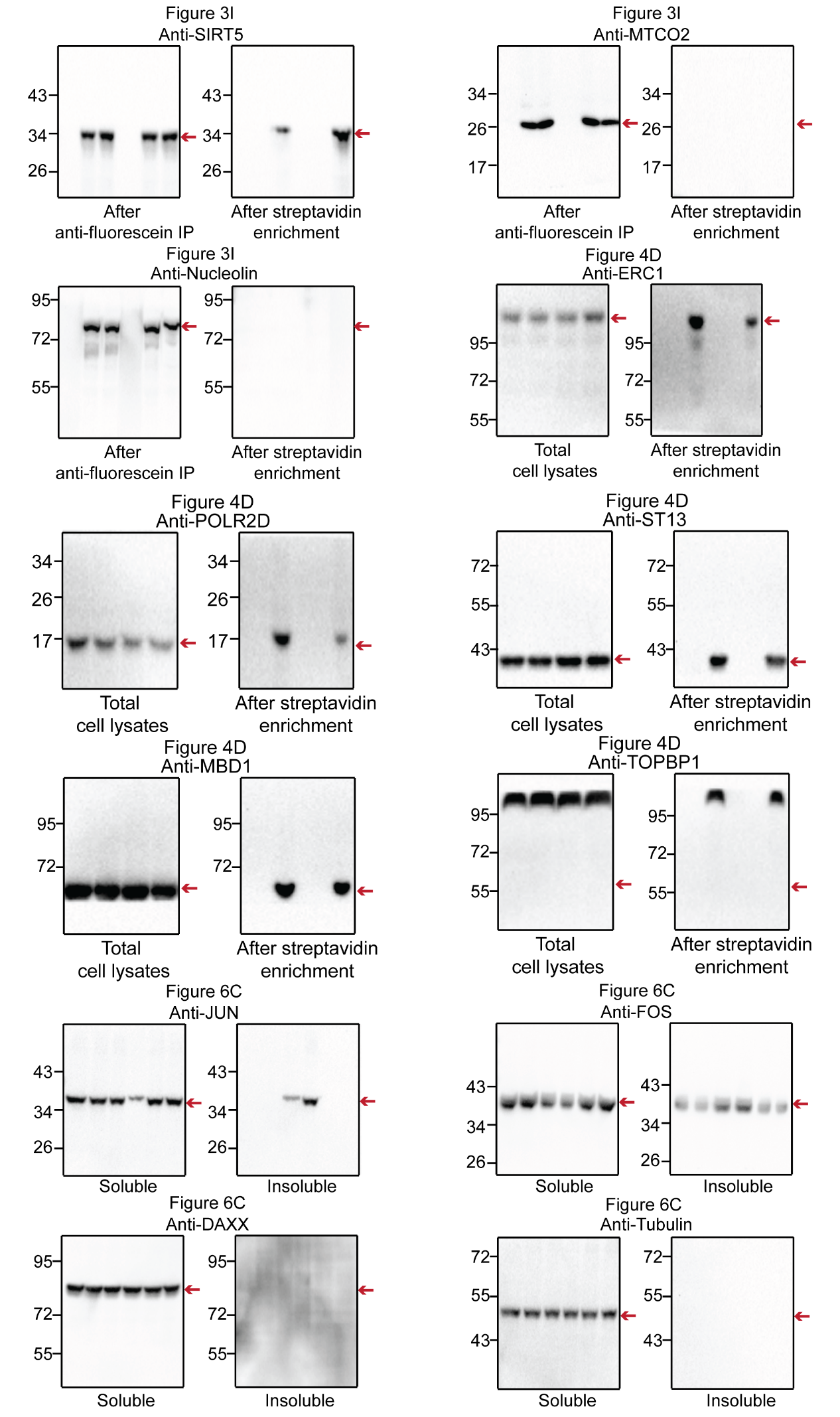

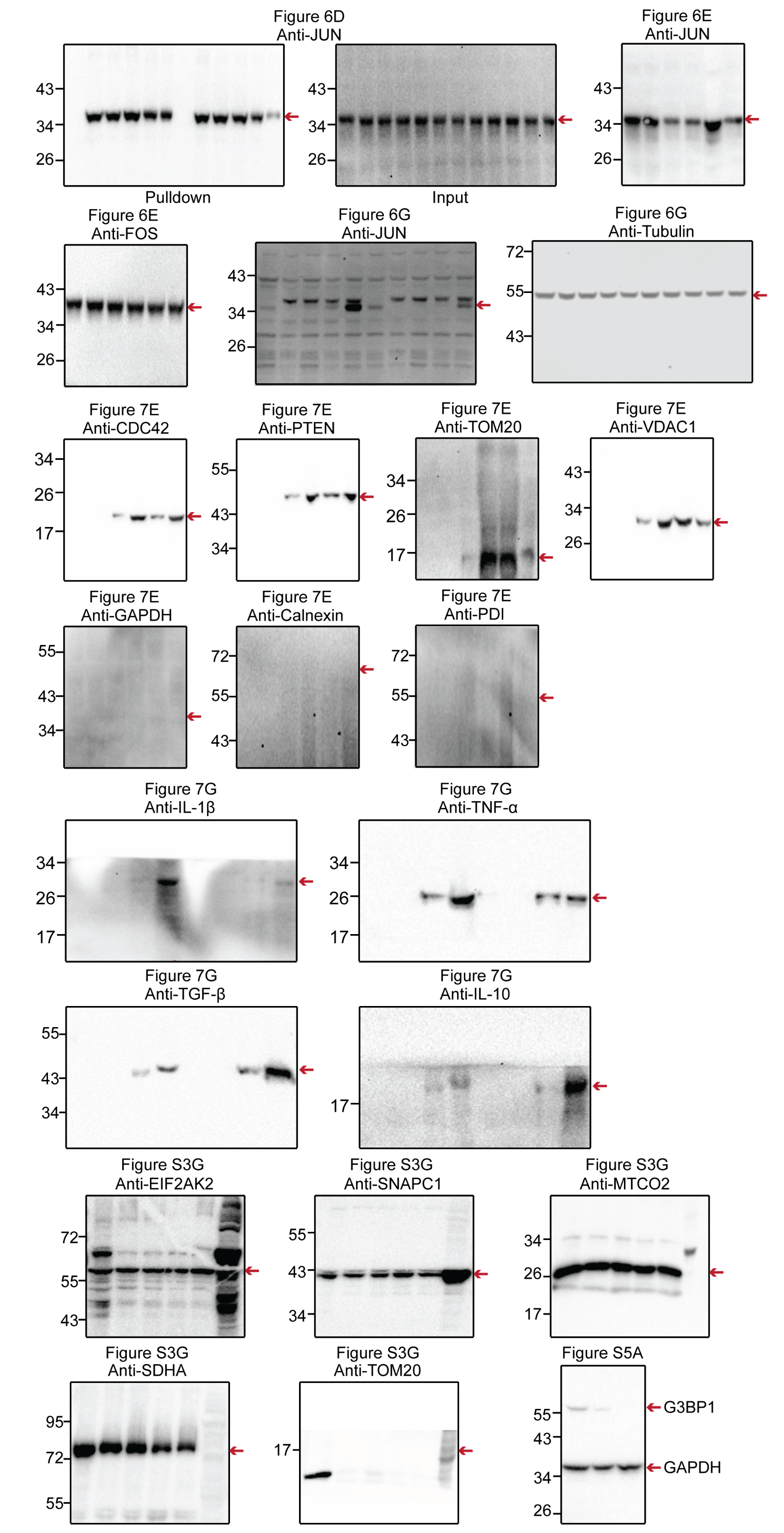

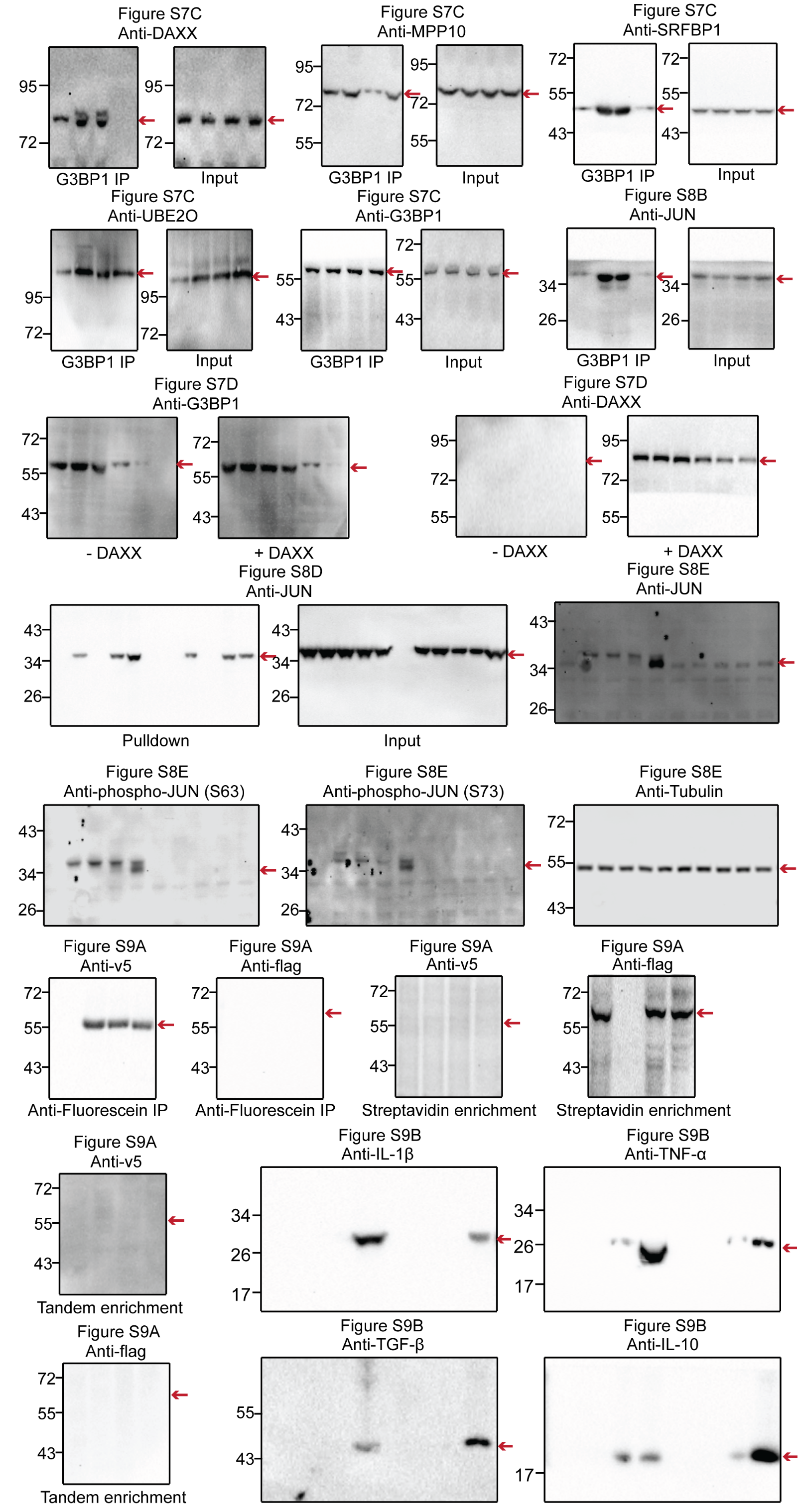

**Figure S10. Uncropped blots used in this study.**

**Supplementary tables**

**Table S1. The list of mitochondrial matrix proteins that originate from the cytosol or OMM.** Related to Figure 3. The fold-change values and p-values for unfiltered proteins are shown in Tab 1. For ROC analysis to determine fold-change cutoffs, true positive proteins (TPs) were GOCC-annotated nuclear-encoded mitochondrial matrix proteins and false-positive proteins (FPs) were GOCC-annotated plasma membrane proteins, which are listed in Tab 2. By filtering with ROC cutoffs and p-values (adj P < 0.05), final lists of OMM-to-mitochondrial matrix and cytosol-to-mitochondrial matrix proteins were generated, which are shown in Tabs 3 and 4, respectively. In Tabs 3 and 4, the translocated proteins are compared to 4 different mitochondrial datasets to determine mitochondrial specificity. The submitochondrial annotation was obtained from MitoCarta 3.0.

**Table S2. The list of proteins that traffick from cytosol to nucleus under arsenite-induced stress.** Related to Figure 4. The fold-change values and p-values for unfiltered proteins are shown in Tab 1. For ROC analysis to determine fold-change cutoffs, true positive proteins (TPs) were “gold+” nuclear proteins from Supplementary Table 3 (Tab 3) of *Branon et al. Nature Biotechnology. 2018* ^7^ and false-positive proteins (FPs) were GOCC-annotated mitochondrial matrix proteins, which are listed in Tab 2. By filtering with ROC cutoffs and p-values (adj P < 0.05), final lists of cytosol-to-nucleus translocated proteins were generated, shown in Tab 3.

**Table S3. The list of proteins that traffick from nucleolus/nucleus to stress granules under arsenite stress.** Related to Figure 5. The fold-change values and p-values for unfiltered proteins are shown in Tab1. For comparison of TransitID labeling against -light or -H_2_O_2_ negative controls, true positive proteins (TP1) were GOCC-annotated nucleolar proteins and false-positives (FP1) were GOCC-annotated mitochondrial matrix proteins. For comparison of TransitID labeling against APEX2-NES spatial reference, true positive proteins (TP2) were GOCC-annotated stress granule proteins and false-positives (FP2) were known OMM proteins assembled in *Hung et al. eLife. 2017* ^8^. For comparison of TransitID labeling against LOV-TurboID-NLS spatial reference, true positive proteins (TP3) were GOCC-annotated nucleolar proteins and false-positives (FP3) were GOCC-annotated nuclear pore proteins. The three sets of TPs and FPs are listed in Tab 2. Three datasets generated during data analysis, including dual labeled proteins, stress-dependent translocated proteins and stress-dependent translocated proteins specifically to SG, are listed in Tab 3. The final nucleolus-to-SG and nucleus-to-SG translocated proteins are listed in Tabs 4 and 5 respectively.

**Table S4. The list of proteins that traffick from stress granules to nucleolus/nucleus during stress recovery.** Related to Figure 5. The fold-change values and p-values for unfiltered proteins are shown in Tab1. For comparison of TransitID labeling against -light or -H_2_O_2_ negative controls, true positive proteins (TP1) were GOCC-annotated stress granule and false-positives (FP1) were GOCC-annotated mitochondrial matrix proteins. For comparison of TransitID labeling against LOV-TurboID-NES spatial reference, true positive proteins (TP2) were GOCC-annotated stress granule proteins and false-positives (FP2) were known OMM proteins assembled in *Hung et al. eLife. 2017* ^8^. For comparison of TransitID labeling against APEX2-NLS spatial reference, true positive proteins (TP3) were GOCC-annotated nucleolar proteins and false-positives (FP3) were GOCC-annotated nuclear envelop proteins. The three sets of TPs and FPs are listed in Tab 2. Three datasets generated during data analysis, including dual labeled proteins, stress-dependent translocated proteins and stress-dependent translocated proteins specifically from SG, are listed in Tab 3. The final SG-to-nucleolus and SG-to-nucleus translocated proteins are listed in Tabs 4 and 5 respectively.

**Supplementary movies**

**Supplementary Movie 1. Photoactivation microscopy of JUN in HeLa cells upon stress induction. Related to Figure S8A. A.** Activation of paGFP-JUN (JUN fused to photoactivateable GFP) in the nucleus of HeLa cells. **B.** Time-lapse imaging of paGFP-JUN following sodium arsenite treatment. paGFP-JUN was activated in the nucleus with 405-nm laser immediately after arsenite treatment and GFP was imaged at 200 millisecond intervals.

**Supplementary Movie 2. Imaging of HeLa cells expressing photoactivatable JUN (paGFP-JUN) and mCherry-PABP during stress recovery, related to Figure 6B. A**. Activation of paGFP-JUN (JUN fused to photoactivateable GFP) in particular stress granules immediately after stress removal. **B.** Time-lapse imaging of paGFP-JUN during stress recovery. The cells were treated with 500 μM arsenite for 1 hour. Immediately after stress removal, paGFP-JUN was activated in particular stress granules by 405-nm laser and GFP was imaged at 200 millisecond intervals.

**Supplementary references**

1.	Rhee, H.-W., Zou, P., Udeshi, N.D., Martell, J.D., Mootha, V.K., Carr, S.A., and Ting, A.Y. (2013). Proteomic mapping of mitochondria in living cells via spatially restricted enzymatic tagging. Science *339*, 1328–1331. 10.1126/science.1230593.

2.	Schäfer, J.A., Bozkurt, S., Michaelis, J.B., Klann, K., and Münch, C. (2022). Global mitochondrial protein import proteomics reveal distinct regulation by translation and translocation machinery. Mol Cell *82*, 435-446.e7. 10.1016/j.molcel.2021.11.004.

3.	Couvillion, M.T., Soto, I.C., Shipkovenska, G., and Churchman, L.S. (2016). Synchronized mitochondrial and cytosolic translation programs. Nature *533*, 499–503. 10.1038/nature18015.

4.	Axten, J.M., Medina, J.R., Feng, Y., Shu, A., Romeril, S.P., Grant, S.W., Li, W.H.H., Heerding, D.A., Minthorn, E., Mencken, T., et al. (2012). Discovery of 7-methyl-5-(1-{[3-(trifluoromethyl)phenyl]acetyl}-2,3-dihydro-1H-indol-5-yl)-7H-pyrrolo[2,3-d]pyrimidin-4-amine (GSK2606414), a potent and selective first-in-class inhibitor of protein kinase R (PKR)-like endoplasmic reticulum kinase (PERK). J Med Chem *55*, 7193–7207. 10.1021/jm300713s.

5.	Dieterich, D.C., Link, A.J., Graumann, J., Tirrell, D.A., and Schuman, E.M. (2006). Selective identification of newly synthesized proteins in mammalian cells using bioorthogonal noncanonical amino acid tagging (BONCAT). Proc Natl Acad Sci U S A *103*, 9482–9487. 10.1073/pnas.0601637103.

6.	McShane, E., Sin, C., Zauber, H., Wells, J.N., Donnelly, N., Wang, X., Hou, J., Chen, W., Storchova, Z., Marsh, J.A., et al. (2016). Kinetic Analysis of Protein Stability Reveals Age-Dependent Degradation. Cell *167*, 803-815.e21. 10.1016/j.cell.2016.09.015.

7.	Branon, T.C., Bosch, J.A., Sanchez, A.D., Udeshi, N.D., Svinkina, T., Carr, S.A., Feldman, J.L., Perrimon, N., and Ting, A.Y. (2018). Efficient proximity labeling in living cells and organisms with TurboID. Nat Biotechnol *36*, 880–887. 10.1038/nbt.4201.

8.	Hung, V., Lam, S.S., Udeshi, N.D., Svinkina, T., Guzman, G., Mootha, V.K., Carr, S.A., and Ting, A.Y. (2017). Proteomic mapping of cytosol-facing outer mitochondrial and ER membranes in living human cells by proximity biotinylation. Elife *6*. 10.7554/eLife.24463.
